# Systematic mapping of orthogonality and domain-swap permissiveness across LysR-type transcriptional biosensors

**DOI:** 10.64898/2026.08.07.743483

**Authors:** Wouter Demeester, Luna Declerck, Marjan De Mey, Brecht De Paepe

**Affiliations:** Centre for Synthetic Biology, Ghent University, Coupure Links c53, S000 Ghent, Belgium

## Abstract

Transcription factor-based biosensors monitor metabolites and control genetic programs, but their wider use is constrained by the limited repertoire of characterized, mutually compatible sensor parts. Here we combine a curated screen of natural LysR-type transcriptional regulators (LTTRs), the largest family of bacterial transcription factors, with systematic domain swapping. Using a standardized construction platform, we convert 17 LTTRs into whole-cell reporters in *Escherichia coli*. Of 16 viable circuits, nine show regulatory activity, including six ligand-inducible biosensors for acetate, benzoate, α-ketoglutarate, chlorohydroquinone, L-homocysteine and salicylate. Mapping interactions across 11 LTTR systems identifies seven mutually orthogonal regulator pairs, providing, to our knowledge, the first orthogonality map for this family. We next construct 108 chimeras across three domain-swap architectures; 69 retain measurable activity, with functional outcomes enriched when the native hinge–ligand-binding-domain association is preserved. As proof of principle, we redesign a cross-reactive regulator: replacing its DNA-binding domain with one from an orthogonal regulator abolishes unwanted promoter crosstalk while preserving ligand-inducible activation of its own target, transferring orthogonality to a previously incompatible pair. Together, natural-diversity screening and domain swapping emerge as complementary routes to expand LTTR biosensor repertoires, revealing a strong link between connector architecture and chimera function.

## Introduction

Transcription factor (TF)-based biosensors couple molecular recognition to gene expression and enable live-cell metabolite sensing, dynamic pathway control and high-throughput strain screening (Fig. 1a) ^1–3^. Despite their broad utility, most engineered systems continue to rely on a small set of well-characterized regulators, such as AraC, LacI and TetR, that are repeatedly repurposed ^1–4^. This limits both the range of detectable ligands and the availability of mutually compatible parts for multi-input circuits. The natural diversity of bacterial TFs offers a large, underused source of alternative sensors spanning diverse ligand classes and regulatory mechanisms ^5–13^.

**Fig. 1.**
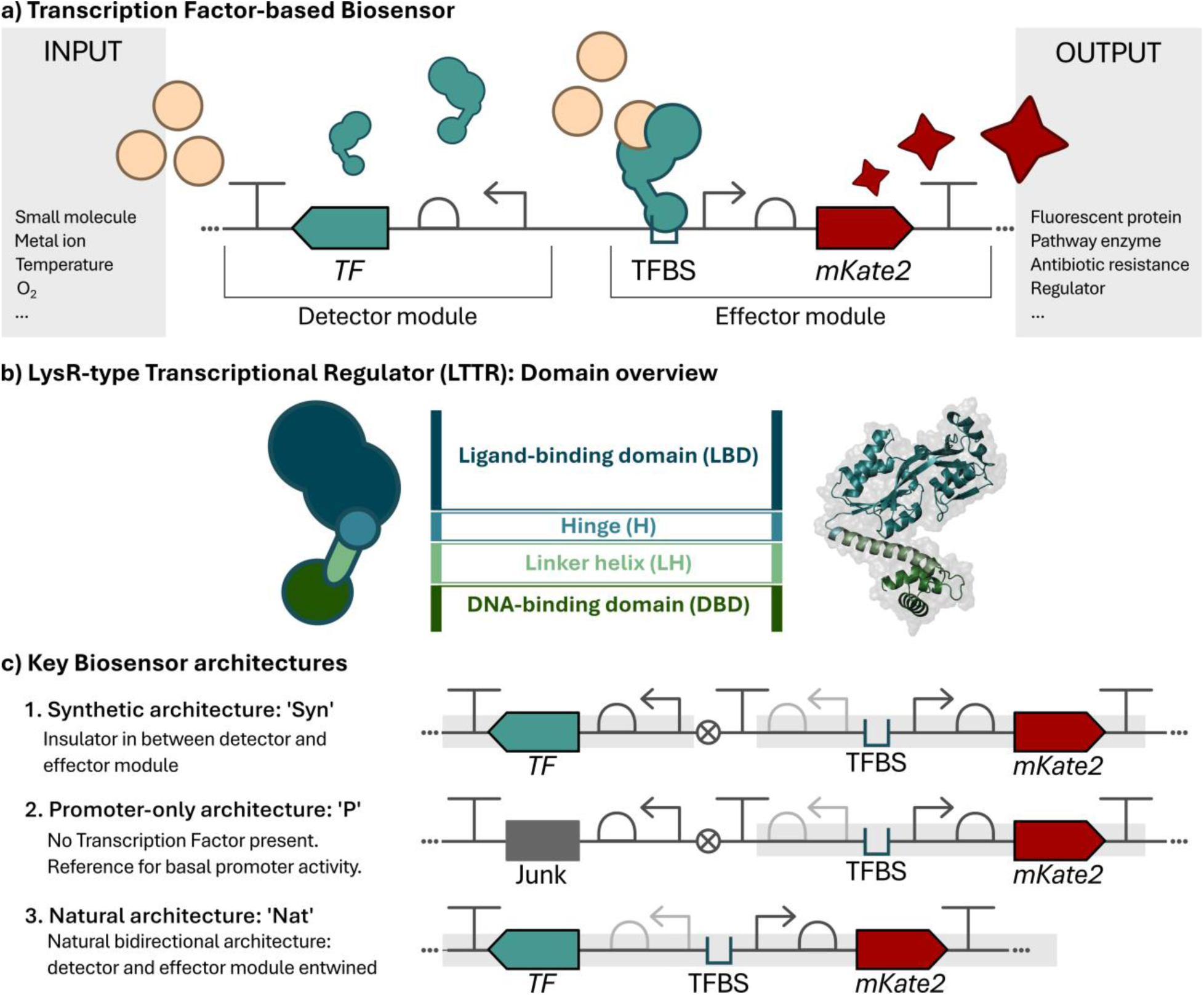
**(a)** General architecture of a transcriptional biosensor circuit. An input signal is detected by a transcription factor (TF), which binds its transcription factor binding site (TFBS) to regulate expression of an output gene (mKate2). **(b)** Domain organization of LysR-type transcriptional regulators (LTTRs), consisting of a DNA-binding domain (DBD), linker helix (LH), hinge (H), and ligand-binding domain (LBD). **(c)** Three biosensor circuit architectures used in this study: a synthetic architecture (Syn) in which detector and effector modules are separated by an insulator sequence, a promoter-only control (P) lacking the transcription factor and containing non-coding (junk) DNA, and a natural architecture (Nat) preserving the native bidirectional promoter organization. TF = transcription factor, TFBS = transcription factor binding site, mKate2 = fluorescent reporter gene, junk = non-coding DNA.

LysR-type transcriptional regulators (LTTRs) are particularly attractive in this context. They constitute the largest family of bacterial TFs ^14–16^ and frequently control metabolic and catabolic pathways through dual-regulatory mechanisms involving basal repression and ligand-dependent activation ^15,17–19^. This behaviour can support low background output and responsive biosensor operation ^15,18,19^. Yet despite the size of the family and the diversity of compounds sensed by its members ^14,15^, LTTRs remain underrepresented in synthetic biology. Their adoption has been limited by structural flexibility and higher-order oligomerization ^20–24^, as well as substantial mechanistic variation among family members ^18,19,25^, which complicates the transfer of knowledge from individual well-studied regulators to the family as a whole.

Transcription factors are also intrinsically modular and are thought to have evolved through the fusion of distinct DNA-binding and ligand-binding proteins into allosterically coupled regulators ^26,27^. Their DNA-binding domains (DBDs) and ligand-binding domains (LBDs) can therefore, in principle, be recombined to generate chimeric regulators with altered sensing or actuation properties. Such domain swapping has been explored across several TF families because it can access protein behaviours that are not readily available from natural diversity alone ^28–32^. In LTTRs, the DBD and LBD are coupled through a linker helix (LH) and hinge (H) region that help transmit allosteric information between sensing and transcriptional control (Fig. 1b) ^26,33^. These connecting elements have been shown to influence DNA or ligand binding, signal transduction, protein folding and oligomerization ^32,34,35^. Studies across LTTRs and other TF families have shown that the connection between recognition modules can strongly influence chimera function ^30–41^. Engineering strategies have therefore focused on this connection, either by adapting native linker regions ^32,34^ or by introducing synthetic sequences ^26,42^. Nevertheless, how the placement and provenance of these connecting elements shape domain-swap outcomes across diverse LTTRs remains poorly understood.

Here we combine natural sensor discovery with systematic domain swapping. Using the previously developed MoBioS construction platform ^43^, we convert a library of natural LTTRs into standardized whole-cell biosensors, compare synthetic and natural architecture circuits (Fig. 1c), and map regulator–promoter orthogonality within the family. We then construct 108 chimeras using three domain boundaries that differ in whether the linker helix and hinge remain associated with the DBD or LBD. Across 17 natural regulators and 11 orthogonality systems, this strategy identifies functional biosensors and mutually orthogonal regulator pairs, reveals an architecture-dependent bias in chimera function toward designs that preserve the native hinge–LBD association, and demonstrates proof-of-principle transfer of orthogonality through replacement of a DNA-binding module. Together, these results position natural-diversity screening and domain swapping as complementary routes to expand LTTR biosensor repertoires, while providing an empirical foundation for mechanistic and predictive investigation of connector compatibility.

## Results

### A standardized LTTR screen yields a broad set of functional biosensors

As a starting point for the development of new biosensors, a library of seventeen LTTRs was assembled by selecting regulators for which the information needed to build a functional biosensor was available, informed by homology to the well-characterized regulator FdeR as reference (Methods; library overview in Table 1) ^32,44^. The library was converted into novel biosensors using the previously developed MoBioS plasmid construction platform ^43^, which enables high-throughput and standardized construction of TF-based biosensors based on the Golden Gate assembly technique ^45^. For each characterized regulator in the library, we compiled a two-page ID sheet documenting its source organism, target promoter, hypothesized ligand and preparation details, biosensor response, and the exact nucleotide sequences of regulator and promoter used for construction; these are collected in Supplementary File 1. Upon integration of the regulator and promoter parts into the platform, “synthetic” biosensor circuits were obtained, referred to as “Syn” followed by the name of the regulator (Fig. 1c). All but one of the biosensor circuits were successfully introduced into the *Escherichia coli* biosensor host; the circuit based on CmpR, an LTTR from *Synechococcus elongatus* responsive to ribulose 1,5-bisphosphate ^46^, was successfully constructed but did not result in viable strains. Two additional circuit architectures were built for each TF. A promoter-only construct (“P”), with junk DNA replacing the regulator, was used to measure basal output, since LTTRs serve as dual-regulators that repress the target promoter in the absence of inducing ligands ^15^. Where the underlying locus architecture allowed, “natural” constructs (“Nat”) were also generated, replicating the native bidirectional promoter context in which the regulator and target gene share an intergenic region (Fig. 1c). “Nat” architectures were built for all regulators except AllS and PqsR, which lack such bidirectional promoter systems.

**Table 1.** LysR-type transcriptional regulators used in this work. For bidirectional promoters, the promoter name reflects both target gene and TF directions. Sequences and promoter annotations are provided in Supplementary File 1.

| Transcription factor | Target promoter | Hypothesized ligand | Original bacterium | Reference |
| --- | --- | --- | --- | --- |
| AllS | *P<sub>_allD</sub>* | Allantoin | *Salmonella typhi* | 47,48 |
| AlsR | *P<sub>_alsS/alsR</sub>* | Acetate | *Bacillus subtilis* | 49,50 |
| AmpR | *P<sub>_ampC/ampR</sub>* | Ampicillin | *Yersinia enterocolitica* | 51,52 |
| BenM | *P<sub>_benA/benM</sub>* | Benzoate, cis,cis-muconate | *Acinetobacter baylyi* | 53,54 |
| CitR | *P<sub>_citA/citR</sub>* | Citrate | *Bacillus subtilis* | 55 |
| CmpR | *P<sub>_cmpA</sub>* | Ribulose-1,5-bisphosphate | *Synechococcus elongatus PCC 6301* | 46 |
| CysB | *P<sub>_cysI/cysB</sub>* | N-acetylserine | *Burkholderia cenocepacia* | 56 |
| GltC | *P<sub>_gltA/gltC</sub>* | α-ketoglutarate | *Bacillus subtilis* | 57,58 |
| LinR | *P<sub>_linE/linR</sub>* | Chlorohydroquinone | *Sphingobium japonicum* | 59,60 |
| LysG | *P<sub>_lysE/lysG</sub>* | L-lysine | *Corynebacterium efficiens* | 61,62 |
| MetR | *P<sub>_metE/metR</sub>* | L-homocysteine | *Salmonella typhimurium* | 63,64 |
| NagR | *P<sub>_nagA/nagR</sub>* | Salicylate | *Ralstonia sp. U2* | 65,66 |
| NahR | *P<sub>_nahR (marionette)</sub>* | Salicylate | *Pseudomonas putida* | 67,68 |
| OccR | *P<sub>_occQ/occR</sub>* | D-octopine | *Agrobacterium tumefaciens* | 69,70 |
| PqsR | *P<sub>_pqsA (truncated)</sub>* | 2-heptyl-4-quinolone | *Pseudomonas aeruginosa* | 71,72 |
| TsaR | *P<sub>_tsaM/tsaR</sub>* | p-toluenesulfonate | *Comamonas testosteroni* | 20,73 |
| TtuA | *P<sub>_ttuB/ttuA</sub>* | L-tartrate | *Agrobacterium vitis* | 74,75 |

The biosensor circuits were subjected to a functionality test *in vivo* by exposing each strain to two ligand concentrations. Due to the unavailability of *N*-acetylserine, the CysB biosensors were not tested with externally added ligand. CitR was tested in the presence of two different candidate molecules, as no ligand has been reported for this regulator. The results of the functional circuits are shown in Fig. 2, with circuits showing no detectable ligand response reported in Supplementary Fig. S1. All strains grew normally in the uninduced state, but upon addition of the respective ligands several strains showed reduced growth (Supplementary Fig. S2): SynMetR strains reached a maximal OD_600_ of 0.2 in the presence of L-homocysteine, while no effect was observed for NatMetR; the CitR “synthetic” sensor was heavily reduced in growth in the presence of either Na-citrate or oxaloacetic acid; and neither SynTtuA nor NatTtuA grew at 20 mM L-tartrate, indicating ligand toxicity. Insolubility of allantoin further interfered with optical density measurements for AllS.

**Fig. 2.**
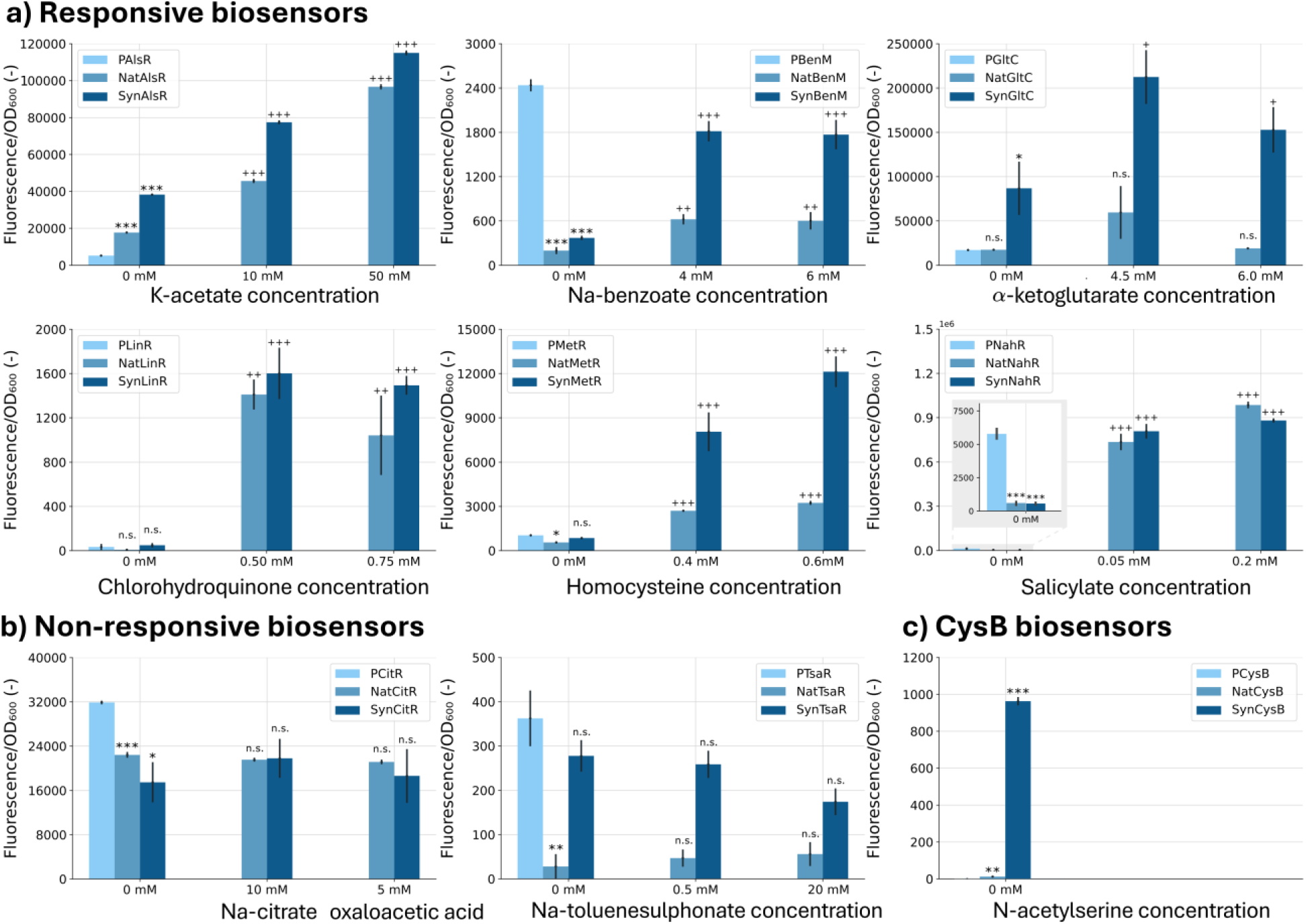
Functional screening of LTTR biosensors in *Escherichia coli*. **(a)** Responsive biosensors. **(b)** Non-responsive biosensors. **(c)** CysB biosensors. For each regulator, the promoter-only control (P) is shown in light blue, the natural architecture (Nat) in blue, and the synthetic architecture (Syn) in dark blue. Two ligand concentrations were tested for each biosensor. For CitR, citrate and oxaloacetate were evaluated because the ligand specificity of this regulator remains uncertain. No induction experiments were performed for CysB because its reported inducer, *N*-acetylserine, was unavailable. Fluorescence was measured after 24 h of growth, corrected for *E. coli* background fluorescence, and normalized to OD₆₀₀. Bars represent the mean of n = 4 biological replicates; error bars indicate the standard error of the mean (SEM). The effect of TF expression (*) was assessed using Welch’s t-tests, whereas the effect of ligand concentration (+) was assessed using one-way ANOVA within each strain. Significance level α = 0.05; */+ *P* < 0.05, **/++ *P* < 0.01, ***/+++ *P* < 0.001; n.s. = not significant. Full test statistics are provided in Supplementary Table S4.

From the library of sixteen viable LTTRs, nine biosensors showed significant up- or downregulation from the basal promoter expression and were therefore considered functional. Among these, the biosensors based on NahR and BenM, as well as NatMetR, showed the most resemblance to the dual-regulatory mechanism characteristic of the LTTR family ^15^. For these systems, as well as LinR and GltC, the difference in biosensor response was already saturated at the lowest tested concentration, indicating that the tested concentrations lie outside the regulators’ response window. Due to the low basal expression of P_linE/linR_, no repression could be discerned for LinR and characterization was restricted to its response to chlorohydroquinone. For regulators GltC, AlsR and CysB, it was not possible to analyse purely repression-based functionality because their ligands are part of the biosensor host’s native metabolism; even in the absence of externally added ligands, a significant upregulation of biosensor output is therefore noted, and addition of the respective ligands further increases the fluorescence output. Specifically, AlsR is responsive to acetate, a central precursor of acetyl-CoA ^76^, and GltC responds to α-ketoglutarate, a central Krebs cycle intermediate ^77^.

In contrast, no activation was noted for the biosensors based on CitR and TsaR (Fig. 2b), but the repression analysis revealed functionality of these regulators *in vivo*, indicating that the lack of induction is related to the added ligand or its transport rather than to circuit failure. TsaR is known to respond to *p*-toluenesulfonate in *Comamonas testosteroni* ^20,73,78^, which can take up the compound using specialized transporters ^79^ that are not present in *E. coli*. For CitR, no inducing ligand has been reported to date, and the two pathway intermediates tested in this work did not produce a measurable response.

Comparison of the “natural” and “synthetic” architectures revealed that, for most regulators, the former resulted in an overall lower biosensor output. The “Nat” circuits recreate the natural architecture of the TF by placing it under control of its native promoter, which alters the response curve relative to the “Syn” counterparts because the regulator is controlled by a different expression system and autoregulation is re-established. MetR provides the clearest example: SynMetR showed strong induction by L-homocysteine but also severely reduced growth, while NatMetR retained the response with no growth defect at the same concentrations. This is most parsimoniously explained by excessive TF synthesis in the synthetic architecture being buffered in the natural architecture by a weaker expression system and negative autoregulation. The same trend is seen for CitR-based biosensors, and to a lesser extent for AlsR, GltC and CysB. Taken together, the study delivers nine functional biosensors covering a diverse set of ligand classes, and shows that circuit architecture itself is an important tuning layer for these regulators.

### Orthogonality within the LTTR family is sparse but present

Functional biosensors were further characterized in terms of orthogonality, a feature that describes the behaviour of such circuits when expressed alongside other biosensors. A pair of TFs is orthogonal when the regulators show no interaction with each other’s target promoters (Fig. 3a). To study this, orthogonality circuits were constructed in MoBioS by combining TF and promoter parts from different regulators, referred to as “PTF_A_SynTF_B_”. As an example, PBenMSynAlsR carries the BenM promoter controlling *mKate2* expression but incorporates the AlsR regulator. Two additional characterized LTTRs, FdeR and ChiR ^32,44,80^, were included to broaden the comparison space, yielding eleven systems and 110 regulator-promoter combinations. Circuits in which CitR was expressed as the transcription factor (SynCitR, 10 combinations) were excluded from the matrix, because CitR expression led to heavily reduced growth (Supplementary Fig. S4). Strains that use P_CitR as the target promoter (in combination with the ten other TFs) were retained. In addition, the two biosensor strains, PBenMSynCysB and PNahRSynCysB, did not reach the stationary phase at the point of analysis and were also disregarded. Lastly, the PFdeRSynLinR strain was not viable, leaving 97 new-to-nature TF-promoter combinations for orthogonality analysis (Fig. 3b).

**Fig. 3.**
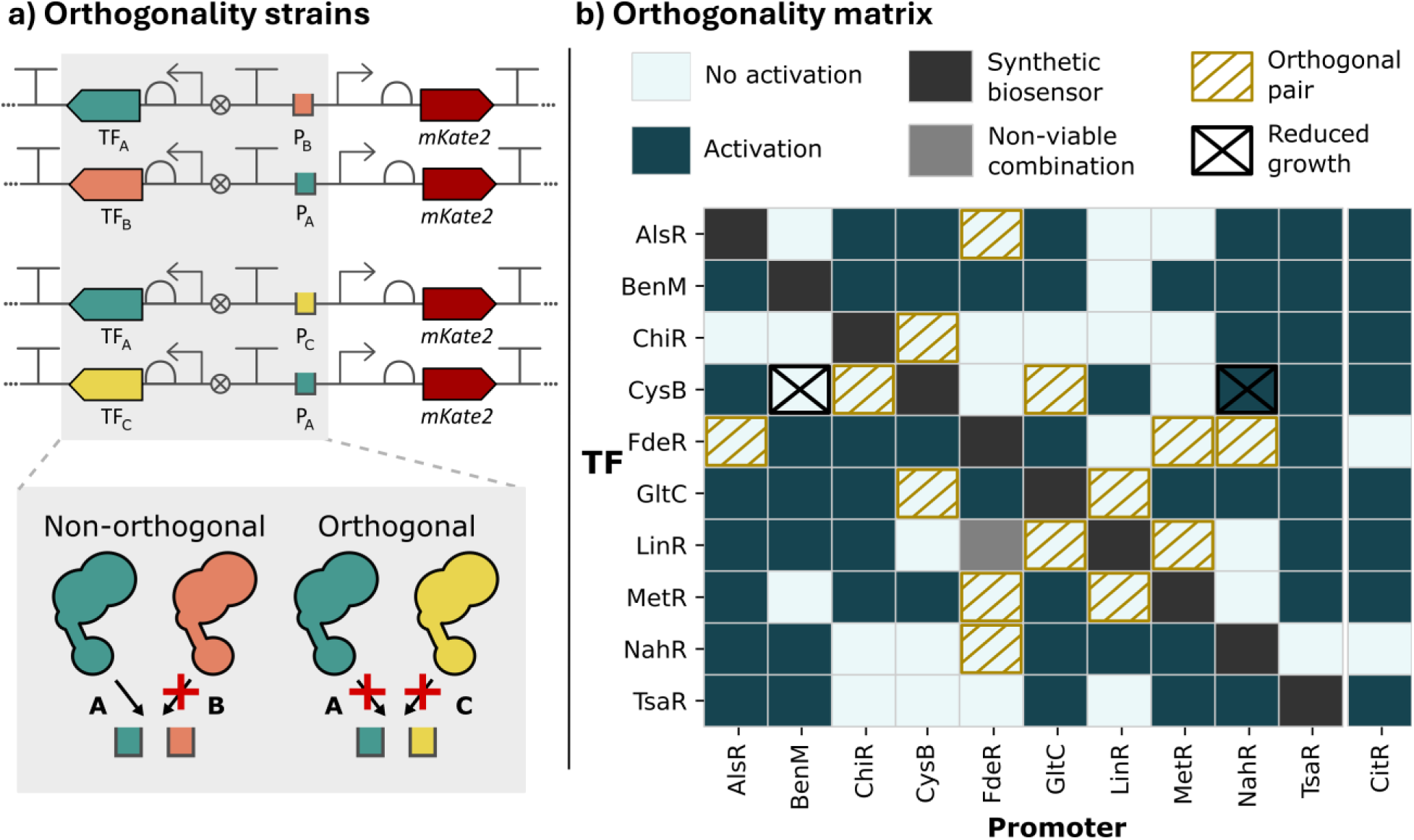
Orthogonality mapping across LTTR biosensors. **(a)** Schematic of the orthogonality assay. Each construct combines the transcription factor (TF) of one biosensor with the promoter of another. Orthogonal regulator-promoter pairs (A–C) show no cross-activation, whereas non-orthogonal pairs (A–B) exhibit promoter activation. **(b)** Matrix summarizing all regulator-promoter combinations tested across the eleven LTTR biosensor systems *in vivo* (based on the per-promoter fluorescence output shown in Supplementary Fig. S3), including the externally characterized regulators FdeR and ChiR. Each cell indicates whether the TF activated the target promoter (dark teal, “Activation”) or not (light teal, “No activation”). Non-viable regulator-promoter combinations are shown in grey, synthetic autologous biosensors in dark grey, and strains exhibiting reduced growth at the endpoint are marked with a cross (Supplementary Fig. S4). The CitR regulator was excluded because of growth inhibition. Members of orthogonal regulator-promoter pairs are highlighted in gold hatching (no activation in both directions).

The fluorescence output of these circuits was measured in the absence and presence of the appropriate ligands. For each TF-promoter combination, the effect of heterologous TF expression (versus the promoter-only control) and the effect of ligand addition (versus the uninduced strain) were tested against the basal promoter output using Welch’s t-tests (full statistics in Supplementary Table S5). A pair of regulators was therefore called orthogonal only when neither of the two heterologous combinations (regulator A’s promoter with regulator B’s TF, and vice versa) showed significance in either test. Within the tested set, 40 out of 97 TF-promoter combinations showed no detectable cross-activity. However, orthogonality is defined in pairs, where both regulators are expected to work in isolation. Among these 40 non-cross-reactive combinations, seven regulator pairs were fully orthogonal (no crosstalk in both directions) and are highlighted in Fig. 3b (gold hatching). These pairs constitute the first orthogonality set reported for LTTR-based biosensors and provide an initial compatibility map for circuit construction within this family.

### Domain swapping across LTTRs reveals a diverse chimera landscape shaped by connector architecture

Building on the characterized biosensor set and the orthogonality map established above, we next asked whether these regulators could be used as starting points for systematic domain swapping, *i.e.* whether the DBD, LBD and connecting linker helix and hinge domains can be combined in new configurations to generate functional chimeric regulators. This approach can both expand the range of orthogonal regulator pairs and enable LBD screening against validated DBD-promoter readouts.

To study domain swapping as an engineering strategy, seven previously characterized LTTRs (AlsR, BenM, ChiR, GltC, LinR, MetR and TsaR) were subjected to the three domain-swapping designs depicted in Fig. 4a. Because TsaR exhibited functionality in the biosensor setup but remained unresponsive to its ligand, TsaR was used only as a DBD donor. In total, 108 chimeras were constructed, of which eight resulted in non-viable circuits upon introduction into the *E. coli* biosensor host. Chimera nomenclature follows the parental source of the swapped domains, in accordance with the MoBioS nomenclature ^43^: for example, a chimera of AlsR’s DBD and BenM’s LBD, H and LH is referred to as SynAlsR_DBD_BenM_LBD_H_LH_ (Fig. 4a).

**Fig. 4.**
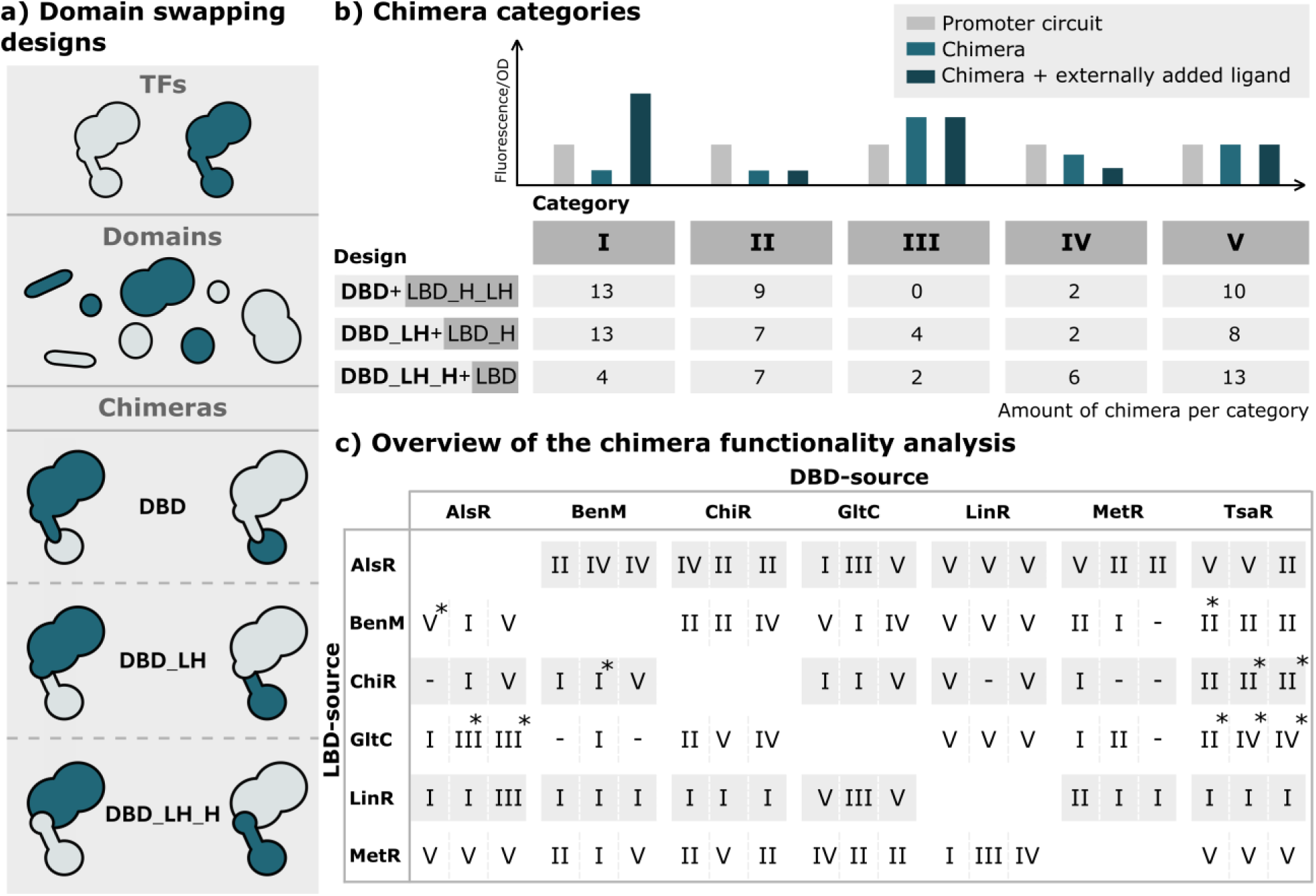
Domain-swapping strategy and functional analysis of LTTR chimeras. **(a)** Schematic overview of the three domain-swapping designs. Chimeras were constructed by exchanging the DNA-binding domain (DBD), the DBD together with the linker helix (DBD_LH), or the DBD together with the linker helix and hinge (DBD_LH_H) between pairs of LTTRs while retaining the ligand-binding domain (LBD) of the recipient regulator. **(b)** Functional classification of the generated chimeras. Category I comprises ligand-inducible chimeras showing induction upon ligand addition, with some also displaying repression of basal promoter activity following chimera expression. Categories II and III comprise chimeras with a significant effect on promoter output but are not ligand-inducible. Category IV contains chimeras exhibiting significant repression upon ligand addition, whereas Category V contains non-functional chimeras showing no significant effect on promoter activity. The table summarizes the number of chimeras assigned to each category for the three domain-swapping designs. **(c)** Overview of the functionality of all generated chimeras, grouped according to donor DNA-binding domain (DBD source) and recipient ligand-binding domain (LBD source). Category assignments (I–V) correspond to the classifications shown in panel B. For each DBD-LBD pairing, category designations are given from left to right: DBD, DBD_LH and DBD_LH_H. Non-viable constructs are indicated by “–”, and strains exhibiting reduced growth are marked with an asterisk (*). DBD = DNA-binding domain, LH = linker helix, H = hinge, LBD = ligand-binding domain.

Chimeras were characterized on both axes of native LTTR behaviour, *i.e.* change in basal promoter output and response to ligand addition, and grouped into five phenotypic categories (Fig. 4b). Category I chimeras showed inducible activation upon ligand addition and represented the most complete (or fully functional) domain-swapping outcome (*e.g.*, SynTsaR_DBD_LinR_LBD_H_LH_). Category II chimeras repressed basal promoter output but did not activate upon ligand addition (*e.g.*, SynTsaR_DBD_LH_BenM_LBD_H_). Category III chimeras showed upregulation in the absence of ligand but no further response to ligand addition (*e.g.*, SynAlsR_DBD_LinR_LBD_H_LH_). Intriguingly, a fourth category was established with chimeras that responded to ligand with decreased rather than increased output (*e.g.*, SynChiR_DBD_LH_H_BenM_LBD_), suggesting inversion of the usual allosteric logic. Category V contained chimeras with no detectable regulatory effect on the target promoter (*e.g.*, SynMetR_DBD_AlsR_LBD_H_LH_). A detailed breakdown of the chimeric biosensor data is shown in Fig. 5.

**Fig. 5.**
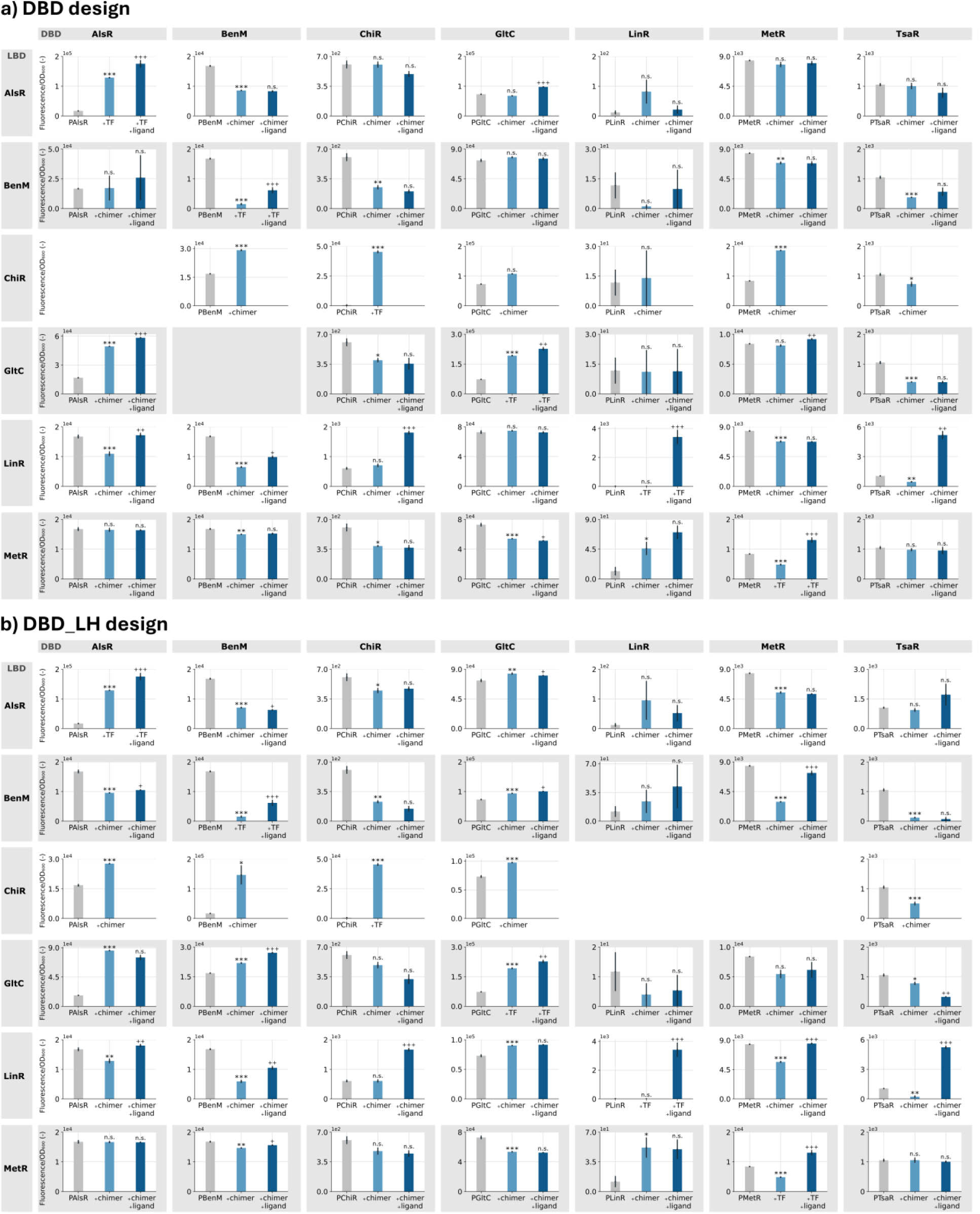
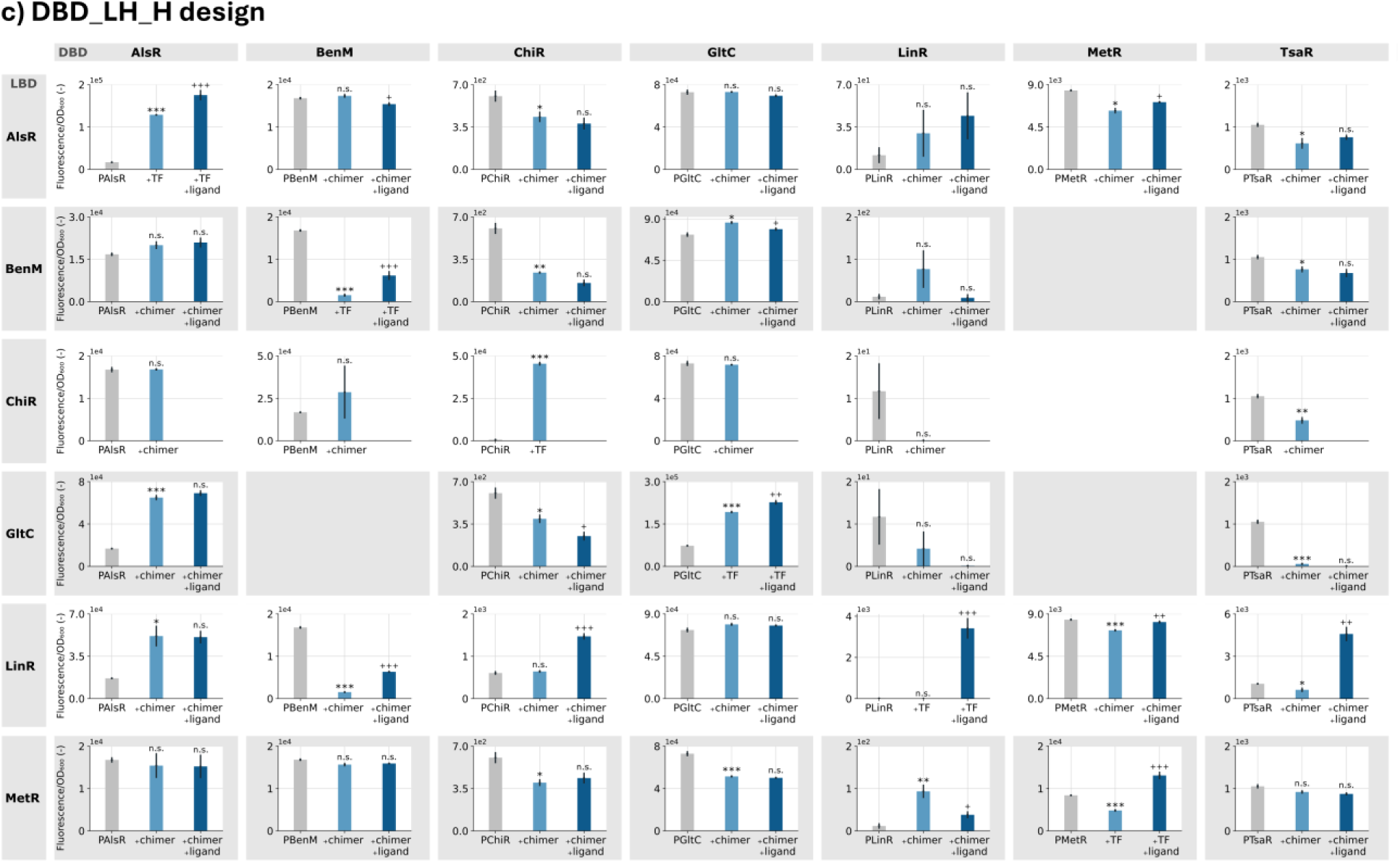
Domain-swapped chimeras rewire promoter output across DBD and LBD donor combinations. **a)–c)**, Domain-swapping designs (Fig. 4a) analysed by DBD donor (columns) and LBD donor (rows). Grey bars show basal output of each promoter circuit (“P” followed by its regulator). Adding a domain-swapped chimera tests its effect on promoter output (light blue), and, where viable, the chimera was assayed with externally added ligand (dark blue; ligand concentrations in Supplementary Table S1). Non-viable domain-swapping strains are left blank. Values were collected after 24 h of growth; fluorescence was corrected for *Escherichia coli* background and normalised to optical density. Bars show the mean of four biological replicates (*n* = 4); error bars, standard error. The effect of expressing a transcription factor or chimera on promoter output (*) and the effect of ligand induction on chimera biosensor output (+) were assessed by Welch’s *t*-tests (α = 0.05): */+ *P* < 0.05, **/++ *P* < 0.01, ***/+++ *P* < 0.001, n.s. *P* > 0.05. Growth curves are in Supplementary Fig. S5 and full statistics in Supplementary Table S6. DBD, DNA-binding domain; LBD, ligand-binding domain; TF, transcription factor; OD₆₀₀, optical density at 600 nm; LH, linker helix; H, hinge.

The central observation of this screen is that LTTR domain swapping opens a broad and functionally diverse landscape of chimeric regulators, whose extent is not obvious from the mechanistic complexity of the family. Thirty chimeras (27.8%) fell into category I, twenty-three (21.3%) into category II, six (5.6%) into category III, and ten (9.3%) into category IV, meaning that 69 of 108 chimeras (∼64%) retained some measurable regulatory activity (categories I–IV; termed *functional* hereafter), whereas the remaining constructs (category V, including the eight non-viable circuits) were *non-functional*. Half of the 36 possible DBD-LBD pairings (18/36) yielded at least one inducible (category I) chimera, and from the remaining, 12 yielded category II or III chimeras only, so 30 of 36 pairings yielded at least one functional (category I–IV) chimera.

The internal structure of this landscape depends strongly on the design choice made when constructing the chimera. Both the DBD-only and DBD+LH designs produced more functional chimeras than the DBD+LH+H design, in which a new hinge was introduced upstream of the LBD. Across the library, the DBD+LH+H design contained the largest fraction of inactive or ligand- repressive chimeras and accounted for a disproportionate share of non-viable constructs: half of the eight non-viable combinations followed this design. Conversely, functional activity was recovered almost exclusively in the two designs that kept the hinge coupled to its original LBD: 29 of the 30 functional pairings yielded their category I–III chimera(s) in a DBD or DBD+LH design, the only exception being the AlsR-LBD/TsaR-DBD pairing with a DBD_LH_H design (Fig. 4b).

This pattern suggests that the hinge is not a trivial boundary that can be moved at will, but rather a key component of functional LTTR allostery. The category II, III and IV phenotypes are consistent with this interpretation. For category II chimeras, the new protein structure may abolish ligand recognition or formation of the induced conformation, resulting in a permanently repressed state, since LTTR activation requires conformational changes in the LBD to propagate to DBD movement. Disrupted inter-domain communication could also inhibit oligomerization, which is essential for LTTR function. For category III chimeras, perturbation of the connecting domains may bias the regulator toward a continuously activated conformation, a phenotype previously noted in studies engineering LBD-DBD connections ^35^. Category IV chimeras suggest that connector perturbation can reverse the direction of regulation altogether, a phenomenon previously observed when altering the length of the hinge domain in the related NodD regulator ^35^. Although further structural work will be required to distinguish lost induction, shifted response windows, constitutive activation, or true reversal of mechanism, the broad distribution of outcomes suggests that the LH and H domains act as active determinants of regulatory logic rather than as neutral connectors ^31,32,34^.

### Rational domain replacement can transfer orthogonality

The orthogonality map from the natural-diversity screen suggested a practical engineering question. If one LTTR pair is non-orthogonal because a regulator recognizes a noncognate promoter, can orthogonality be transferred by replacing the DBD with one from an orthogonal regulator? During the orthogonality study (Fig. 3), BenM was able to bind the target promoter of LinR, P_linE/linR_, resulting in non-orthogonality between the two regulators. In contrast, no such interactions were found between the GltC and LinR systems. Hence, the orthogonality between GltC and LinR can in principle be transferred to the non-orthogonal BenM-LinR-based biosensor pair by swapping the DBD of BenM with the DBD of GltC (Fig. 6a). Two such BenM-GltC chimeras were functional in the domain-swapping experiment, namely SynGltC_DBD_LH_BenM_LBD_H_ and SynGltC_DBD_LH_H_BenM_LBD_. As the latter belonged to category IV (the mechanism of which is unresolved), the orthogonality transfer experiment focused on SynGltC_DBD_LH_BenM_LBD_H_ (Fig.4 and 5).

**Fig. 6.**
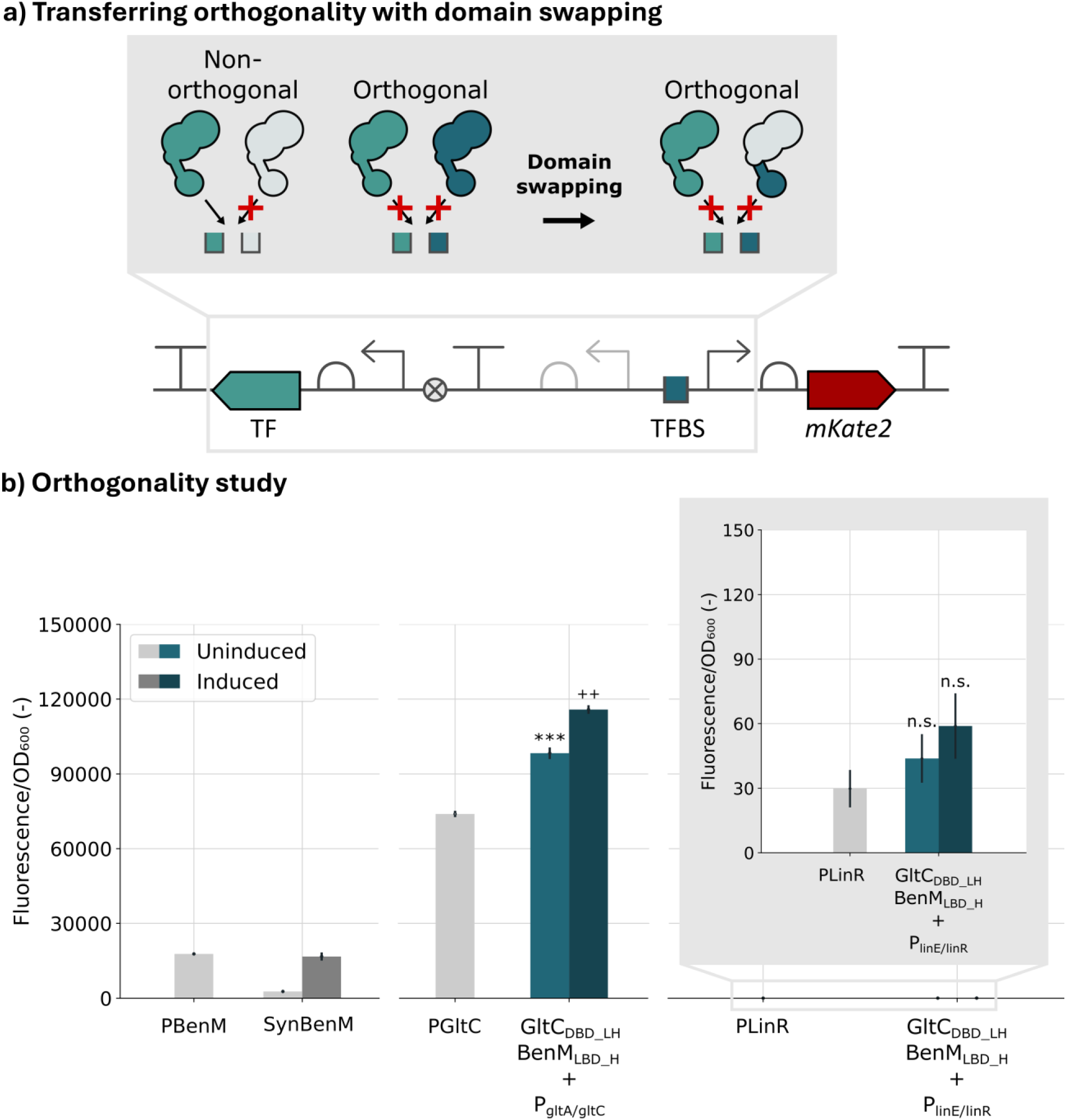
Rational transfer of orthogonality by domain swapping. **(a)** Concept: swapping in the DBD of a regulator that is orthogonal to LinR (GltC) into a regulator that is not (BenM) is hypothesized to transfer the orthogonal phenotype to the chimera. **(b)** The chimera SynGltC_DBD_LH_BenM_LBD_H_ retains ligand-responsive activity with the cognate GltC promoter but shows no interaction with the LinR promoter, consistent with transfer of orthogonality. Reference strains in grey; grey box shows a close-up of the chimera with P_linE/linR_. Genetic circuit parts follow SBOL conventions.

As a proof-of-principle test of this design, this SynGltC_DBD_LH_BenM_LBD_H_ chimera was expressed in the presence of P_linE/linR_ to test functional recognition and inducibility of the chimeric TF with this promoter region. Fig. 6b shows the outcome, clearly indicating that there is no interaction between the chimera and the LinR target promoter. Ligand induction of the chimera with the GltC promoter resulted in a significant upregulation of the biosensor output (Welch’s t-test: t = −6.06, p = 0.0012 < 0.05), consistent with the category I classification of this chimera in the DBD+LH design (Fig. 4). When the target promoter was exchanged with P_linE/linR_, the chimera-based biosensor showed no significant upregulation upon ligand addition (Welch’s t-test: t = −0.58, p = 0.57 > 0.05). Therefore, the domain-swapped chimera maintained orthogonality towards LinR. While based on a single chimera, this proof of principle demonstrates the feasibility of using the orthogonality map to guide rational chimera construction, and shows that domain swapping can be used not only to alter what a regulator senses but also to determine where it can be safely deployed inside larger genetic circuits.

## Discussion

This study shows that LTTR diversification can be tackled most effectively when natural discovery and synthetic domain swapping are treated as parts of the same engineering program. The natural- diversity screen delivered immediate value by converting a broad set of underused regulators into standardized biosensor circuits. Without extensive protein engineering, a substantial fraction of the tested LTTRs already functioned in a heterologous host, demonstrating that the family contains a large reservoir of sensor parts that can be mobilized more quickly than is often assumed. The biosensors recovered cover diverse ligand classes, spanning amino acid metabolism (MetR, CysB), aromatic compound degradation (BenM, NahR, LinR, TsaR), central metabolism (GltC, AlsR) and citrate metabolism (CitR). Several map directly to applied biosensor needs: LinR-based circuits are candidates for detecting lindane in contaminated water ^81,82^; MetR is a candidate sensor for homocysteine, a marker associated with cardiovascular and Alzheimer’s disease ^83,84^; NahR and BenM target salicylate and benzoate, both of industrial interest for the sustainable production of aspirin precursors ^85–87^ and food preservatives ^88–90^; and AlsR and GltC, which sense central-metabolism molecules, could function as readouts of microbial fitness during strain optimization ^91–93^.

At the same time, the orthogonality screen made clear that natural functionality is not sufficient for higher-order circuit design. Compatibility is limited, and identifying mutually usable parts requires explicit testing. Orthogonality in LTTRs is intrinsically harder to define than in repressor-only families because both repression and activation must be tracked simultaneously. At this stage, a case-by-case interpretation combining orthogonality matrices, fluorescence outputs and growth behaviour provides the most defensible workflow. Yet the value of such maps is not only navigational: by resolving how sequence variation shapes compatibility, they lay the groundwork for generating orthogonal hinges de novo, shifting orthogonality mapping from a discovery bottleneck to a lightweight validation step. Seven mutually orthogonal LTTR pairs were identified here, establishing the first orthogonality set for this family and a starting point for that predictive design.

The domain-swapping experiments extend this picture by revealing a broad and structured chimera landscape whose internal organization depends on connector architecture. More than half of the chimeras retained measurable regulatory activity, showing that LTTRs are sufficiently modular to support large-scale domain swapping. Yet they are not modular in a simplistic sense. The results consistently point to the connector region, especially the hinge and its relationship to the LBD, as an important contributor to whether a swapped regulator remains usable. This is consistent with the observation that some chimeras preserved inducibility, whereas others became constitutively biased, permanently repressed, or inverted their input–output response. Together, these outcomes suggest that the exchanged domains altered not only recognition properties but also the propagation of conformational information across the protein. Beyond LTTRs, engineering strategies for transcription factors often assume that DNA and ligand recognition can be swapped relatively independently ^26,28,42^, but our data support a more constrained framework in which successful domain swapping is favoured when functional allosteric coupling is preserved.

The orthogonality-transfer experiment is particularly informative because it connects the natural and synthetic parts of the study into one coherent design strategy. The natural-diversity screen identified a scarce but real set of orthogonal LTTR pairs. The swapping study then showed that one of these orthogonal DNA-recognition modules could be transplanted into a different ligand-response background to repair a cross-reactive system. These results suggest a tractable route toward circuit- compatible LTTR design, in which orthogonal DBDs identified from the natural-diversity map can be paired with desired ligand-response backgrounds using architectures that best preserve allostery. Broader application of the strategy will require additional cases, but a framework is now in place: orthogonality information from the natural-diversity map provides a direct selection criterion for choosing domains to swap.

The dataset reported here was generated in a single host under a focused set of ligand conditions, and chimera phenotypes were classified from *in vivo* endpoint behaviour. These choices kept the screen tractable at scale (108 chimeras) while allowing direct comparison between natural regulators and chimeras. Incomplete responses for some regulators likely reflect ligand transport, metabolism or response-window mismatches rather than intrinsic biosensor failure; this is consistent with the observation that TsaR and CitR repressed their target promoters despite the absence of detectable induction. Cell-free characterization to remove host–circuit confounders ^26,94–96^, biophysical assays to dissect the category IV phenotype, and dose-response profiling of the most promising biosensors in additional hosts are natural next steps that build directly on the parts and architecture-dependent trend identified here. Even with the present scope, the dataset is sufficient to establish practical architectural preferences for LTTR domain swapping and to motivate mechanistic investigation of the sequence and structural features underlying connector compatibility.

Together, the combined dataset delivers a working foundation for LTTR engineering. Natural diversity provides a rapid path to new biosensor parts and a map of compatibility relationships. Domain swapping expands that map into a much larger synthetic landscape, in which DNA specificity, ligand response and orthogonality can begin to be combined deliberately. The central lesson is that LTTRs are engineerable when their connecting domains are treated as active components of regulation. Resolving the structural rules that govern connector compatibility could transform domain swapping from an empirical search into predictive, bottom-up design of allosteric regulators, establishing a generalizable framework for programmable biosensing across the LysR family and beyond.

## Methods

### Strains and growth conditions

All experiments were carried out in *E. coli* TOP10. Strains were grown in MOPS EZ Rich Defined Medium with kanamycin (50 µg/mL) for plasmid selection. All medium components and chemical inducers were obtained from Merck (Diegem, Belgium) unless otherwise stated. Chemical inducers were incorporated into the medium where appropriate: ampicillin sodium salt (A9518), α-ketoglutarate (75890), chlorohydroquinone (224081), L-homocysteine (69453), *p*-toluenesulfonate sodium salt (152536), allantoin (05670), 2-heptyl-4-quinolone (SML0747), L-(+)-tartaric acid (T109), D-(+)-octopine (Toronto Research Chemicals, O239850), benzoic acid sodium salt (B3420), L-lysine (L5501), potassium acetate (P5708), oxaloacetic acid (O4126), sodium citrate dihydrate (W302600), salicylic acid (105910) and (±)-naringenin (N5893). Ligand concentrations used in the orthogonality study and the domain-swapping experiments are listed in Supplementary Table S1. Salicylic acid and naringenin were dissolved in ethanol (99.8% purity); ethanol was evaporated from the wells before introducing the growth medium. All other inducers were dissolved in Milli-Q water and filter-sterilized (0.2 µm).

### Genetic part mining

Candidate LTTR regulators were selected on the availability of the information required to build a functional biosensor: a target promoter with identifiable TF-binding sites (for regulators co-localized with a regulon gene around a bidirectional promoter ^15^, the intergenic region was taken from the NCBI genome browser ^97^) and a known or candidate ligand. Regulators native to *E. coli* were excluded to limit host–circuit interactions, and a mix of Gram-positive and Gram-negative origins was included. As an additional criterion, the candidate set was informed by a protein-protein BLAST homology search (https://blast.ncbi.nlm.nih.gov/Blast.cgi) ^98^ seeded with FdeR from *Herbaspirillum seropedicae* ^32,44^, a well-characterized LTTR and the first for which domain swapping was demonstrated ^32^, used here as a reference query; hits were clustered by E-value and percent identity. Seventeen regulators were selected (Table 1). Sequences were codon-optimized for *E. coli* and made BsaI-HFv2 and PaqCI restriction-site-free for MoBioS compatibility ^43^.

### Plasmid construction

All biosensors were built in the MoBioS plasmid framework ^43^, which has a pBBR1-MCS2 broad-host origin of replication ^99^ and a kanamycin resistance marker. Synthetic biosensor circuits (“Syn”), promoter-only controls (“P”) and natural-architecture circuits (“Nat”) were assembled by Golden Gate cloning (Fig. 1c) ^45^. Domain-swapping plasmids were built starting from the “Syn” biosensors via primer design within the protein sequence, PCR with PrimeSTAR HS DNA polymerase (Takara), gel purification (innuPREP PCRpure Kit, Analytik Jena), and CPEC assembly ^100^ with Q5 DNA polymerase (New England Biolabs). The case-study chimera SynGltC_DBD_LH_BenM_LBD_H_ was constructed in the same workflow. All plasmids were sequence-verified by Macrogen (Amsterdam, the Netherlands) following purification with the QIAprep Spin Miniprep Kit (Qiagen, Venlo, the Netherlands) and transformed by electroporation. Full primer and plasmid and strain lists are provided in Supplementary Tables S2 and S3.

### In vivo fluorescence experiments

Biosensor circuits were analysed in 96-well plate format using a Tecan plate reader (Tecan Group Ltd., Männedorf, Switzerland) at 30 °C with 200 rpm shaking. Optical density at 600 nm and mKate2 fluorescence (excitation 588 nm, emission 633 nm) were measured every 10 min. Number of flashes was set to 25 and the settle time to 150 ms. Assay plates were covered with a Triton X-100-coated lid (Greiner Bio-One, Vilvoorde, Belgium) to prevent condensation. Endpoint analysis was performed after 24 h of growth. All experiments were performed in n = 4 biological replicates.

### Data processing and statistical analysis

Data were analysed in Python v3.8.5 in Jupyter Notebooks ^101^ with the pandas package ^102^. Fluorescence output was normalized per time point to optical density and corrected for medium and host background using a control strain (SynJunk; *E. coli* TOP10 carrying the MoBioS platform with non-functional DNA parts) according to the following normalization equation ^43^:

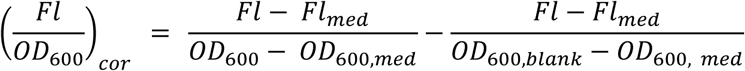

Statistical analyses were performed with the statsmodels ^103^ and scipy ^104^ packages. For the functionality assay, the effect of TF expression on basal promoter output was tested with Welch’s t-tests, and the effect of ligand addition was tested with one-way ANOVA per biosensor strain. For the orthogonality analysis, Welch’s t-tests were used for both the uninduced/promoter comparison and the induced/uninduced comparison; combinations significant in either test were considered non-orthogonal. For the domain swapping and case-study experiments, Welch’s t-tests were used for both repression and induction tests, with significance level α = 0.05.

## Data availability

The data supporting the findings of this study are openly available in the Zenodo repository under DOI [10.5281/zenodo.21838846] (https://doi.org/10.5281/zenodo.21838846), released without restriction under a Creative Commons Attribution 4.0 International (CC-BY-4.0) licence. The repository contains the raw per-well optical density (OD₆₀₀) and mKate2 fluorescence time-series with plate-layout maps for all assays, and the source-data tables underlying Figs 2, 3, 5 and 6. Regulator identity sheets for the characterized LysR-type regulators, compiling source organism, target promoter, hypothesized ligand and its preparation, biosensor response curve, and the exact nucleotide sequences of the regulator and target promoter used for biosensor construction, are provided in Supplementary File 1. Source data underlying Figs 2, 3, 5 and 6 are available in the Zenodo repository.

## Code availability

The custom analysis code used to process the raw data and generate the figures and statistics (Jupyter notebooks and associated Python functions) is openly available in the same Zenodo repository (DOI [10.5281/zenodo.21838846] (https://doi.org/10.5281/zenodo.21838846) under an MIT licence.

## Author contributions

W.D. conceived the study. W.D. and L.D designed and constructed all plasmids and performed all in vivo experiments. W.D. analysed the data. W.D. and B.D.P. drafted the manuscript, with critical revision by L.D. and M.D.M. B.D.P. and M.D.M. supervised the study. All authors read and approved the final version.

## Acknowledgments and funding

W.D. held an FWO PhD grant (FWO 1SC6820N) and L.D. holds an FWO PhD grant (grant no. 1SHCB24N) from the Fonds Wetenschappelijk Onderzoek–Vlaanderen (FWO). B.D.P. holds an FWO postdoctoral fellowship (1246323N). The authors thank the UGent Core Facility “HTS for SynBio” for training, support, and access to the instrument park; the Core Facility is supported by FWO (grants I011118N and I000925N) and BOF (grants BAS018-18, BAS020-131, BOF/BAS/2022/114, and BOF/COR/2022/002).

## Supplementary Information

***<u>Supplementary File 1</u> | Regulator identity sheets*** Two-page identity sheet for each of the 14 characterized LysR-type regulators, compiling the source organism, target promoter, hypothesized ligand and its preparation (supplier, stock concentration, solvent, storage), the biosensor response curve, and the exact nucleotide sequences of the regulator and target promoter used for biosensor construction. Sheets are not included for CmpR (constructed but non-viable in *E. coli*) or for LinR and MetR (biosensor parts reported previously in ^43^). Provided as a standalone PDF and intended as a reusable resource for biosensor part databases.

***<u>Supplementary File 2</u>***

***Supplementary Fig. S1 | Non-responsive LTTR biosensors*** Per-strain endpoint fluorescence and 24 h growth curves for the seven LTTRs that showed no statistically significant induction in the functional screen (AllS, AmpR, LysG, NagR, OccR, PqsR and TtuA). Documents that “non-responsive” is not the same as “non-functional.”

***Supplementary Fig. S2 | Growth curves of the functional biosensors*** 24 h OD₆₀₀ trajectories for the responsive biosensor strains of Fig. 2, confirming that fluorescence differences are not driven by growth defects.

***Supplementary Fig. S3 | Orthogonality fluorescence output*** Corrected, OD-normalised fluorescence of all orthogonality strains, grouped per promoter. These bar plots are the underlying data on which the orthogonality matrix in Fig. 3 is based.

***Supplementary Fig. S4 | Growth curves of the orthogonality strains*** 24 h OD₆₀₀ trajectories for the orthogonality strains, grouped per regulator. Supports the reduced-growth annotations in Fig. 3 and the exclusion of CitR-containing strains.

***Supplementary Fig. S5 | Per-design domain-swap growth curves*** Full 24 h OD₆₀₀ trajectories for the three domain-swapping designs (DBD; DBD+LH; DBD+LH+H), grouped by chimera category.

***Supplementary Table S1. | Ligand concentrations used across all assays*** Inducer concentrations used in the orthogonality and domain-swapping induction experiments; functionality-assay ligand preparations are given per regulator in Supplementary File 1.

***Supplementary Table S2. | Primer list*** Oligonucleotide primers used to construct the domain-swap plasmids (S2a) and the orthogonality case-study plasmids (S2b).

***Supplementary Table S3. | Plasmid and Strain list*** All plasmids and strains used in this work.

***Supplementary Table S4. | Statistical analysis — functionality assay*** T-/F-values and p-values (Welch’s t-tests for TF-expression effect; one-way ANOVA for ligand induction) for every strain in the functional screen; significant values in bold.

***Supplementary Table S5. | Statistical analysis — orthogonality assay*** Welch’s t-test statistics (TF-expression and ligand-addition tests) for all orthogonality combinations; significant values in bold.

***Supplementary Table S6. | Statistical analysis — domain-swapping analysis*** Welch’s t-test statistics for chimera-expression and ligand-induction effects across the three domain-swap designs; significant values in bold.

