## Supplementary File 1 for "Systematic mapping of orthogonality and domain-swap permissiveness across LysR-type transcriptional biosensors"

### AllS:allantoin

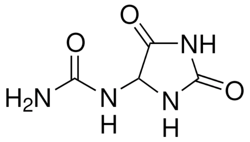


Ligand: Allantoin
Source: Sigma- Aldrich 05670
CAS ID: 97-59-6
Stock: 50 g/L
Solvent: EtOH
Storage: 2-8 °C (2 months)

Response of SynAllS

**No data available**

Metadata

| | |
|---|---|
| Host | *Escherichia coli TOP10 (F- mcrA Δ(mrr-hsdRMS-mcrBC) Φ80lacZΔM15 ΔlacX74 recA1 araD139 Δ(araleu)7697 galU galK rpsL (StrR) endA1 nupG)* |
| Origin | *Salmonella typhi* |
| Parts | AllS: GenBank AE014613.1 |
| | $P_{allD}$: Promoter of *allD* :GenBank AE014613.1 |
| Architecture | Synthetic |
| Experiment | 150 µL EZ Rich medium, 96-well black microtiter plate<br>PerkinElmer Ensight, exc. 588 nm, emm. 633 nm at 30 °C<br>Fluorescence after 24 h of growth |
| Notes | The tested allantoin concentrations interfered with optical density measurements |

#### AllS:allantoin

| | |
|---|---|
| *allS* | GATGATGGCGAGCATTCACCTGCGGTAGCTA**TTAACAGCGTACCGTGTTGAAAATGGATA**<br>PaqCI recognition sequence PaqCI restriction site<br>**AAAAACCGGCTATTTCGGGTTGTCGTTGGGTAAATAATTTCACAATGTCTTCTACCGC**<br>**TTTGCCTCCGCCAAATTTATGCCAGGCGAGGCTTAACGGTGAGGGCGGACGCATCGT**<br>**AGGGATGGTACAGCTCACCAGTTCGTTTTTGTCGATCAGCGTTTGACAAAGCGGCTG**<br>**CGGAACAAAACCGATACCGACGCCTGCCAGATGAGCGGCGATTTTGGTTTCCATATC**<br>**AGGCACGATAATTTCCTTCTGCCCCGGAAGGCGCCAGGCTACCCGCTTGGTCAGCGT**<br>**ACGCGCGCTGTCTTCGATATTAATTGCCGGGAAACGGCGCAACTGCGCTTCTGTTAGA**<br>**GGCCCGCTGACGTGAGCTAAAGGATGATCCGCGGACATCACAAAACGCCATTGTACG**<br>**GAACCTAAAGGGTCGAGCATAAAGGTATTAGCCAACGGCTCGGTGCCAGTTACGCCA**<br>*allS*<br>**ATCGCCAGCGAAAAACCTTCATACAGGAGCGAATCCCAGACGCCCATATAAATTTGC**<br>**CGGGAAAAGTGAAACTGGGTGAAAGGGTAACGCGCATTAAGCCACGATAAGAGACT**<br>**GGCGACCGCTTGCGGGCTATAGAGCAGGTTATTCACCACAATATTGACCTGACGCTCT**<br>**ACGCCGTCGTTAACCTGGCGTAGCTCGTCAGGCATGCTGTCCAGCCACGCAAGCCAG**<br>**TCTTTTGCCTGAGAGAGAAGGTGAGAGCCCGCCGCCGTCAACGATACGCTTCGCGTT**<br>**GTGCGAAAGAACAGCCCTACGCCGGTATTTTCCTCCAGCAGTTTGATGCGGTAGCTA**<br>**GTAGTTGCCGTGGTTTTACACAGGCGCTCCGCCGCTTTGGAGAAACTCCCCGTTTCGG**<br>**CAACGCTGATAAAGGTACGCAGTGTTTCCGGGTCAAACAT**AGAGACCTCCTTGGACCTG<br>BsaI restriction site BsaI recognition sequence GoldenRBS<br>GTACGTCGCTAGCGGGTAGTGTGACGTGGCGCAGGTGATGGACTTCATGCTGAC<br>PaqCI restriction site PaqCI recognition sequence |
| $P_{allD}$ | GATGATGGCGAGCATTCACCTGCGGTACTTACGGTCTCTACAT**GAGATTCCTTCATATCGAG**<br>PaqCI recognition sequence PaqCI restriction site BsaI recognition sequence BsaI restriction site<br>**AATTAACTATTTGCGCTTGCGAATTTTACTGGATATGAGAATTATAAGATATCTGATTT**<br>**AAATAAATCAATATATAAAATTTTATTAATTGATAATTTTATTATGAATGATAAGGAAA**<br>**AAGAACAACTCCTTAATAATTGCACCCATTGCAATTGCACGAGCAGTGTTAATGTGCG**<br>$P_{allD}$<br>**CTTCACGGTATATATCCGGGAGGGTAAATAGAGTGATTAACGTCACATCAAATTTATT**<br>**AACAAAAGAATTCAAACAACGGCTAAAGGATTAGTTATGATTTTCTAAAAAAGCTTTT**<br>**TATTTAAGCAAAAAGATAAAAGCCGCCTGATTATCAAATAAATTCTAATTATATTTTTT**<br>**TGCCTGTCTGGATCACATAATCATTTTATTTTCCCGGTATGTTAATCGCAGTCATGCTTC**<br>PaqCI restriction site PaqCI recognition sequence<br>**ACACCGTCGTTAAAAAGGAAGACAG**ATGGTGGCGCAGGTGATGGACTTCATGCTGAC |

### AlsR:K-acetate

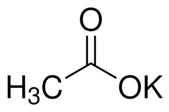


Ligand: K-acetate
Source: Sigma- Aldrich P5708
CAS ID: 127-08-2
Stock: 200 g/L
Solvent: $H_2O$
Storage: 25 °C (2 months)

#### Response NatAlsR and SynAlsR

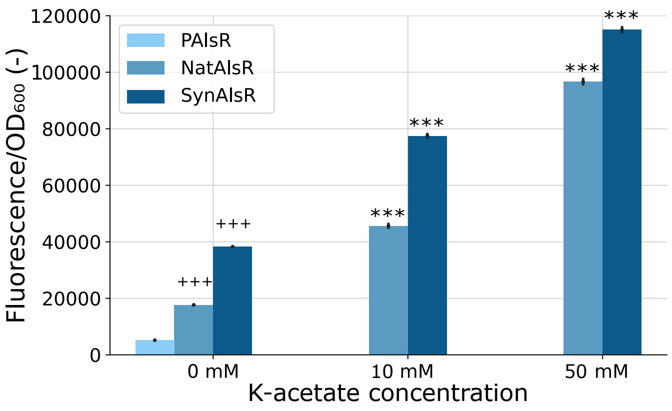


#### Metadata

| | |
|---|---|
| Host | *Escherichia coli TOP10 (F- mcrA Δ(mrr-hsdRMS-mcrBC) Φ80lacZΔM15 ΔlacX74 recA1 araD139 Δ(araleu)7697 galU galK rpsL (StrR) endA1 nupG)* |
| Origin | *Bacillus subtilis* |
| Parts | AlsR: GenBank AL009126.3 |
| | $P_{alsS/alsR}$: Intergenic region between *als*R and *als*S; GenBank AL009126.3 |
| Architecture | Natural and Synthetic |
| Experiment | 150 µL EZ Rich medium, 96-well black microtiter plate<br>PerkinElmer Ensight, exc. 588 nm, emm. 633 nm at 30 °C<br>Fluorescence after 24 h of growth |
| Notes | / |

### AlsR:K-acetate

| | |
|---|---|
| *alsR* | GATGATGGCGAGCATTCACCTGCGGTAGCTA**TCATGTACCTGCATCACTCTCTTTAGTTCT**<br>PaqCI recognition sequence PaqCI restriction site<br>**TGTCTGCTGACAATTTGAAATATGAATAAAATGCTTTATAAGCGGATTATGATTGTCT**<br>**TTGCGGTAAGCGATAACCCATTCTGCATTCAGCTGGATCTGATCCATTTTGCGATAGG**<br>**TCACATCCAGATTAAACAGTTTCTTTGCGCTGCTCGGAACAAAGGTCATACCAATACC**<br>**TGCGCTAACCAGACCAATCACCATTTGATATTCTGTGGCTTCCTGGACAATGTTGGGT**<br>**CTGAAGCCCGCTTGTTCACAGAACTGAATAAAATCCATGTATAGAGTAGGCCATGCTT**<br>**CTTTAGCAACAGTAATAATTGGTTCATCTCTTAAATCCTCAATCGTAATGCTTTCTTTG**<br>**CTGGTCAGCGGATGCTGTTTCGGCAGTGCCAGAACACACGGGCTGCTCTGTGCGGTT**<br>**TCAATATGCAGTGCTGTATGCTGTAAGGGAGGATGAAGTATACCAATATCAATGTTG**<br>*alsR*<br>**CCCTTTAGTAGCTCCTCCTGCTGCCTAGACGAGGATATTTCACGCAGTTCTATTTTCAC**<br>**GCTCGGAAACTTTTTACGGTATTCACGAACAATCGGAGGCAGAAATTCATAGGTTGC**<br>**GCTACCAACAAAACCAATAACCAGCAGACCCTGTTCACCACGTGCGGTACGCTGTGC**<br>**CAGTTCAATACCCTGACCAATCTGCATCAGGGCCATACGACAATGATTCAGAAAAATT**<br>**TCACCGGCTGCGGTCAGTTCAACAAAACGTTTGGTACGTTTCAGCAGGGTAACTCCG**<br>**ACTTCTTCCTCCAGCTGTTTGATCTGCTGGCTGAGAGGAGGCTGCGTCATGTTCAGCC**<br>**GCCGGGCAGCCTTTCCGAAATGAAGCTCTTCGGCTACTGCGATAAAGTATTGAAGAT**<br>**GGCGAAGCTCCAT**AGAGACCTCCTTGGACCTGGTACGTCGCTAGCGGGTAGTGTGACGTGG<br>BsaI restriction site BsaI recognition sequence GoldenRBS PaqCI restriction site<br>CGCAGGTGATGGACTTCATGCTGAC<br>PaqCI recognition sequence |
| $P_{alsS/alsR}$ | GATGATGGCGAGCATTCACCTGCGGTACTTACGGTCTCTCCAT**TCAATATGCATTCCTTTCC**<br>PaqCI recognition sequence PaqCI restriction site BsaI recognition sequence BsaI restriction site<br>**ATAGGTTAATAATTCGTATTACATATTAATCATAAGGCGAATCGATATTGGAGGTCA**<br>**ATTTCCAAAGAGTGTATAGTGAAACTTATCACAAGATATTTAAAATTTTACGTTTAAA**<br>$P_{alsS/alsR}$<br>**ATGCATAATAAGGAGTGAGGGTGTTGACAAAAGCAACAAAAGAACAAAAATCCCTT**<br>**GTGAAAAACAGAGGGGCGGAGCTTGTTGTTGATTGCTTAGTGGAGCAAGGTGTCAC**<br>**AC**ATGGTGGCGCAGGTGATGGACTTCATGCTGAC<br>PaqCI restriction site PaqCI recognition sequence |

#### AmpR:ampicillin

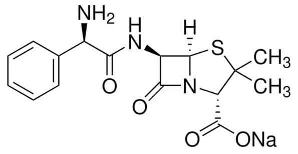


Ligand: ampicillin sodium salt
Source: Sigma- Aldrich A9518
CAS ID: 69-52-3
Stock: 50 µg/ml
Solvent: $H_2O$
Storage: -20 °C (2 months)

Response of NatAmpR and SynAmpR

**No data available**

Metadata

| | |
|---|---|
| Host | *Escherichia coli TOP10 (F- mcrA Δ(mrr-hsdRMS-mcrBC) Φ80lacZΔM15 ΔlacX74 recA1 araD139 Δ(araleu)7697 galU galK rpsL (StrR) endA1 nupG)* |
| Origin | *Yersinia enterocolitica* |
| Parts | AmpR: GenBank X63149.1 |
| | $P_{ampC/ampR}$: Intergenic region between *amp*C and *amp*R*;* GenBank X63149.1 |
| Architecture | Natural and Synthetic |
| Experiment | 150 µL EZ Rich medium, 96-well black microtiter plate<br>PerkinElmer Ensight, exc. 588 nm, emm. 633 nm at 30 °C<br>Fluorescence after 24 h of growth |
| Notes | / |

#### AmpR:ampicillin

| | |
|---|---|
| *ampR* | GATGATGGCGAGCATTCACCTGCGGTAGCTA**TTATTCACCTTCACTTTTCATTTCGCTCAG**<br>PaqCI recognition sequence PaqCI restriction site<br>**CAGCCACAGTGCAAAATCGCGCATTGCCGGTGTTTCGGTACGGCTCTGCAGACGTGT**<br>**CAGCCAATAACCACCTAAGCTAACGGTGGTGCTAAACGGCTGAATAATACGTTCTGA**<br>**ACGCAGCAGATGCATAAACATTGCCGGAGGTGCCAGTGCAATACCAATTTCTGCCTG**<br>**TGCTGCTTCCAGCATGCTAACGCTGCTATCAAACATCATAATCTGCTGGCTCGGACTC**<br>**GGAACCGGACCACCGGCTGCTGCAAACCATGCTGACCATTCATCACGACGATAACTA**<br>**CGCAGCAGCATGAAACGCTGCAGATCGGTCGGATGACGCAGATCTTTTGCCAGGCTC**<br>**GGTGCACACAGCGGTGCCAGCGGAGCATCACACAGAAACTGGCTTTCGGTGCCATGC**<br>**CATGCACCATTACCATAACGAATTGCATAATCCAGACCTTCGGCAACCACATCAACCC**<br>*amp*R<br>**GATTATTATGGGTACTCAGCAGAATATCAACATGCGGACTATGCTGCTGAAAATCAC**<br>**GCAGACGGCTCAGCAGATAACCGGTTGCAAATGCACCAACAACACCAACACGAACTT**<br>**TTTCACGCATAATACCTGCGCTAAAACGATCCAGGGTATGTGCAATACGATCAAAGCT**<br>**ATCATTCAGAATCGGCAGCAGATTTTCACCTTCGGTGGTCAGAACCAGACCACGGCTA**<br>**ATACGAATAAACAGACGACAATTCAGACGCTGTTCCAGTGCTTTAACCTGCTGGCTAA**<br>**TTGCTGCATGGGTAACATTCAGTTCAATTGCTGCTTTGGTAAAGCTCAGCTGACGTGC**<br>**TGCTGCTTCAAATGCACGCAGGCTATTCAGCGGAATATAGCTACGCACCAT**AGAGACC<br>BsaI restriction site BsaI recognition sequence<br>TCCTTGGACCTGGTACGTCGCTAGCGGGTAGTGTGACGTGGCGCAGGTGATGGACTTCATGC<br>GoldenRBS PaqCI restriction site PaqCI recognition sequence<br>TGAC |
| $P_{ampC/ampR}$ | GATGATGGCGAGCATTCACCTGCGGTACTTACGGTCTCTCCAT**AGATTGACTTGTTAGATTT**<br>PaqCI recognition sequence PaqCI restriction site BsaI recognition sequence BsaI restriction site<br>**TCTATTATCAAGTGCTAAATATAATCGATTGTTATCCATAGTCAATCATTGCAGAATTC**<br>$P_{ampC/ampR}$<br>**TTCACGCAAAAGGAGCCACACTGCATGCCATTTATCAGTCTATGGAAGATTTACTA**AT<br>PaqCI restriction site<br>GGTGGCGCAGGTGATGGACTTCATGCTGAC<br>PaqCI recognition sequence |

### BenM:Na-benzoate

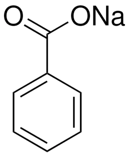


Ligand: Na-benzoate
Source: Sigma- Aldrich B3420
CAS ID: 532-32-1
Stock: 100 g/L
Solvent: $H_2O$
Storage: 25 °C (2 months)

#### Response NatBenM and SynBenM

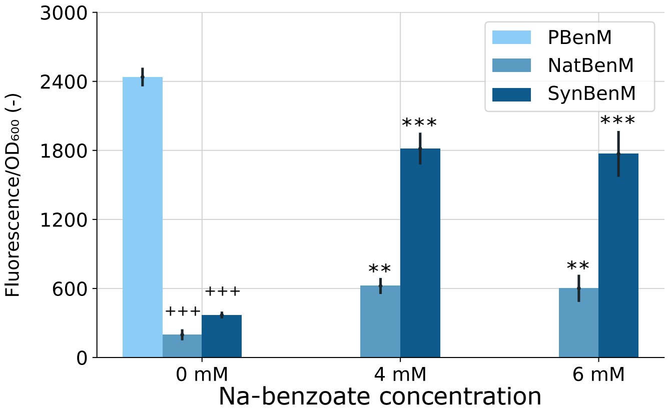


#### Metadata

| | |
|---|---|
| Host | *Escherichia coli TOP10 (F- mcrA Δ(mrr-hsdRMS-mcrBC) Φ80lacZΔM15 ΔlacX74 recA1 araD139 Δ(araleu)7697 galU galK rpsL (StrR) endA1 nupG)* |
| Origin | *Acinetobacter baylyi* |
| Parts | BenM: GenBank AF009224.2 |
| | $P_{benA/benM}$: Intergenic region between *ben*A and *ben*M*;* GenBank AF009224.2 |
| Architecture | Natural and Synthetic |
| Experiment | 150 µL EZ Rich medium, 96-well black microtiter plate<br>PerkinElmer Ensight, exc. 588 nm, emm. 633 nm at 30 °C<br>Fluorescence after 24 h of growth |
| Notes | / |

#### BenM:Na-benzoate

| | |
|---|---|
| *benM* | GATGATGGCGAGCATTCACCTGCGGTAGCTA**TTACCAGTTTGGCGGCTCAGTAAAACCTT**<br>PaqCI recognition sequence PaqCI restriction site<br>**CATAGGCATATATCTGGCGAATCGTTTCGTACAACGAATAAATATAAGTGCTTTCCTC**<br>**CATATTGCGCACGGCAATATAGATGGGCGTGATGGCATCGGGATCTAGAAGCGGCAC**<br>**ATAGCTCAGATTAAAAAGCTGGATACTTTGCGTACTGGCCGGTACCAATGAAATTCCT**<br>**TCACCCGCTGCCACCAATCCAAGCGCCAGTTGAACTTCACGGACTTCATTAATTTTGG**<br>**TGGGTTCTAAGCCATGATCAGAAAAGATATTCATTACATGGGTAGAAAAATTAGGCT**<br>**TAGGCGAGCTTGGATAAAGCAAAATTTTTTCATCAATTAAATCATTTAAGTGCACGCC**<br>**TTTATCTTTCATTTGATTAAGCGGATGGCTGGCATGTACGGCTACCATCAAACGTTCA**<br>**TTACGCAATAAGGTACGTTTAATGGCGGGATCACTGATTTTAAGTCGACCAAATCCG**<br>*benM*<br>**GCATCGATACGGCCTTCTTTTAATGCTTCGGTTTGTGCCTTGGTGCCCATTTCGTAAA**<br>**GTTCAATGCGTAAATTGGGGTGTGCCTGACGATACAAATGAATAATCCGTGGCAGTA**<br>**AACCAAATAGGAGCGAACCCACAAATCCGATACGAATGGTTTTTTCAACCGAGGCAA**<br>**TGCGCTTGGTCATGGAAACCATTTGATCGACATTCGACAATAGTTTGATGGCGTACTG**<br>**ATAAAAAAAGTGACCCTCAGGTGTTGTCTTGACCGGTCTGCTGCCGCGCTCTAAAAGC**<br>**TGAATCCCCAATTCTTCTTCAAGATTTTGAATTTGTCGGCTTAAGGGCGGCTGTGCAA**<br>**TGCATAATTTGTCTGCGGCTTTGGTGAAGCTTTGCTCCTCAACCACAGCCACAAAATA**<br>**GCGGAGATGTCTAAGTTCCAT**AGAGACCTCCTTGGACCTGGTACGTCGCTAGCGGGTAGT<br>BsaI restriction site BsaI recognition sequence GoldenRBS<br>GTGACGTGGCGCAGGTGATGGACTTCATGCTGAC<br>PaqCI restriction site PaqCI recognition sequence |
| $P_{benA/benM}$ | GATGATGGCGAGCATTCACCTGCGGTACTTACGGTCTCTCCAT**TTAAAAATACTCCATAGGT**<br>PaqCI recognition sequence PaqCI restriction site BsaI recognition sequence BsaI restriction site<br>**ATTTTATTATACAAATAATGTGTTTGAACTTATTAAAACATTCTTTTAAGGTATAAACA**<br>**AGCAAGAAAGACAAGAAGAAGGCAGGGGCTTGACCCATTAAATGCTTTCTTCAATTT**<br>$P_{benA/benM}$<br>**GGAAAATTGAAAGCTGAAATGGATATTCGTTTTATTTGTCGGTTCTGCCGTTAAGTAA**<br>**ACATTTTATGCGTTGCGTTGTTTAATTGAATGTTTGACTAAGCACAGCGTTTTGCTCTG**<br>**GCCTAGACAAGTTTCTTATTTTGGAATGTTGGAGAAAGGAT**ATGGTGGCGCAGGTGATG<br>PaqCI restriction site PaqCI recognition sequence<br>GACTTCATGCTGAC |

### CitR

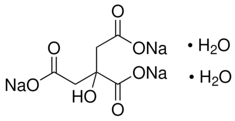


Ligand: Na-citrate
Source: Sigma- Aldrich W302600
CAS ID: 6132-04-3
Stock: 200 g/L
Solvent: $H_2O$
Storage: 25 °C (2 months)

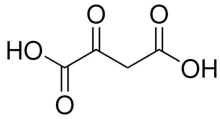


Ligand: oxaloacetic acid
Source: Sigma- Aldrich O4126
CAS ID: 328-42-7
Stock: 100 g/L
Solvent: $H_2O$
Storage: -20 °C (2 months)

#### Response of NatCitR and SynCitR

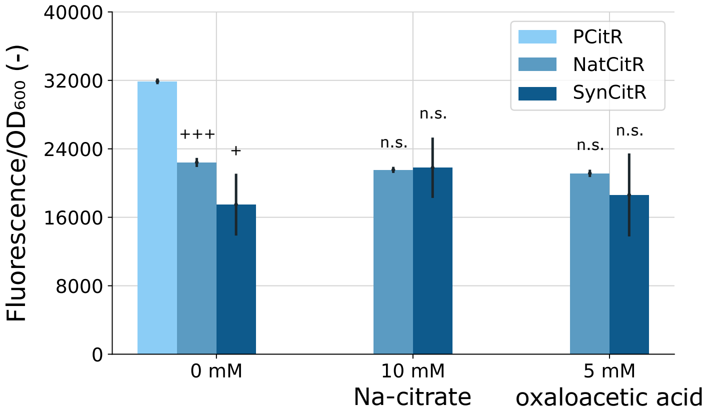


#### Metadata

| | |
|---|---|
| Host | *Escherichia coli TOP10 (F- mcrA Δ(mrr-hsdRMS-mcrBC) Φ80lacZΔM15 ΔlacX74 recA1 araD139 Δ(araleu)7697 galU galK rpsL (StrR) endA1 nupG)* |
| Origin | *Bacillus subtilis* |
| Parts | CitR: GenBank AL009126.3 |
| | $P_{citA/citR}$: Intergenic region between *cit*R and *cit*A; GenBank AL009126.3 |
| Architecture | Natural and Synthetic |
| Experiment | 150 µL EZ Rich medium, 96-well black microtiter plate<br>PerkinElmer Ensight, exc. 588 nm, emm. 633 nm at 30 °C<br>Fluorescence after 24 h of growth |
| Notes | No induction of the biosensors by the two tested ligands |

### CitR

| | |
|---|---|
| *citR* | GATGATGGCGAGCATTCACCTGCGGTAGCTA**TTAGAAATGAAAGTGGCTCAGAAAGTCCA**<br>PaqCI recognition sequence PaqCI restriction site<br>**GGAACTTTTTCTCTTTCTTGTTTTCATACAGTGCAATGGCATATGCACCGGCATAAGG**<br>**CAGCTGAACGCTCTGATACGGAATGCGAATCATTTGTTTTTCTGCCAGTTCACGTTTA**<br>**ACGGTGCTCAGCGGCAGAAAGCTAACGCCCAGACCTTCTTTGATAAAGCGCTTGGTA**<br>**ATATGTGTCTGGGTAACTTTCATGGTACGAACAAACGGAAAGGTCATACGAACCTGA**<br>**CGCAGCAGATCATCCCAGTAATCAGGATGATTGTGGGTCAAAAGCAGGTATTGTTCG**<br>**AGCACCTCTTTGGCATCAATTTCGTTGTCTTCTATGAAACGCTTATCGGGCGGGGCAA**<br>**CAAGGACGACAGGGTCTTTGTATAAGCAGTGGCAGCTTAAAGAAGAAGACTGTACCT**<br>**TTAAACAGCTTAGTCCGATATCTGCTTCACCGGCTTTAATCAGGCTTGCAATTTCTGC**<br>*citR*<br>**GCTTTCAAAAATGGTAACGGCCATTTCGGTTTCGGTATTCATTGCGGTATAACGTTTC**<br>**ATAACGCTCGGCAGAACGGTATCTGCAATCAGCGGACTAACTGCCAGCTGCAGGGTC**<br>**TGACTATAACCCTGACGAACACGATGCAGTTCTGCCATGCTATTTTCATAATCATCCA**<br>**GCAGCCTGAGCGCATACGGCAAATACGCCCTGCCTTCATCAGTCAGCTGGATTTGCCT**<br>**GCCTTTACGCTCGAACAGCTTGCAGCTGATTTCTTTTTCTAATTGCTTGATATGTACGG**<br>**TCACAGTAGGCTGAGATAAGAAAAGCGTTTCCGCCGTTTTTCGGAAGTTCTCATATTT**<br>**TGCAGCGGTCACAAAGGTGTGAAGCCATTTGAAATCCAT**AGAGACCTCCTTGGACCTGG<br>BsaI restriction site BsaI recognition sequence GoldenRBS<br>TACGTCGCTAGCGGGTAGTGTGACGTGGCGCAGGTGATGGACTTCATGCTGAC<br>PaqCI restriction site PaqCI recognition sequence |
| P$_{citA/citR}$ | GATGATGGCGAGCATTCACCTGCGGTACTTACGGTCTCTCCAT**TTCTATTCTCCCTCTGATT**<br>PaqCI recognition sequence PaqCI restriction site BsaI recognition sequence BsaI restriction site<br>**AATATTTTTAATTAATTCCCTTTAAAATATTGATTATTTTTTAAATATTATATTTACTAT**<br>P$_{citA/citR}$<br>**AATAACAGAAAAGGATAGGGGGAATACAA**ATGGTGGCGCAGGTGATGGACTTCATGCTG<br>PaqCI restriction site PaqCI recognition sequence<br>AC |

#### CysB

**No ligands tested**

Response NatCysB and SynCysB

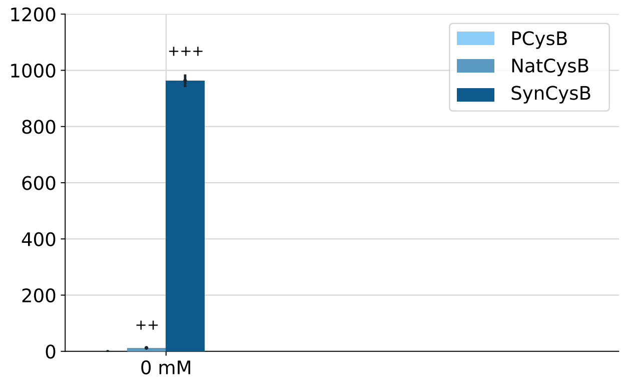


Metadata

| | |
|---|---|
| Host | *Escherichia coli TOP10 (F- mcrA Δ(mrr-hsdRMS-mcrBC) Φ80lacZΔM15 ΔlacX74 recA1 araD139 Δ(araleu)7697 galU galK rpsL (StrR) endA1 nupG)* |
| Origin | *Burkholderia cenocepacia* |
| Parts | CysB: GenBank AM747720 |
| | $P_{cysI/cysB}$: Intergenic region between *cys*B and *cysI;* GenBank AM747720 |
| Architecture | Natural and Synthetic |
| Experiment | 150 µL EZ Rich medium, 96-well black microtiter plate<br>PerkinElmer Ensight, exc. 588 nm, emm. 633 nm at 30 °C<br>Fluorescence after 24 h of growth |
| Notes | / |

### CysB

| | |
|---|---|
| *cysB* | GATGATGGCGAGCATTCACCTGCGGTAGCTA**TTACAGTTCATAGCTTTCTGCTTCACCTTT**<br>PaqCI recognition sequence PaqCI restriction site<br>**CAGTGCCTGTTCAATCAGTTTACGATTCAGTGTCGGACTCAGCAGTTCAACCAGGGTA**<br>**TAAACATAGCTACGCAGATATGCACCCTGTTTCAGTGCAACACGGGTAACATTGCTAC**<br>**CAAACAGGTGACCAACCGGAATCAGACGCAGACCACGATCACGTTCCGGATTAAATG**<br>**CAATATCTGCCATAATACCCACACCTAAACCTAATTCCACATAGGTTTTGATCACATC**<br>**GGCATCAATTGCTTCCAGAACAATATCCGGACTCAGACCACGCAGTGCAAAGGCATG**<br>**ATTGATTTTTTTGCGACCTGCAAATGCATCATCATAGGTAATCAGCGGATACTGTGCC**<br>**AGATCATCTAAGGTAACCGGTTTACGTTCCAGCAGCGGATGATCTGCCGGAACAACT**<br>**GCTGCATGATGCCACTGAAAACACGGCAGGCTAACCAGTTCTTTATAATCGCTAATTG**<br>*cysB*<br>**CTTCGGTTGCAATTGCCAGATCTGCCTGATCATGAATAACCATTTCTGCAACCTGTGT**<br>**CGGGCTACCCTGCAGAATGCTCAGATGCACTTTCGGAAAGCGTTTTTTGAATTCTGCA**<br>**ATTGCTGCAGGCAGGCTATAACGTGCCTGTGTATGGGTTGCTGCAATGGTCAGATTA**<br>**CCCTGATCCTGTGCTGCATAATCTTTGCCCACACGTTTCAGGCTTTCAACTTCTTGCAG**<br>**AATACGTTCAACGCTTGCCAGAATAATACGACCCGGTTCGGTCAGACTACGAACACG**<br>**TTTACCATGACGGGTAAAGATTTCAACGCCCAGTTCATCTTCCAGTTCAATAATTGCT**<br>**TTGCTAACACCAGGCTGGCTGGTATACAGTGCTTTTGCTGCTTCGGTCAGGTTAAAAT**<br>**TCTGGCGAACTGCTTCACGAACAAAACGAAACTGATGCAGGTTCAT**AGAGACCTCCTT<br>BsaI restriction site BsaI recognition sequence<br>GGACCTGGTACGTCGCTAGCGGGTAGTGTGACGTGGCGCAGGTGATGGACTTCATGCTGAC<br>GoldenRBS PaqCI restriction site PaqCI recognition sequence |
| $P_{cysI/cysB}$ | GATGATGGCGAGCATTCACCTGCGGTACTTACGGTCTCATCAT**TTATAACCCTTCCGCATAT**<br>PaqCI recognition sequence PaqCI restriction site BsaI recognition sequence BsaI restriction site<br>**CAACAGAATTTTTTAGTCGTTTGAAATATAAGGCGAGTTTATTACGATTCACCGGAGT**<br>$P_{cysI/cysB}$<br>**TTTTCAAATATGGATATCTGTTTTCGTCATTAGCAATCCAGCCGGCAGCGGATGCGGC**<br>**GGTGCGGCGGAACGGAGATTCGGGCGCGACGTGGCTAGGACGTCGCGCGGTCACCA**<br>**CGAAAACCCTGGGGTCCCCGA**ATGGTGGCGCAGGTGATGGACTTCATGCTGAC<br>PaqCI restriction site PaqCI recognition sequence |

### GltC:α-Ketoglutaric acid

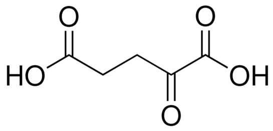


Ligand: α-Ketoglutaric acid
Source: Sigma- Aldrich 75890
CAS ID: 328-50-7
Stock: 100 g/L
Solvent: $H_2O$
Storage: 2-8 °C (2 months)

#### Response NatGltC and SynGltC

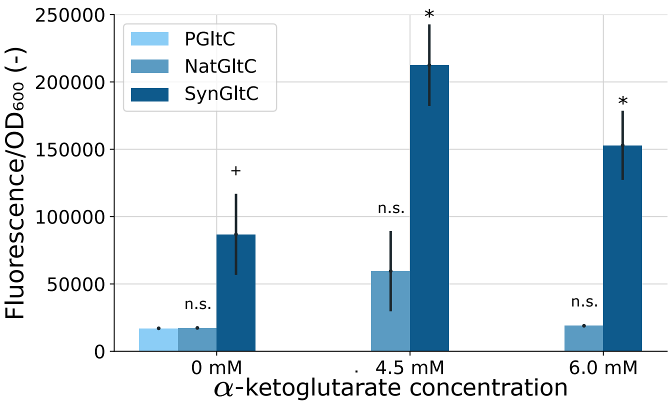


#### Metadata

| | |
|---|---|
| Host | *Escherichia coli TOP10 (F- mcrA Δ(mrr-hsdRMS-mcrBC) Φ80lacZΔM15 ΔlacX74 recA1 araD139 Δ(araleu)7697 galU galK rpsL (StrR) endA1 nupG)* |
| Origin | *Bacillus subtilis* |
| Parts | GltC: GenBank AL009126.3 |
| | $P_{gltA/gltC}$: Intergenic region between *Glt*C and *gltA;* GenBank AL009126.3 |
| Architecture | Natural and Synthetic |
| Experiment | 150 µL EZ Rich medium, 96-well black microtiter plate<br>PerkinElmer Ensight, exc. 588 nm, emm. 633 nm at 30 °C<br>Fluorescence after 24 h of growth |
| Notes | / |

### GltC:α-Ketoglutaric acid

| | |
|---|---|
| *gltC* | GATGATGGCGAGCATTCACCTGCGGTAGCTA**TTACTGGTACTGTTCCAGTTTGCTAAAAA**<br>PaqCI recognition sequence PaqCI restriction site<br>**ACTGGATGACAAATTCATAAAAATCATTTGCGCTCGGTGCCAGTTCACGATTTTTCGG**<br>**TTTGATAATACCAACGGTACGTTTAACCTGCGGAAATTCAATCGGGATTTTAACTGTA**<br>**AAACGCGGTGTGGTTTCTGCAAAGGTGCTTTCCGGCAGCAGGGTAACACCCATACCT**<br>**GCGCTAACCAGACCTTTAATTGCATCCAGATCTTCACCTTCGGTGCTAACCAGCGGTG**<br>**CAAAACCTGCCTGTTTACAGGTATCAATTGCCATTTCACGCAGAACAAAACCTTCCGG**<br>**AAACAGAACAAACTGATCATTGCGCAGATCAATCAGATGAACGGTTTTCTGTTTTGCC**<br>**AGCGGATGATTCAGCGGAACCAGTGCATAGATTTTCTCGGTAAACAGGATTTTGCCG**<br>**GTAATATCTGAAAAATTGGTCGGAACAGGACCCAGCAGTGCCAGATCAATATCGCGG**<br>*gltC*<br>**TTGCGAACTGCTTCAATCAGGAATTTATAGCTACCCTGACGCAGCAGAAATTCGACGT**<br>**GCGGATATTCTTCTTTAAATGCGCTAATAACGGTCGGCAGCAGCTGGCTTGCCAGGCT**<br>**GGTCGGAAAACCAATTTTAACGGTGCCACGATGCGGATCCAGATATTCATCGATTTGT**<br>**TCTTTGGCGTAATCAATGGCTTTCATGGCGGTTTTAACATGGATCAGAAATTCTTTAC**<br>**CAATCGGGGTCAGTTTAATGTTACGACCTTCACGTTCAAACAGGGTAACGTTCAGTTC**<br>**TTCTTCCAGATTTGCAATCTGACGGCTAATTGCGCTCTGTGCAACATGCAGATGATCT**<br>**GCTGCTTCGCTAACATGTTCACGTTCTGCAACTTCCATGAAATAGCGCAGCTGACGCA**<br>**GTTCCAT**AGAGACCTCCTTGGACCTGGTACGTCGCTAGCGGGTAGTGTGACGTGGCGCAGG<br>BsaI restriction site BsaI recognition sequence GoldenRBS PaqCI restriction site PaqCI recognition sequence<br>TGATGGACTTCATGCTGAC |
| $P_{gltA/gltC}$ | GATGATGGCGAGCATTCACCTGCGGTACTTACGGTCTCTCCAT**GTTTGTCTCACATCCATCT**<br>PaqCI recognition sequence PaqCI restriction site BsaI recognition sequence BsaI restriction site<br>**ATCTCATTTTGAGATTCTTTTGATCTAAATTATATATTGTTTATATCGTTTTGAAAACC**<br>$P_{gltA/gltC}$<br>**TACAATGATTATAGAGTTGTTAGATTTTATGACCGGTATTATCGGAAATTGATCGGGG**<br>**GAGAGGAATT**ATGGTGGCGCAGGTGATGGACTTCATGCTGAC<br>PaqCI restriction site PaqCI recognition sequence |

### LysG:L-lysine

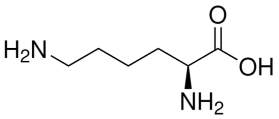

Ligand: L-lysine
Source: Sigma- Aldrich L5501
CAS ID: 56-87-1
Stock: 100 g/L
Solvent: $H_2O$
Storage: 25 °C (2 months)

Response of NatLysG and SynLysG

**No data available**

Metadata

| | |
|---|---|
| Host | *Escherichia coli TOP10 (F- mcrA Δ(mrr-hsdRMS-mcrBC) Φ80lacZΔM15 ΔlacX74 recA1 araD139 Δ(araleu)7697 galU galK rpsL (StrR) endA1 nupG)* |
| Origin | *Corynebacterium efficiens* |
| Parts | MetR: GenBank BA000035.2 |
| | $P_{lysE/lysG}$: Intergenic region between *Lys*G and *Lys*E*;* GenBank BA000035.2 |
| Architecture | Natural and Synthetic |
| Experiment | 150 µL EZ Rich medium, 96-well black microtiter plate<br>PerkinElmer Ensight, exc. 588 nm, emm. 633 nm at 30 °C<br>Fluorescence after 24 h of growth |
| Notes | / |

#### LysG:L-lysine

| | |
|---|---|
| *LysG* | GATGATGGCGAGCATTCACCTGCGGTAGCTA**TTAGGTACGCAGACCTGCACGTGCTGCAT**<br>PaqCI recognition sequence PaqCI restriction site<br>**CAACAACGGCATCGGTCAGACGTGCCAGCAGACGGCTTTCCAGACGCCAACGCTGCC**<br>**AATACAGCGGTGTATCAACAACTTTCTCATCCAGCTGAACAACATCACCGGCTGCCAG**<br>**CATCGGTGCAGCCTGTGCTTCCGGCAGCAGACCCCAACCTAAACCTAAACGAACTGCT**<br>**TCACCAAAACCTTCGGCACTCGGAACAACGCTCACACGACGACGGGCAACTGCACCA**<br>**TCAACACGACCTTCCAGATCACGATCCTGCAGAACATCATTCGGACCAAAACGCAGA**<br>**ACCGGCATACGAACCCAATCCGGCTGACCATCAACGGTATAACGTGCACGCAGTTCC**<br>**GGTGTGGCAACCGGCAGATGACGCATAACACCCAGACGCAGAACTTCGCAACCTGCA**<br>**ACCGGATCTGCTTCACGTGTAACGGCACCCAGAACGCTACCACGACGCAGCAGGCTC**<br>*lys*G<br>**AGGGTATGTGCTTCATCTTCAACACGCAGTGTCAGGGTAACTGCACCCCAATGTGCAA**<br>**CTTCGGCAAAAACAGGCGGAAACCAGGTGCTCAGGCTATCTGCATTAATTGCAACGG**<br>**TCAGCGGAATTTCATCCAGACGTTCTGCCAGCTGTTCACGGGTTTCTGCCTGCAGCAG**<br>**TGCCATTTTACGTGCTGCCTGAACCAGAACTTCACCGGCTTCGGTTGCAACTGCAGGC**<br>**TGGGTACGACTAACCAGAACACGACCAACTGATTTTTCCAGTGCTTTAATACGCTGGC**<br>**TAACTGCGCTCGGGCTAATGCTCAGTGCCAGGCTTGCATTTTCAAAGCTACCTTCATC**<br>**AATAATGGTCAGCAGGGTATCCAGATGAATCGGGTTCAT**AGAGACCTCCTTGGACCTGG<br>BsaI restriction site BsaI recognition sequence GoldenRBS<br>TACGTCGCTAGCGGGTAGTGTGACGTGGCGCAGGTGATGGACTTCATGCTGAC<br>PaqCI restriction site PaqCI recognition sequence |
| $P_{lysE/lysG}$ | GATGATGGCGAGCATTCACCTGCGGTACTTACGGTCTCTTCAT**GAAGCAATTCTAAACCATG**<br>PaqCI recognition sequence PaqCI restriction site BsaI recognition sequence BsaI restriction site<br>**TTCAGTCTCTATCATTTTACTTAATGTGGTTTGTGCGCGAACATGCGGGAC**ATGGTGGC<br>$P_{lysE/lysG}$ PaqCI restriction site<br>GCAGGTGATGGACTTCATGCTGAC<br>PaqCI recognition sequence |

#### NagR:salicylate

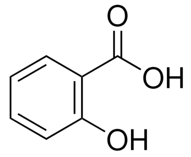


Ligand: salicylate
Source: Sigma- Aldrich 69453
CAS ID: 69-72-7
Stock: 20 g/L
Solvent: EtOH
Storage: -20 °C (2 months)

Response of NatNagR and SynNagR

**No data available**

Metadata

| | |
|---|---|
| Host | *Escherichia coli TOP10 (F- mcrA Δ(mrr-hsdRMS-mcrBC) Φ80lacZΔM15 ΔlacX74 recA1 araD139 Δ(araleu)7697 galU galK rpsL (StrR) endA1 nupG)* |
| Origin | *Ralstonia sp.* |
| Parts | NagR: GenBank AF036940.2 |
| | $P_{nagA/nagR}$: Intergenic region between *Nag*R and *Nag*A*;* GenBank AF036940.2 |
| Architecture | Natural and Synthetic |
| Experiment | 150 µL EZ Rich medium, 96-well black microtiter plate<br>PerkinElmer Ensight, exc. 588 nm, emm. 633 nm at 30 °C<br>Fluorescence after 24 h of growth |
| Notes | / |

#### NagR:salicylate

| | |
|---|---|
| *nag*R | GATGATGGCGAGCATTCACCTGCGGTAGCTA**TTAGGCTTCACTAAACAGTTCAACAAACA**<br>PaqCI recognition sequence PaqCI restriction site<br>**GCTGACGCAGCCACATATTACCCGGATCGCGGTTATATTTGGCGTGCCAAAAAAGAT**<br>**TAATGGCAATATCCGGCAGTTTTGCCGGATGCGGACTGGTGGTCAGACCAAACGGAA**<br>**CTTCACAACGAACTGCAAAACGCTGCGGAACGGTTGCAATCAGATCGGTGCTATGCA**<br>**GAATCGGACCAATTGCAATAAAATGCGGAACAACCAGACGCATACGACGTTTAATAC**<br>**CTGCACGTTCCAGCAGACCATCAACTTCACCATGACCGGTATTCAGGGCAACAACACC**<br>**AACATGTTCCAGTTCGCTAAACTGTTTCAGACTCATCGGGCTTTTTGCGCTCGGATGA**<br>**TCTTTGCGAAACATACAAACATAACGATGACGAAACAGACGACGCTGAAAAAAACCT**<br>**GTCTGCAGTTCCGGCAGCAGACCCAGGGCCAGATCAACTGCGCCTGATTCCATATCTT**<br>*nag*R<br>**CTTTCAGATTACCTGCATTCGGACGCAGGGTGCTAATCTGAATATGCGGTGCACGCTG**<br>**TGCCAGTGCTTCCATCAGCGGAGGCATGAAATACATCTCGCCAATATCGGTCATTGCC**<br>**AGATTAAAGGTACGGGTGCTTGCAAACGGATCAAAGCTATCACGTGTGGTCAGTGCG**<br>**GTCTGCAGGGTATTCAGTGCATAAATAACCGGTTCTGCCAGATGCAGTGCATACGGT**<br>**GTCGGTTCCATACCTTTGCTGGTACGCAGAAACAGGTCATCTTTCAGTGCTGCACGCA**<br>**GACGTTTCAGGCTATTGCTAACTGCCGGTTGGGTCAGACCCAGTTTTTCGCCTGCGGT**<br>**GCTAACGCTACGATCCAGCAGCAGCTGATTAAAAACAACCAGCAGATTCAGATCAAT**<br>**ATCACGCAGATCCAT**AGAGACCTCCTTGGACCTGGTACGTCGCTAGCGGGTAGTGTGACGT<br>BsaI restriction site BsaI recognition sequence GoldenRBS PaqCI restriction site<br>GGCGCAGGTGATGGACTTCATGCTGAC<br>PaqCI recognition sequence |
| $P_{nagA/nagR}$ | GATGATGGCGAGCATTCACCTGCGGTACTTACGGTCTCTCCAT**GATGCCTCACCATTATTCA**<br>PaqCI recognition sequence PaqCI restriction site BsaI recognition sequence BsaI restriction site<br>**TGCTGGTGATTTTAACTATCAGACTTGATCTATAGCGCTATACCGATCGACGCGCCAG**<br>$P_{nagA/nagR}$<br>**AATCGCAGCCATTCGGAGACAACTGAAAAAAGAGCTTGC**ATGGTGGCGCAGGTGATGG<br>PaqCI restriction site PaqCI recognition sequence<br>ACTTCATGCTGAC |

**NahR:salicylate**

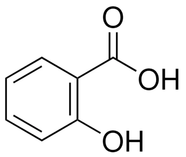


Ligand: salicylate
Source: Sigma- Aldrich 69453
CAS ID: 69-72-7
Stock: 20 g/L
Solvent: EtOH
Storage: -20 °C (2 months)

Response of NatNahR and SynNahR

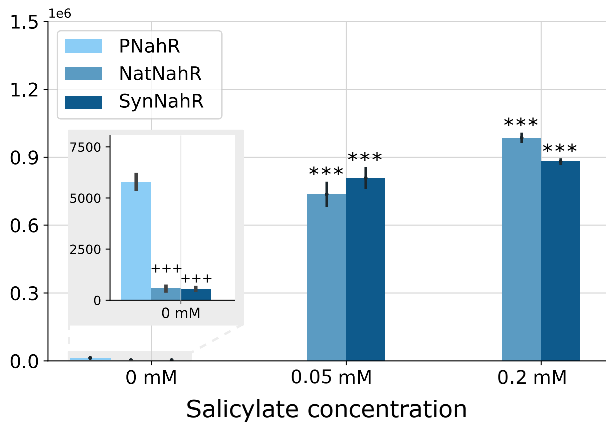


Metadata

| | |
|---|---|
| Host | *Escherichia coli TOP10 (F- mcrA Δ(mrr-hsdRMS-mcrBC) Φ80lacZΔM15 ΔlacX74 recA1 araD139 Δ(araleu)7697 galU galK rpsL (StrR) endA1 nupG)* |
| Origin | *Pseudomonas putida* |
| Parts | NahR: Addgene pAJM.771 |
| | $P_{nahRmarionette}$: Addgene pAJM.771 |
| Architecture | Natural and Synthetic |
| Experiment | 150 µL EZ Rich medium, 96-well black microtiter plate<br>PerkinElmer Ensight, exc. 588 nm, emm. 633 nm at 30 °C<br>Fluorescence after 24 h of growth |
| Notes | / |

#### NahR:salicylate

| | |
|---|---|
| *nahR* | GATGATGGCGAGCATTCACCTGCGGTAGCTA**TCAATCCGTAAACAGGTCAAACATCAGTT**<br>(PaqCI recognition sequence; PaqCI restriction site)<br>**GCCGCAACCAAATATTGGCTAGGTCCTTGTGGTACTTCGCATGCCAGAACATGTTGAT**<br>**GGCTATTTCAGGCAAGACGACTGGGTGCGGCAAGGCGCTTAGGCCGAAGGGCTCTAC**<br>**GCAGCAGTCGGCTAAACATATCGGCACAGTGGCGAGCAGATCGGTGCGCTGGAGGA**<br>**TGTGGCCAACGGCGGCGAAGTGCGGCACTTCCAGACGGATGTCGCGCCGGATGCCGA**<br>**CCCGTGTCATGTACGTGTCCACCTCGCCGTGGCCGGTGCCAGCGGCGATGACACGCA**<br>**CGTGGCCGTAGGAACAGAAGCGCTCCAGAGTCAGGGGTTCGCGGGTGACTGGATGG**<br>**TCCTTGCGACATAGGCACACGTAGTGATTCTGGAGCAGCCGGCGCTGAAAGAAGCCA**<br>**GTTTGCAGATTGGGAAGCAGGCCCACGGCCAAGTCCACGGTTCCGTTCTGCAAGGCC**<br>*nahR*<br>**TGCATCAGGCTCATCGAACTGTCGCGCACCGTACTGATCACGCAATTGGGGGCCTGG**<br>**TGAGCCAGCGCATCCATCAGCCGCGGCATGAAGTAGATCTCGCCAATGTCGGTCATG**<br>**GCCAGGGTGAAGGTACGCTCGCTGGTCAGCGGATCGAAGCTTTCATGGTGCTGTAGG**<br>**GCGTTGCGCAGTGCGTGCATGGCCGAAGTGACGTGCTCGGCCAGATGCGCGGCATAG**<br>**GGTGTGGGTTCCATTCCCTGATGTGTGCGCACGAAGAGTGGGTCCTGTAGCGAGGTG**<br>**CGCAGGCGTTTCAGCGCATTGCTCACGGCAGGCTGGGTCAGGCCCAGGTTCTCCGCA**<br>**GTGACAGATACGCGTCTGTCGACCAGCAACTGGTTGAACACCACCAGCAGGTTTAAA**<br>**TCCAGGTCACGCAGTTCCAT**AGAGACCTCCTTGGACCTGGTACGTCGCTAGCGGGTAGTGT<br>(BsaI restriction site; BsaI recognition sequence; GoldenRBS)<br>GACGTGGCGCAGGTGATGGACTTCATGCTGAC<br>(PaqCI restriction site; PaqCI recognition sequence) |
| $P_{nahR\ mar.}$ | GATGATGGCGAGCATTCACCTGCGGTACTTACGGTCTCTCCAT**GGGGCCTCGCTTGGGTTA**<br>(PaqCI recognition sequence; PaqCI restriction site; BsaI recognition sequence; BsaI restriction site)<br>**TTGCTGGTGCCCGGCCGGGCGCAATATTCATGTTGATGATTTATTATATATCGAGTGG**<br>$P_{nahR\ mar.}$<br>**TGTATTTATTTATATTGTTTGCTCCGTTACCGTTATTAACAGCTGTCACCGGATGTGCT**<br>**TTCCGGTCTGATGAGTCCGTGAGGACGAAACAGCCTCTACAAATAATTTTGTTTAATA**<br>**CTAGAGAAAGAGGGGAAATACTAG**ATGGTGGCGCAGGTGATGGACTTCATGCTGAC<br>(PaqCI restriction site; PaqCI recognition sequence) |

**OccR:D-octopine**

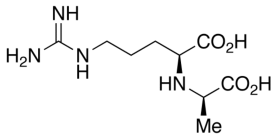


Ligand: D-octopine
Source: Toronto Research Chemicals O239850
CAS ID: 34522-32-2
Stock: 2.6 g/L
Solvent: $H_2O$
Storage: 2-8 °C (2 months)

Response of NatOccR and SynOccR

**No data available**

Metadata

| | |
|---|---|
| Host | *Escherichia coli TOP10 (F- mcrA Δ(mrr-hsdRMS-mcrBC) Φ80lacZΔM15 ΔlacX74 recA1 araD139 Δ(araleu)7697 galU galK rpsL (StrR) endA1 nupG)* |
| Origin | *Agrobacterium tumefaciens* |
| Parts | OccR: GenBank AF242881.1 |
| | $P_{occQ/occR}$: Intergenic region between *occ*R and *occ*Q*;* GenBank AF242881.1 |
| Architecture | Natural and Synthetic |
| Experiment | 150 µL EZ Rich medium, 96-well black microtiter plate<br>PerkinElmer Ensight, exc. 588 nm, emm. 633 nm at 30 °C<br>Fluorescence after 24 h of growth |
| Notes | / |

#### OccR:D-octopine

| | |
|---|---|
| *occ*R | GATGATGGCGAGCATTCACCTGCGGTAGCTA**TTACTCCATTAGGCCGTTCTGCTTCATCAA**<br>PaqCI recognition sequence PaqCI restriction site<br>**GTCATCATGAAACCTCCAGAATTCGGTTGTGAAACGATCGACGATGGTTGAGGGAGC**<br>**GCCAATTGCTGACCGGACTTCGAGGAATCCGGCGTCAATGAAGATCGAGAACGGTCG**<br>**CAGTACGATCCTGTCCGTGAACTCGATCGCCGCGGCTGGATCGATAATTGCGATCCCG**<br>**GCGCCTTCGCGGACGAGACTTAGCGCAGTATGCGACAGGCTCACTTCAATTGACGGC**<br>**CGGCGTTGAATACCACCAATCGCCACCTCTACCCGCATGGCGAAGAGAGTGCCAGTC**<br>**TCCTGTTTTATAATACGCTCACCGGCAAGGTCCTGTGGCGTAACACGATCCAGACCTG**<br>**CCAGACGATGACCCATCGGAACTGCAACAACTGCTGGCAGGCTACGGGTTTCAATCA**<br>**GAAAACCCTGACGTTCCTGTGGGCCATCGGCATAACCGATGTCGGCCCTGCCGGACG**<br>*occ*R<br>**CAACGGCTTCCATGACCATGCTTGAGGGCAGTCCCATTAGGGATACCTGGAGATTTG**<br>**GTCTGTCACGGATGAACTGAGCAAGAAACCGCGGCAAGAGGCCGTTTGCCAGAGCA**<br>**GGCATTGCAGCAATGCGGAGCGTCCCCGCTGCCTGCCTGCCGATGTCGGCAGCCAGG**<br>**TTGCCTATATGATTAAGCCCGACGAACGCCCGATCGACCTCTTTCCACAGCGTCTTTG**<br>**CCTCCTGTGTCGGGATAATATGGTTCCCACGCCTCTCGAAGAGCTGCAGTTTTGTCGC**<br>**CTGTTCAAAGTCCTTGATTAGGCGACTGATGGCCGGCTGAGTCACCAGCATTAGTTCA**<br>**GCCGCCGCCGTCATTTGCCCCGTCAGCATGACTGCCCGGAACGCCTCGACCTGCCTGA**<br>←<br>**GATTCAT**AGAGACCTCCTTGGACCTGGTACGTCGCTAGCGGGTAGTGTGACGTGGCGCAGG<br>BsaI restriction site BsaI recognition sequence GoldenRBS PaqCI restriction site PaqCI recognition sequence<br>TGATGGACTTCATGCTGAC |
| $P_{occQ/occR}$ | GATGATGGCGAGCATTCACCTGCGGTACTTACGGTCTCTTCAT**AAACGCACCATAACATCTG**<br>PaqCI recognition sequence PaqCI restriction site BsaI recognition sequence BsaI restriction site<br>**CTTATTCTTGCCCGGTCATTATGAATTTGACCGAATGCATATCGAATGTAAAGCTCAC**<br>$P_{occQ/occR}$<br>**CCTATAAATCACAACTCTTCCGGGCCAACCGGGATCAGACGTGCGGCGAAACGGTCG**<br>**CGCGCAGGGAACGCT**ATGGTGGCGCAGGTGATGGACTTCATGCTGAC<br>PaqCI restriction site PaqCI recognition sequence |

#### PqsR:2-heptyl-4-quinolone concentration

Ligand:2-heptyl-4-quinolone concentration
Source: Sigma- Aldrich SML0747
CAS ID: 40522-46-1
Stock: 5 g/L
Solvent: DMSO
Storage: 2-8 °C (2 months)

Response of SynPqsR

**No data available**

Metadata

| | |
|---|---|
| Host | *Escherichia coli TOP10 (F- mcrA Δ(mrr-hsdRMS-mcrBC) Φ80lacZΔM15 ΔlacX74 recA1 araD139 Δ(araleu)7697 galU galK rpsL (StrR) endA1 nupG)* |
| Origin | *Pseudomonas aeruginosa* |
| Parts | PqsR: GenBank AE004091.2 |
| | $P_{pqsA}$: Promoter of *pqs*A:<br>GenBank AE004091.2 |
| Architecture | Synthetic |
| Experiment | 150 µL EZ Rich medium, 96-well black microtiter plate<br>PerkinElmer Ensight, exc. 588 nm, emm. 633 nm at 30 °C<br>Fluorescence after 24 h of growth |
| Notes | / |

#### PqsR:2-heptyl-4-quinolone concentration

| | |
|---|---|
| *pqs*R | GATGATGGCGAGCATTCACCTGCGGTAGCTA**TTATTCCGGTGCTGCACGCTGACGATATG**<br>PaqCI recognition sequence PaqCI restriction site<br>**CCAGTGCTTTCGGACCACTACGACGCTGTGCGGTTTCAACAATGCTCGGCTGCCATGC**<br>**CGGTGCATCATCAAAACGCTGGCGACCCAGTTCACGCAGACGCTGACGTGCACTTTC**<br>**CAGAAAACGCAGAAAGCTACGTTCGCTTTCCAGTGCGGTGTTATAATAGCAGTACAC**<br>**TTTGGTATCAATACCACCAGGTTCATACAGTTCGCTCAGAACTGCCAGGGTGCCATTA**<br>**CGCAGACGTTCTTCAACAAAATAATGCGGTGCAATACCCCAACCAACACCGGCTTCCA**<br>**CCAGACGCAGCATATCATCGAAGTTTTCCACAAACAGAACTTTATCGCTAACCGGACG**<br>**CAGCAGATTACTATGCTGACCGCTACGGCTACCCAGGCTAATCTGACGATAATTGGCC**<br>**AGGCTGGCAATGCTATGCAGGCTTGCATTACACAGCGGATGCTGCGGATGTGCAACA**<br>*pqs*R<br>**ACAAATGCTTTGGTATAACCCAGAACGCACTGATTAAAGCGGCTAATTTTCAGTTCTT**<br>**CATCAATGGTGATGGCGATATCAATTTCGGCATTATCCTGTTTAATGGTTGCCAGGCT**<br>**ATCTGCCGGACTGGTACGAATCAGGCTAACCATGTTAAAATCATCCAGCAGCACGCT**<br>**GCTAACGGTATCACAAAAGCTAGGCGGAATTGCGGTATCCAGCAGAACACGCAGATT**<br>**ACGCGGACCTTTATTCAGATTGAAGGCAATATCACCAATCAGCTGCTGATAATTCAGC**<br>**AGGCTACGCATATACGGAATCAGACGCAGTGCCTGTTCGGTCGGTTCAACTTTATAAC**<br>**CATCACGACGAACCAGTTCAACACACAGATCAATTTCCAGATTGCTAACTGCTGAGCT**<br>**AACTGCGGTATGGCTTTTACGCAGAATACGTGCTGCGCTGCTAATGCTACCGCTTGCA**<br>**ATAACCTGCAGAAACATATTCACGTGGTTCAGATTATGAATCGGCAT**AGAGACCTCCTT<br>BsaI restriction site BsaI recognition sequence<br>GGACCTGGTACGTCGCTAGCGGGTAGTGTGACGTGGCGCAGGTGATGGACTTCATGCTGAC<br>GoldenRBS PaqCI restriction site PaqCI recognition sequence |
| P$_{pqsA}$ | GATGATGGCGAGCATTCACCTGCGGTACTTACGGTCTCTGCAT**CCCGGTGGGTGTGCCAAA**<br>PaqCI recognition sequence PaqCI restriction site BsaI recognition sequence BsaI restriction site<br>**TTTCTCGCGGTTTGGATCGCGCCGATTGCCGCGGCCTACGAAGCCCGTGGTTCTTCTC**<br>P$_{pqsA}$<br>**CCCGAAACTTTTTCGTTCGGACTCCGAATATCGCGCTTCGCCCAGCGCCGCTAGTTTCC**<br>**CGTTCCTGACAAAGCAAGCGCTCTGGCTCAGGTATCTCCTGATCCGGATGCATATCGC**<br>**TGAAGAGGGAACGTTCTGTC**ATGGTGGCGCAGGTGATGGACTTCATGCTGAC<br>PaqCI restriction site PaqCI recognition sequence |

### TsaR:*p*-toluenesulfonate

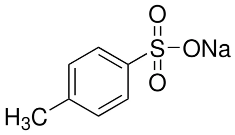


Ligand: *p*-toluenesulfonate
Source: Sigma- Aldrich 152536
CAS ID: 657-84-1
Stock: 40 g/L
Solvent: $H_2O$
Storage: 2-8 °C (2 months)

#### Response of NatTsaR and SynTsaR

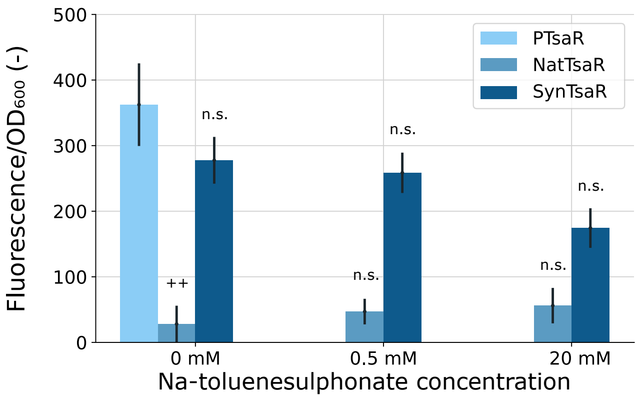


#### Metadata

| | |
|---|---|
| Host | *Escherichia coli TOP10 (F- mcrA Δ(mrr-hsdRMS-mcrBC) Φ80lacZΔM15 ΔlacX74 recA1 araD139 Δ(araleu)7697 galU galK rpsL (StrR) endA1 nupG)* |
| Origin | *Comamonas testosteroni* |
| Parts | TsaR: GenBank AH010657.3 |
| | $P_{tsaM/tsaR}$: Intergenic region between *tsa*R and *tsa*M*;* GenBank AE006468.2 |
| Architecture | Natural and Synthetic |
| Experiment | 150 µL EZ Rich medium, 96-well black microtiter plate<br>PerkinElmer Ensight, exc. 588 nm, emm. 633 nm at 30 °C<br>Fluorescence after 24 h of growth |
| Notes | / |

### TsaR:p-toluenesulfonate

| | |
|---|---|
| *tsa*R | GATGATGGCGAGCATTCACCTGCGGTAGCTA**TTAACCGGTCTGCAGGGCATGATGCTGAA**<br>PaqCI recognition sequence PaqCI restriction site<br>**TCCAACGAATCAGACCTGCTGCTGCCGGTGTAACAGGCAGATCATGACGACGCAGAA**<br>**CATAAATGGTCGGATTCGGCAGTGCATCCTGCAGCGGAATGCTACACAGCTGATCTTT**<br>**AAAGGCATTACGTTCATACAGTGTACGAGGCATGGTGGTCAGCAGATCGCTATGTGC**<br>**AACAACACCAGGCAGTGCCAGAAAACTTTCACAAACTAAACCCAGTTTCGGTTCCGG**<br>**CAGACCATAACGTGCAAATGCATTACGAATAATTGCACCCGGACCACGCGGTGCGCT**<br>**GCTAAATGCCCAACGACATTCTTGCAGTTCTGCCAGACGGGTTGCATTTGCCATCGGA**<br>**TGCTGACGCTGACCAACAATAACAACATCACTAACATACAGCGGCTGGGCTTCCAGA**<br>**TCGGTATCAATATCATGTTTATGTGCTGCGGTCAGGGCAAAATCCAGGGTGCCATCAC**<br>*tsaR*<br>**GCAGTTGAGGGCTAACTGCCGGATACATACCATCACGAACATTAACGGTAACATCCG**<br>**GAAATTCACGTGCAAAGCTTGCCAGTGCCAGCGGCAGTGCTGCCAGTGCAATTGCCG**<br>**GACTTGCTGCAAAGGTAATATGACCTTCCCAACGACCACGCAGCTGACCAATTTCTTC**<br>**TTGTGCACGACGGCTTTCGGTAACAATCAGACGTGCATGTTTCATAAATGCCTGACCA**<br>**AAGCTGGTCAGGCTAACACCACGTTTGGTACGAACCAGCAGCGGTGCTTTCAGTTCA**<br>**TCTTCCAGCTGCTGAATTGCTGCGCTCAGTGCAGGCTGGCTCAGATGCAGCAGCTGTG**<br>**CTGCTGCACGCAGGCTACCAACCTCTTCAATACAAATCAGTGCCTGCAGGGTCTGCAG**<br>**TTTCAT**AGAGACCTCCTTGGACCTGGTACGTCGCTAGCGGGTAGTGTGACGTGGCGCAGGT<br>BsaI restriction site BsaI recognition sequence GoldenRBS PaqCI restriction site PaqCI recognition sequence<br>GATGGACTTCATGCTGAC |
| P$_{tsaM/tsaR}$ | GATGATGGCGAGCATTCACCTGCGGTACTTACGGTCTCATCAT**GGCAATGTCCCGGTTCAGT**<br>PaqCI recognition sequence PaqCI restriction site BsaI recognition sequence BsaI restriction site<br>**GATTGGAAAAACCGATTATATCGCCGAAAAAACGATTTGTTGTGTTCAGTGCTGCGTC**<br>P$_{tsaM/tsaR}$<br>**ACTAGCATCCCCCAAAACCCCAACAAGGAGACACTGC**ATGGTGGCGCAGGTGATGGACT<br>PaqCI restriction site PaqCI recognition sequence<br>TCATGCTGAC |

**TtuA:L-tartrate**

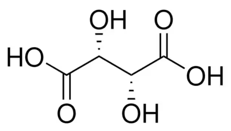


Ligand: L-tartrate
Source: Sigma- Aldrich T109
CAS ID: 87-69-4
Stock: 200 g/L
Solvent: $H_2O$
Storage: 25 °C (2 months)

Response NatTtuA and SynTtuA

**No data available**

Metadata

| | |
|---|---|
| Host | *Escherichia coli TOP10 (F- mcrA Δ(mrr-hsdRMS-mcrBC) Φ80lacZΔM15 ΔlacX74 recA1 araD139 Δ(araleu)7697 galU galK rpsL (StrR) endA1 nupG)* |
| Origin | *Agrobacterium vitis* |
| Parts | TtuA: GenBank U32375.1 |
| | $P_{ttuB/ttuA}$: Intergenic region between *ttu*A and *ttu*B*;* GenBank U32375.1 |
| Architecture | Natural and Synthetic |
| Experiment | 150 µL EZ Rich medium, 96-well black microtiter plate<br>PerkinElmer Ensight, exc. 588 nm, emm. 633 nm at 30 °C<br>Fluorescence after 24 h of growth |
| Notes | / |

#### TtuA:L-tartrate

| | |
|---|---|
| *ttu*A | GATGATGGCGAGCATTCACCTGCGGTAGCTA**CTATGCGTGTTCGAAATAACGATCAGCGT**<br>PaqCI recognition sequence PaqCI restriction site<br>**TGTCGCCCATAAGCTTGATGAGCTGATCCCGGCCTGCGTCCCTAACACCCTTCATCCA**<br>**GGCTGCCGCGAGCGGAAGCGTTTCCAGCCGGCTGTCGTCCGACGTGCGGATCGACAG**<br>**GAACTTCAGGCCTTTGACGGTCAGCTTCGCCGTCCAACGCGGTACGATCGCGGATCC**<br>**GATGCCCGCTGCGACGAGATTGATGATCGTCTGCTTCTCGTCCGCGACCTGCGCGATA**<br>**CGCGGGCGGAGCCCAGCCTTTGCGAATATGTTCATCGTAAGATCGTGACTGTGGGGA**<br>**CGGGACCGGCGATCCGGGACGATGAGCGGTTCGTCTACCAAGGCGTCCACCGTTACC**<br>**TCGCCAGCCGCAGCAAGCGGATTGGTAGATGAAACGGCGACGACCGCTGTTTCGAAA**<br>**AACAAAGGCTTCACCGAGATGGCGCGGCTCAACCCATGCGTAGGCCGGATCAGAATG**<br>*ttuA*<br>**ACGTCAAGCCGTCCGGCCAGAAGCTTAGGCACCAGCCTAATGGTCTTATCTTCGACG**<br>**ATCTGGACGATCGTGCCCGGAGAGACTTCTCGAAAATCCTGTAGAAGCTGCGGGACC**<br>**AGGCCTGCCGCCGCGCTGTCTATCGTTCCAATGCGAAGGAACTGGGAATGCTCATGA**<br>**CTTGCGCCGCGAAACCGATCCGCGAGCTTCTCAAGACGGTCCAGGATTGGCTTGACC**<br>**TCCTCAAGGAGCAAAACACCATCCTGCGTGAGATTAACGCTGCGCGTCGTCCGCACG**<br>**AACAGCTGCGTCCCGAGCGACTCTTCAAGAAGACGGACATGACGACCGAGGGACGCA**<br>**GGTAAAATCCCGGCTTTTTGGGCCGCTCTCCCGAAGTGCAATTCGTCGGCAACAACAA**<br>**CGAAACACCTTAGCTGCTCAAGCTCCAT**AGAGACCTCCTTGGACCTGGTACGTCGCTAGC<br>BsaI restriction site BsaI recognition sequence GoldenRBS<br>GGGTAGTGTGACGTGGCGCAGGTGATGGACTTCATGCTGAC<br>PaqCI restriction site PaqCI recognition sequence |
| $P_{ttuB/ttuA}$ | GATGATGGCGAGCATTCACCTGCGGTACTTACGGTCTCTCCAT**CGAAACAGTCCATTATTTC**<br>PaqCI recognition sequence PaqCI restriction site BsaI recognition sequence BsaI restriction site<br>**ATTTAATTATACAATTATATGAGTGTTGGCTCTGAGGAAAGGGGATAATAGGGTCGCC**<br>$P_{ttuB/ttuA}$<br>**CGATGCCGCTGGAGGGCGGAAGACTTACATCTGGGAGGAGCTCGCT**ATGGTGGCGCAG<br>PaqCI restriction site<br>GTGATGGACTTCATGCTGAC<br>PaqCI recognition sequence |
