## Supplementary File 2 for "Systematic mapping of orthogonality and domain-swap permissiveness across LysR-type transcriptional biosensors"

### Overview

**Supplementary File 1:** Regulator identity sheets (provided as a **separate file**).

---

**Supplementary Figure S1:** Non-responsive LTTR biosensors.

**Supplementary Figure S2:** Growth curves of the functional biosensors.

**Supplementary Figure S3:** Orthogonality fluorescence output.

**Supplementary Figure S4:** Growth curves of the orthogonality strains.

**Supplementary Figure S5:** Per-design domain-swap growth curves.

---

**Supplementary Table S1:** Ligand concentrations (orthogonality and domain-swapping experiments).

**Supplementary Table S2:** Primer list.

**Supplementary Table S3:** Plasmids and strains used in this work.

**Supplementary Table S4:** Statistical analysis — functionality assay.

**Supplementary Table S5:** Statistical analysis — orthogonality assay.

**Supplementary Table S6:** Statistical analysis — domain-swapping analysis.

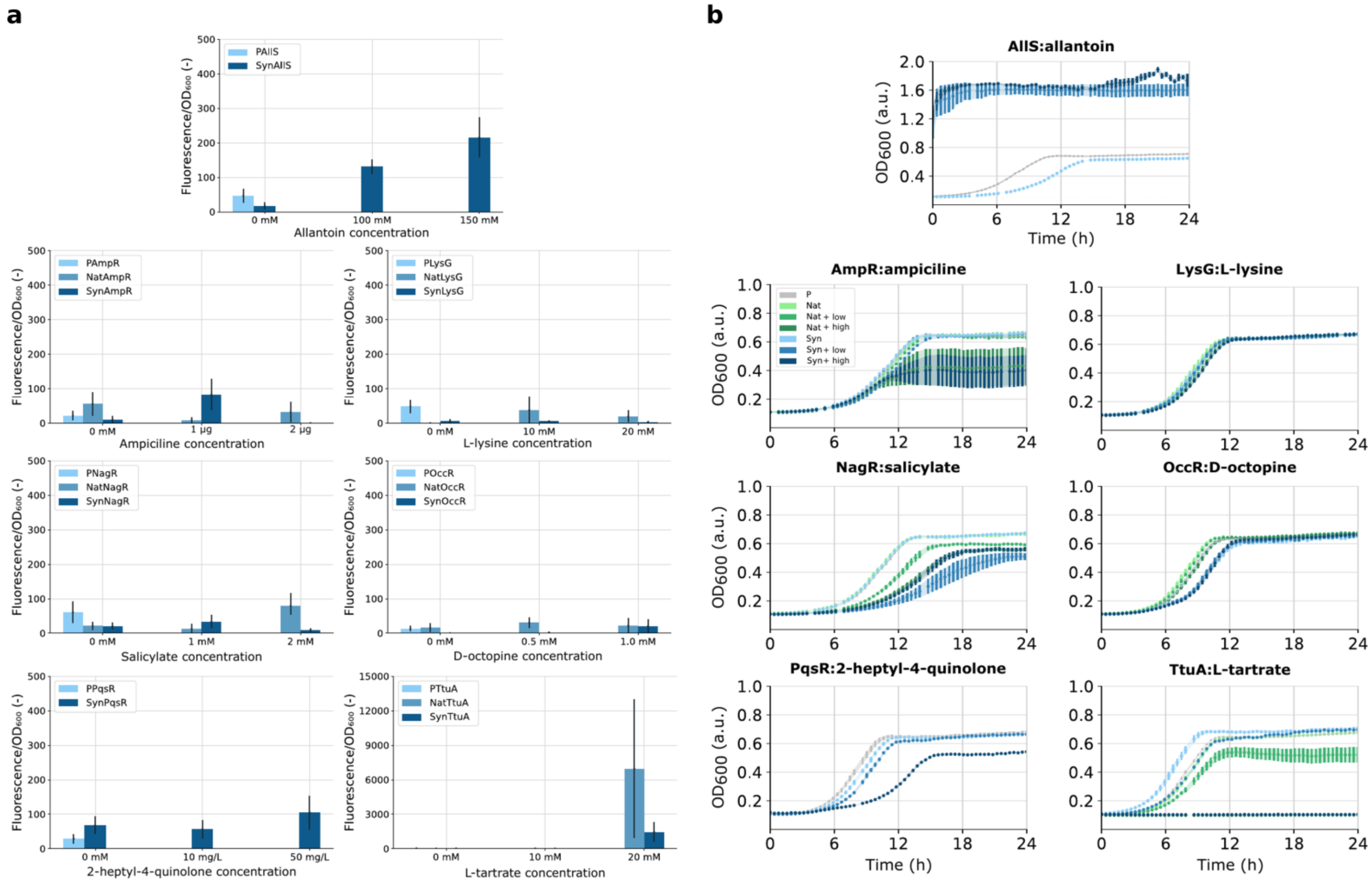


**Supplementary Figure S1 | Non-responsive LTTR biosensors.** Endpoint fluorescence **(a)** and growth **(b)** of the seven library regulators that showed no statistically significant induction in the natural-diversity screen (AllS, AmpR, LysG, NagR, OccR, PqsR and TtuA). **(a)** Corrected, OD600-normalised fluorescence output after 24 h of growth; per regulator the promoter-only control (P) is shown in light blue, the natural-architecture circuit (Nat) in blue and the synthetic-architecture circuit (Syn) in dark blue, each tested at two ligand concentrations. The effect of transcription-factor expression on promoter output was tested with Welch's t-tests and the effect of ligand addition with one-way ANOVA (alpha = 0.05); all comparisons shown here are non-significant **(Supplementary Table S4)**. **(b)** OD600 growth curves of the same strains, with the promoter control in grey, the natural circuit in green and the synthetic circuit in blue (lighter and darker shades denote the low and high ligand concentrations); the ligand of AllS, allantoin, interfered with the optical-density measurement. Bars/curves show the mean of four biological replicates (n = 4); error bars, standard error. OD600, optical density at 600 nm; a.u., arbitrary units; P, promoter circuit; Nat, natural biosensor; Syn, synthetic biosensor.

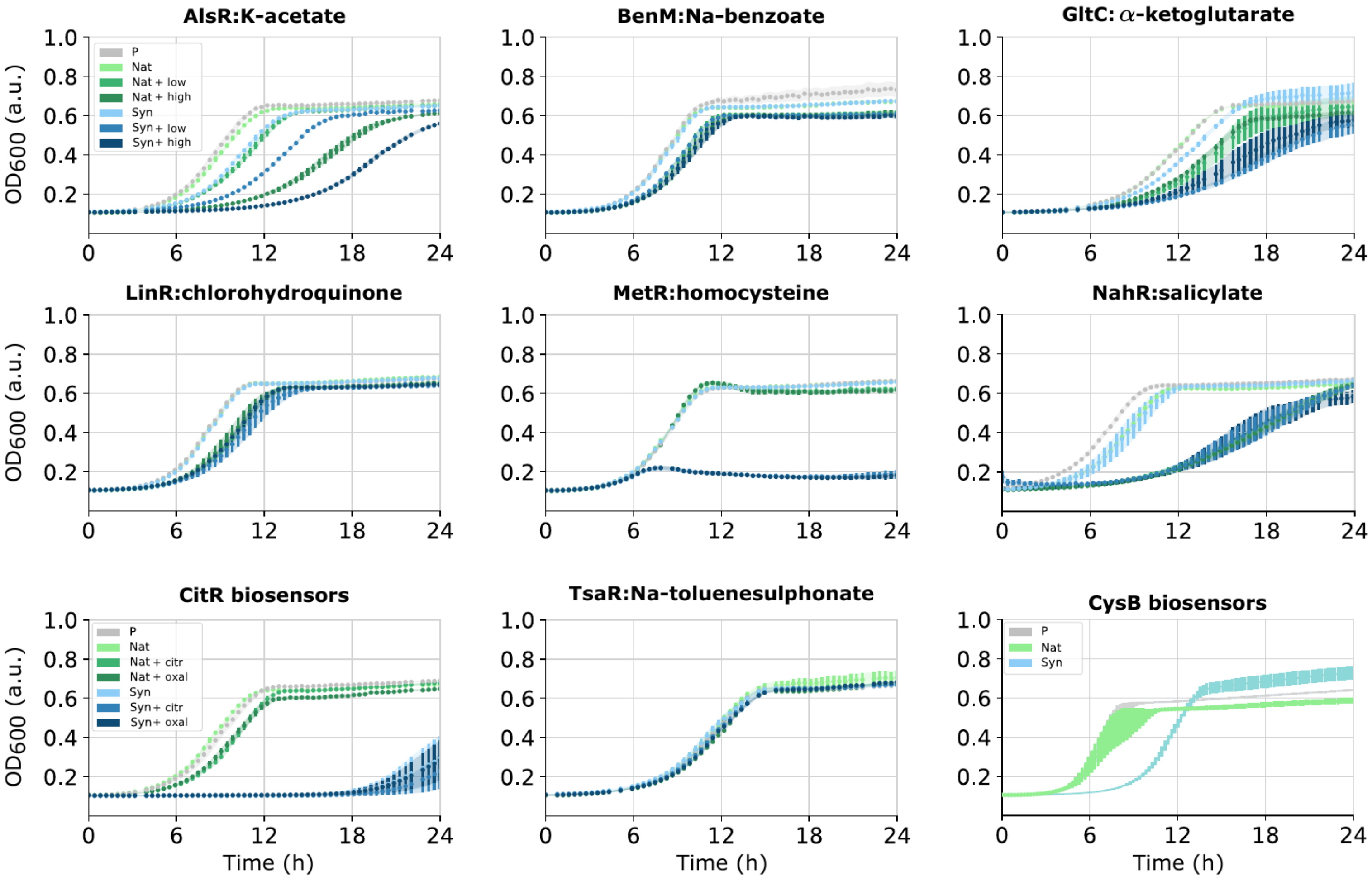


**Supplementary Figure S2 | Growth curves of the functional biosensors.** OD600 growth curves over 24 h for the nine functional biosensor strains (AlsR, BenM, GltC, LinR, MetR, NahR, CitR, TsaR and CysB). Per regulator the promoter-only control (P) is shown in grey, the natural-architecture circuit (Nat) in green and the synthetic-architecture circuit (Syn) in blue, with lighter and darker shades denoting the low and high ligand concentrations tested in the functionality experiment (Fig. 2, for ligand concentrations see Supplementary Table S1). For CitR, two candidate ligands were tested (citr, Na-citrate; oxal, oxaloacetic acid); CysB was not tested with inducing ligand. The curves confirm that the fluorescence differences in Fig. 2 are not driven by growth defects, and show the reduced growth of SynMetR and of the SynCitR strain at high ligand. Mean of four biological replicates (n = 4); error bars, standard error. OD600, optical density at 600 nm; a.u., arbitrary units; P, promoter circuit; Nat, natural biosensor; Syn, synthetic biosensor.

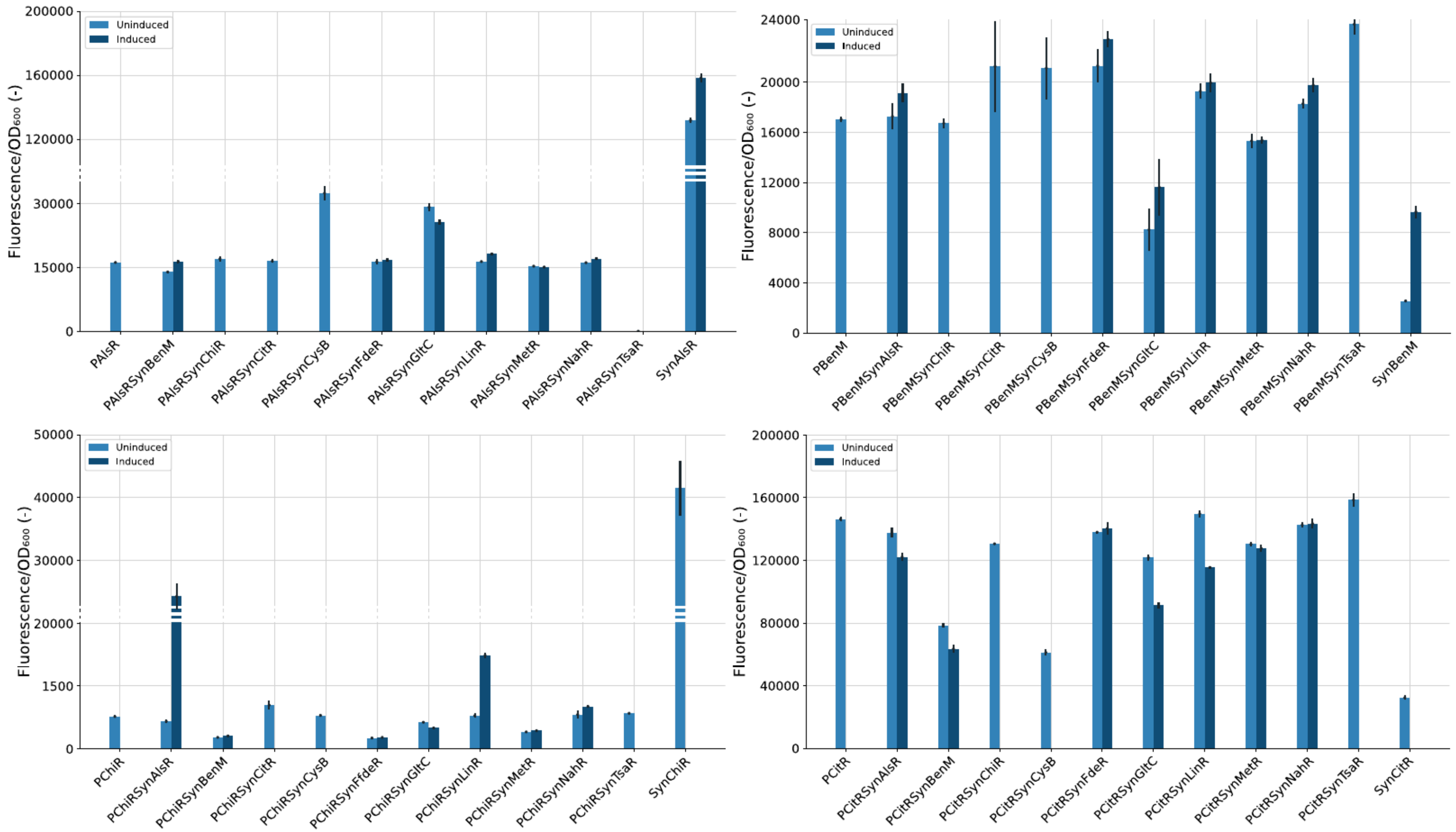


**Supplementary Figure S3 | Orthogonality fluorescence output.** Corrected, OD600-normalised fluorescence output of all orthogonality strains, grouped per target promoter. Orthogonality circuits were built in the MoBioS platform by combining the promoter of one regulator with the transcription factor of another (PTF-A SynTF-B); each panel shows one promoter together with its promoter-only control and the relevant synthetic biosensor reference. Bars give the uninduced (light blue) and induced (dark blue) output after 24 h of growth, and are the data underlying the orthogonality matrix in Fig. 3 (for ligand concentrations, see Supplementary Table S1). The effects of transcription-factor expression and of ligand addition were tested by Welch's t-tests (Supplementary Table S5). Mean of four biological replicates (n = 4); error bars, standard error. OD600, optical density at 600 nm. **(Part 1)**.

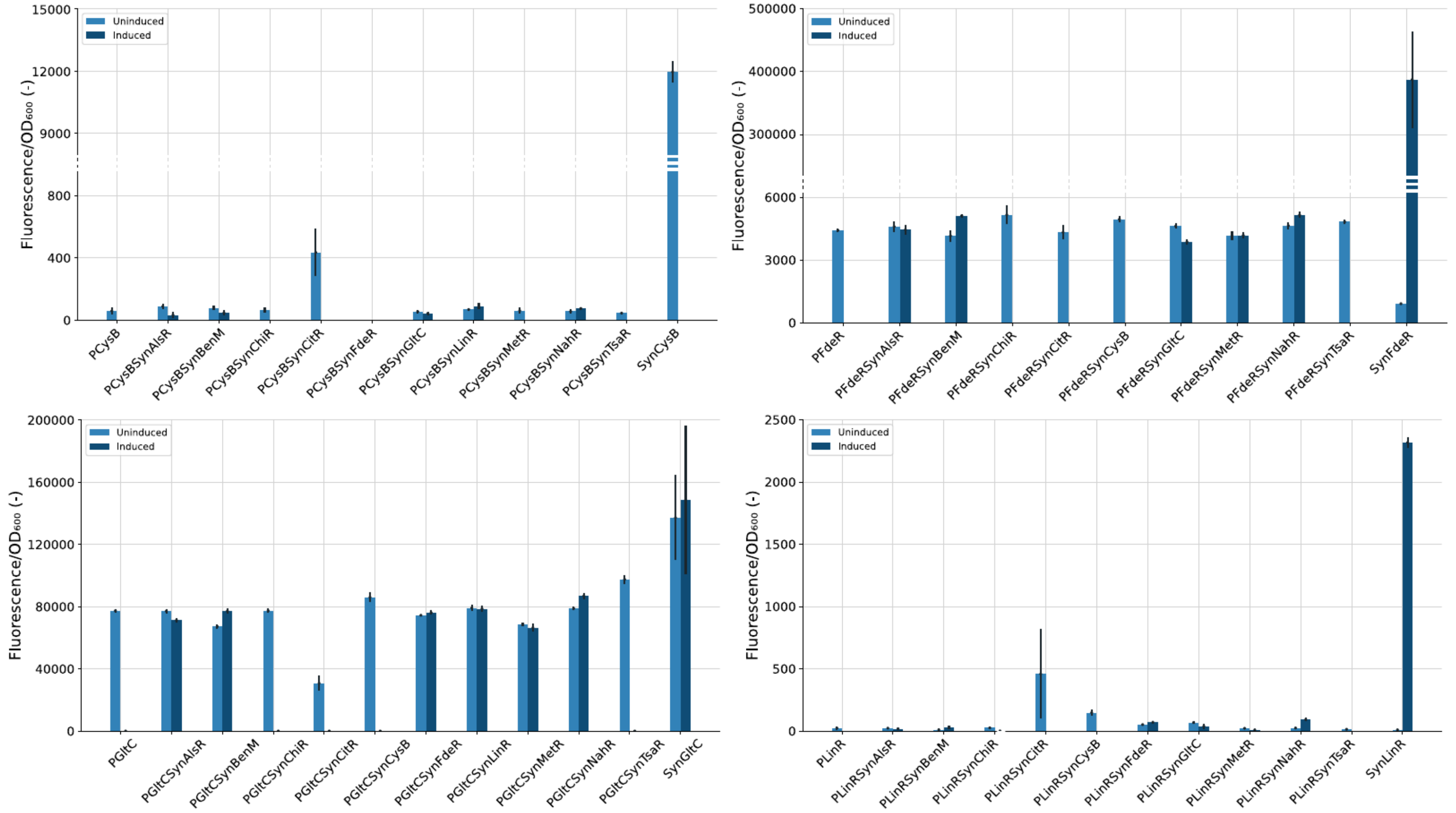


**Supplementary Figure S3 (continued).** Part 2.

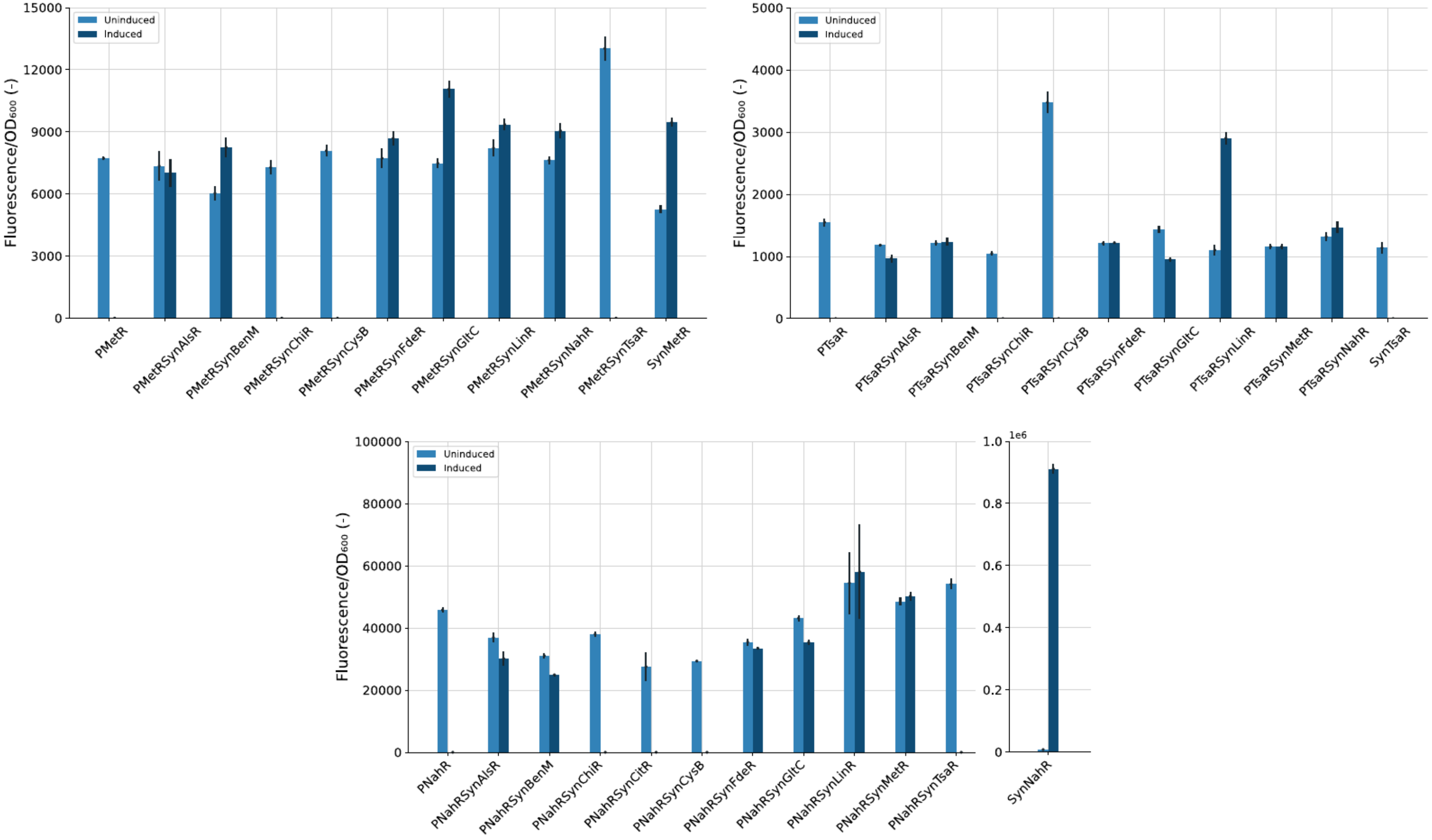


**Supplementary Figure S3 (continued).** Part 3.

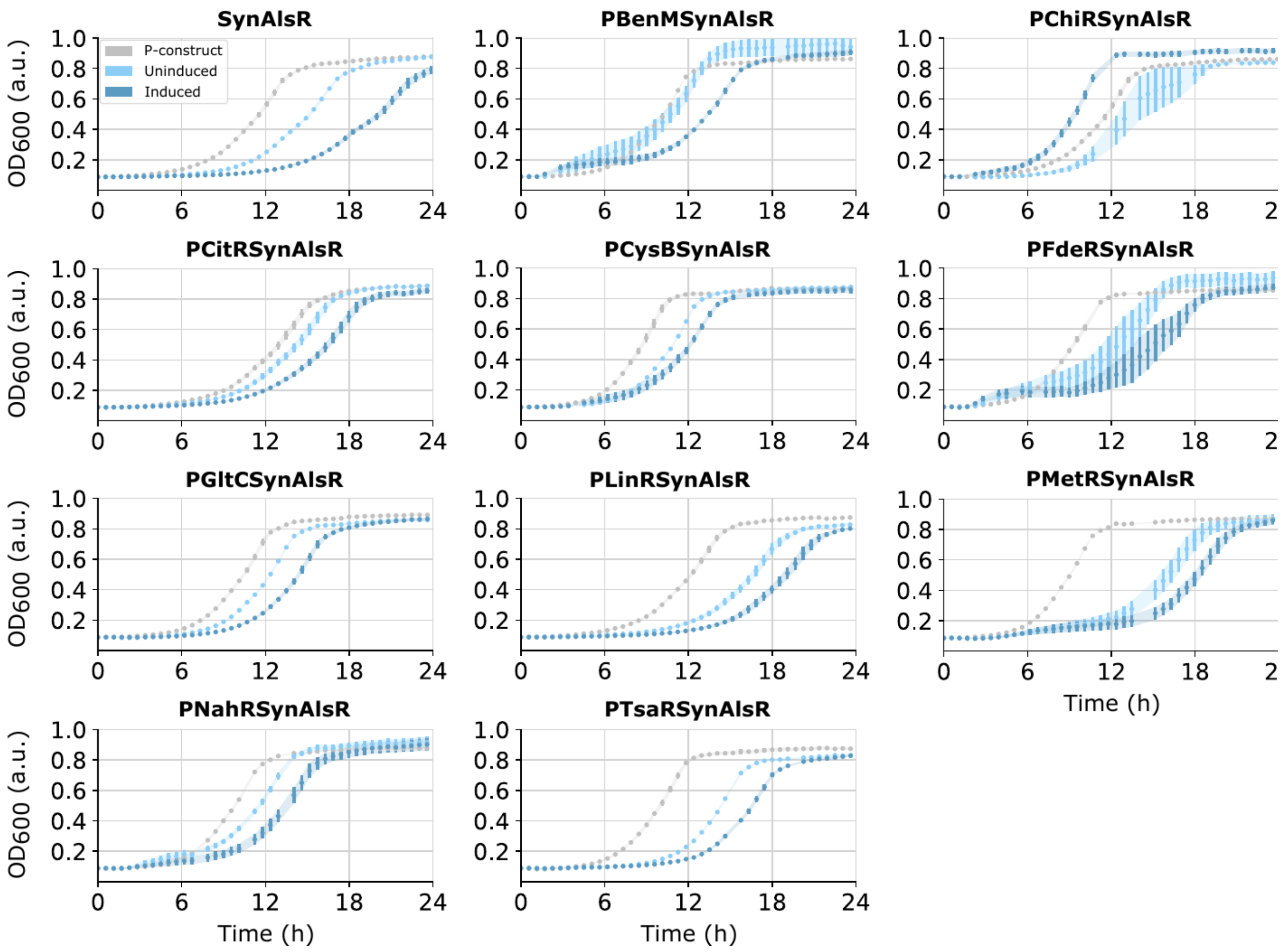


**Supplementary Figure S4 | Growth curves of the orthogonality strains.** OD600 growth curves over 24 h for the orthogonality strains, grouped per transcription factor. Each panel set shows one regulator's synthetic biosensor reference (SynTF) together with every promoter-X + SynTF combination; curves are shown for the promoter-only construct (grey) and for the combination strain in the absence (light blue, uninduced) and, where a ligand was available, presence (dark blue, induced) of the regulator's ligand (for ligand concentrations, see Supplementary Table S1). These curves support the reduced-growth annotations in Fig. 3 and the exclusion of the CitR-expressing strains, which were severely growth-impaired. Mean of four biological replicates (n = 4); error bars, standard error. OD600, optical density at 600 nm; a.u., arbitrary units. (AlsR).

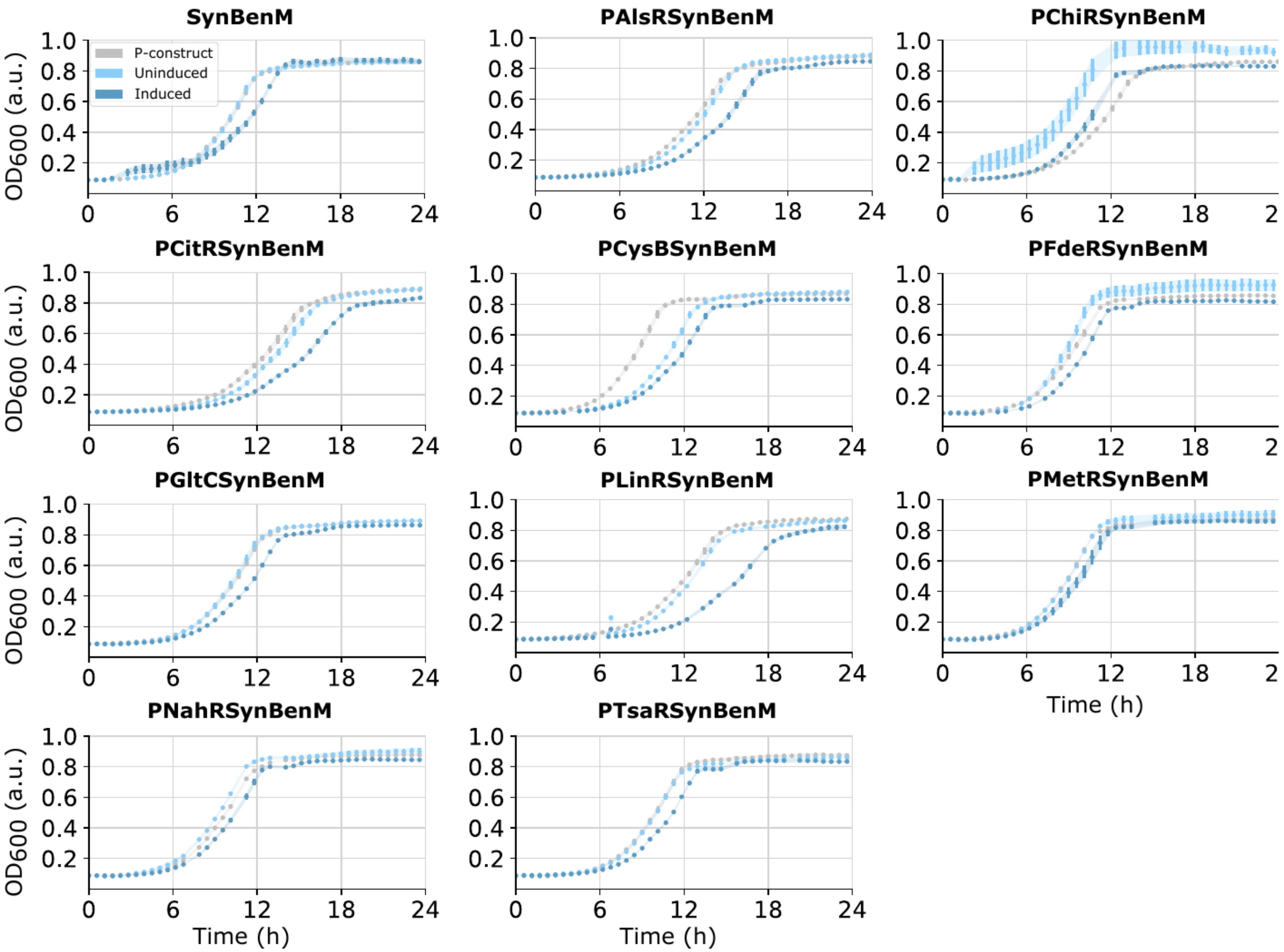


**Supplementary Figure S4 (continued).** BenM.

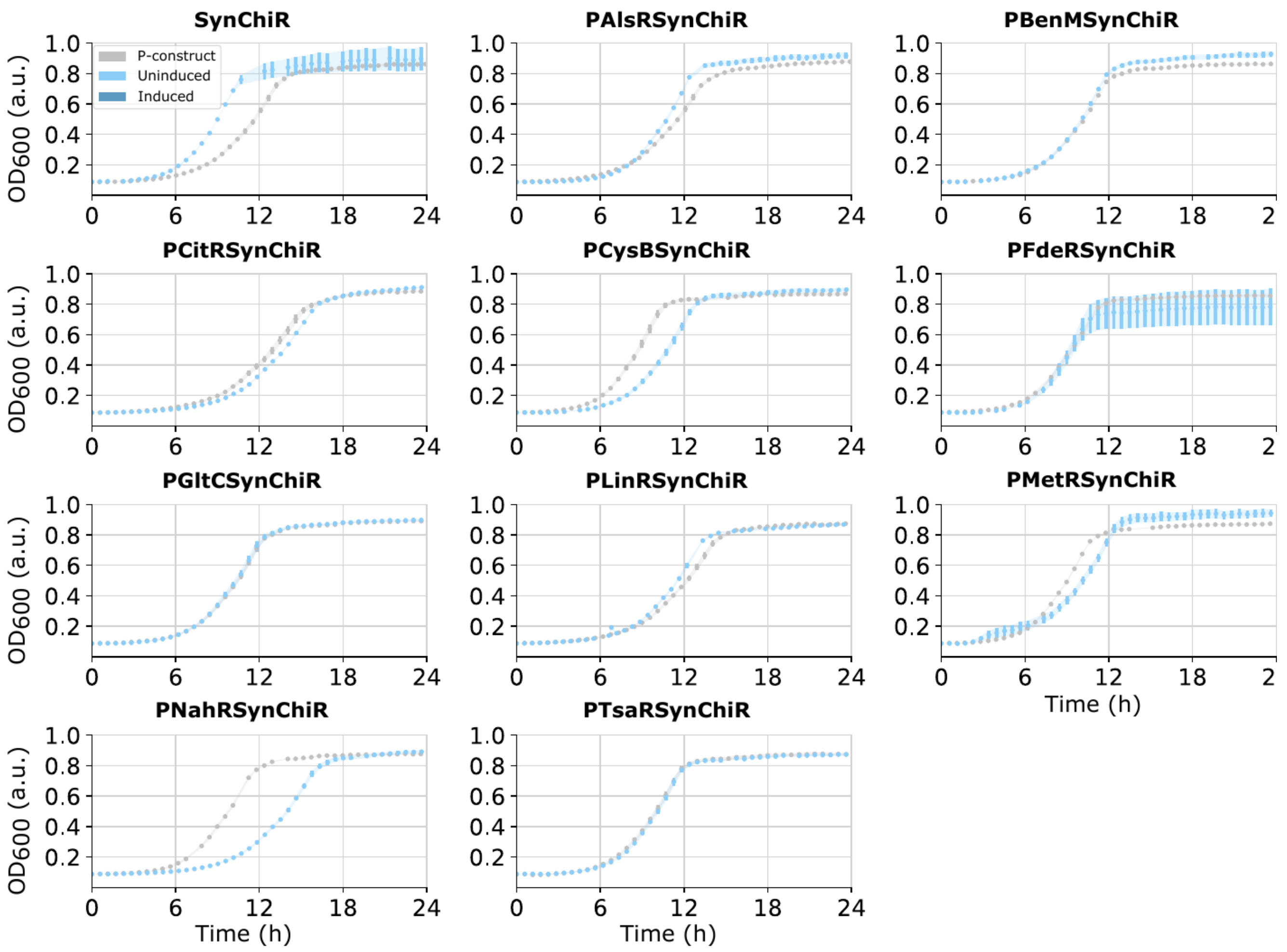


**Supplementary Figure S4 (continued).** ChiR.

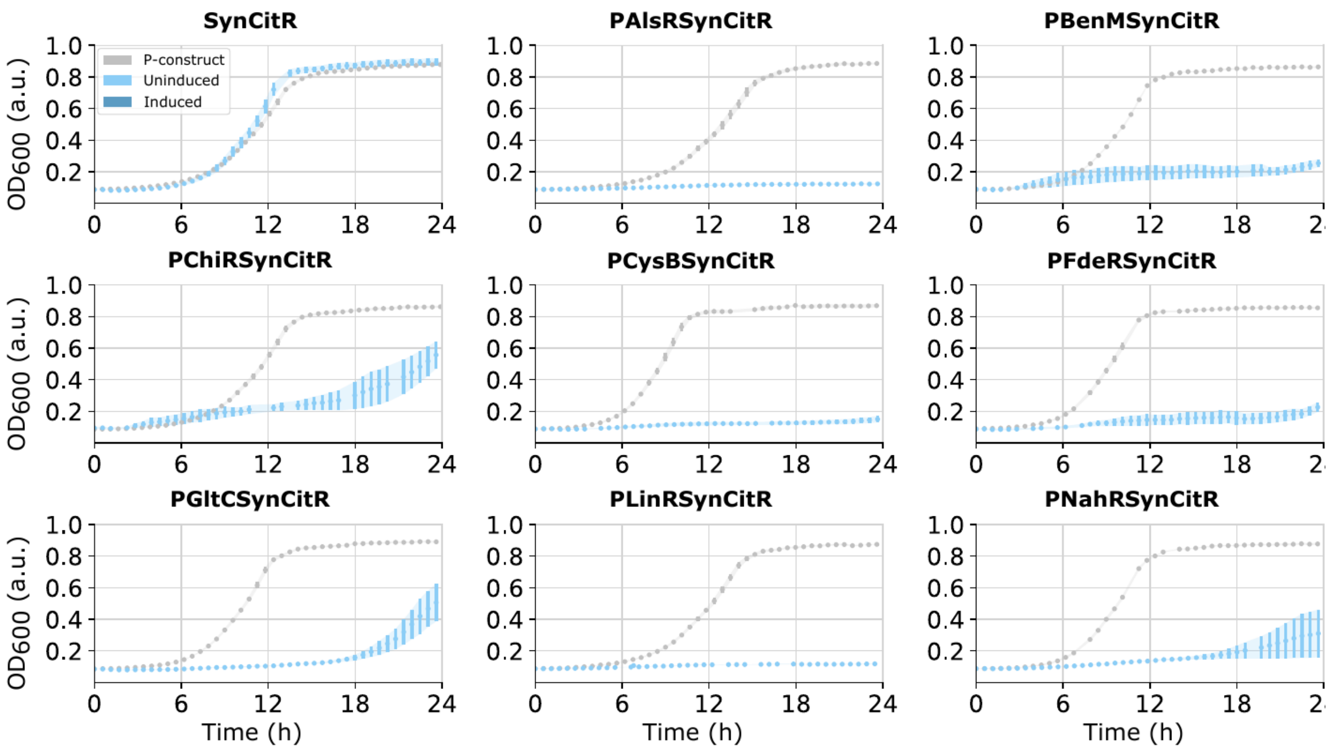


**Supplementary Figure S4 (continued).** CitR.

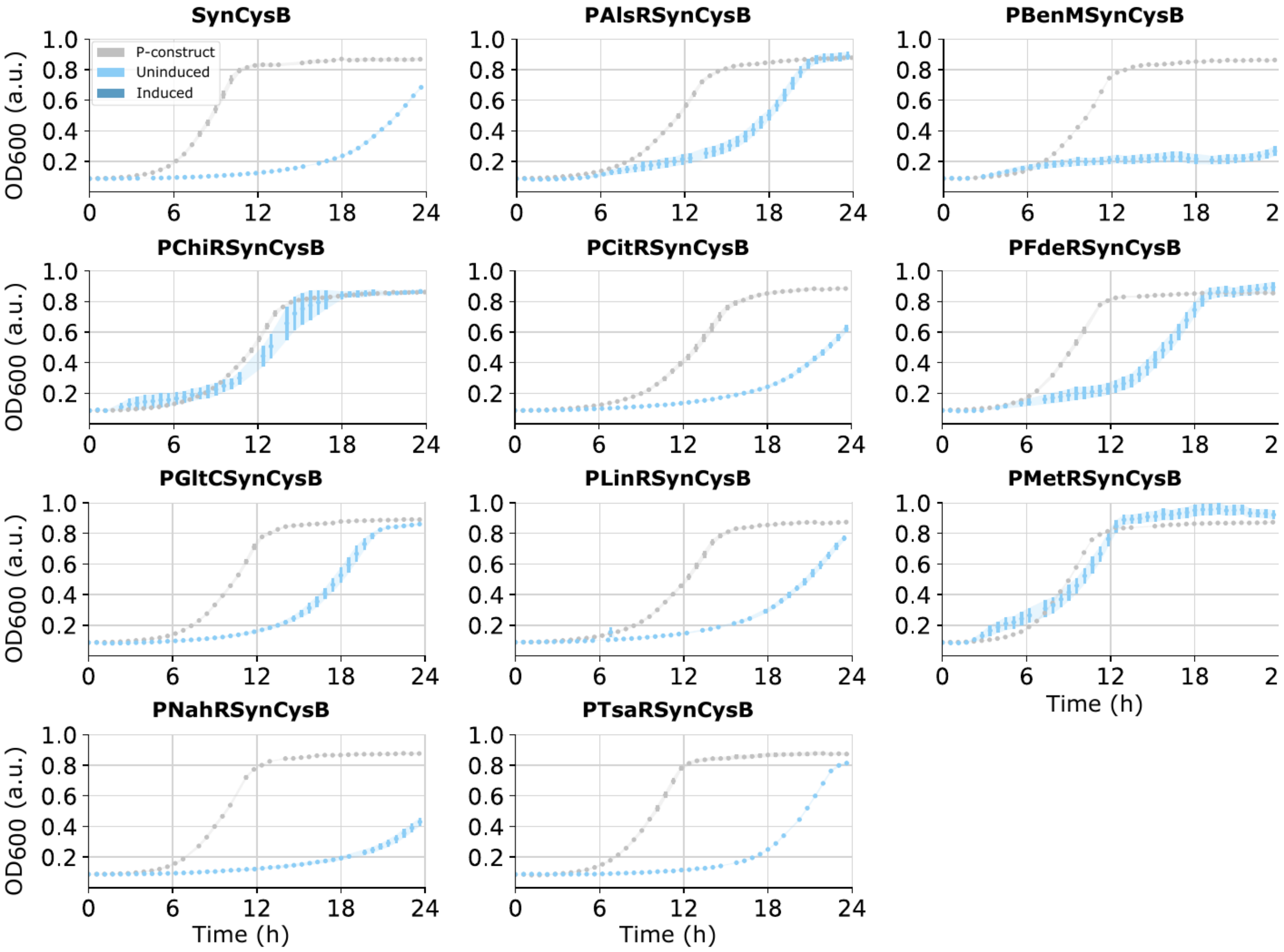


**Supplementary Figure S4 (continued).** CysB.

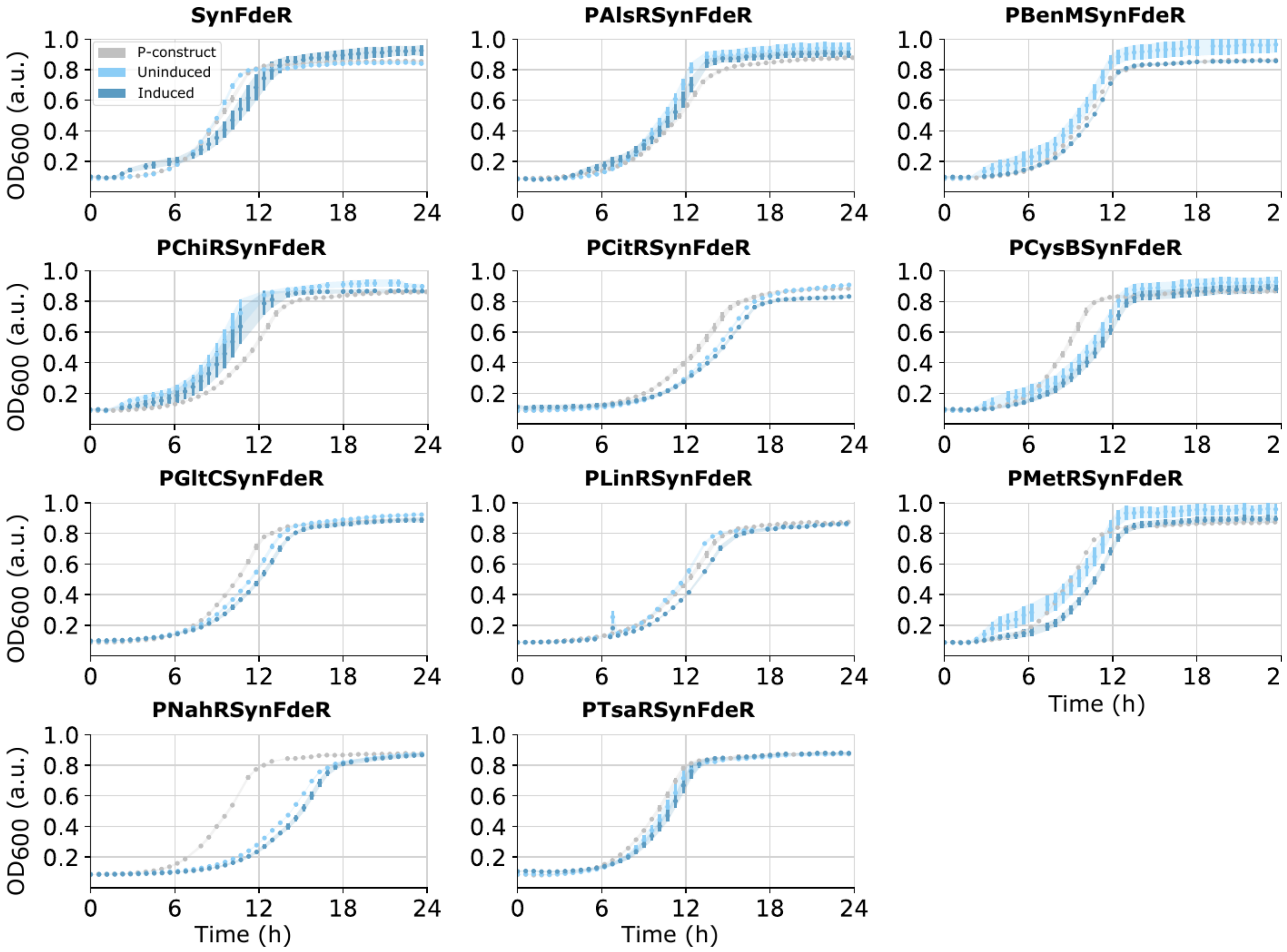


**Supplementary Figure S4 (continued).** FdeR.

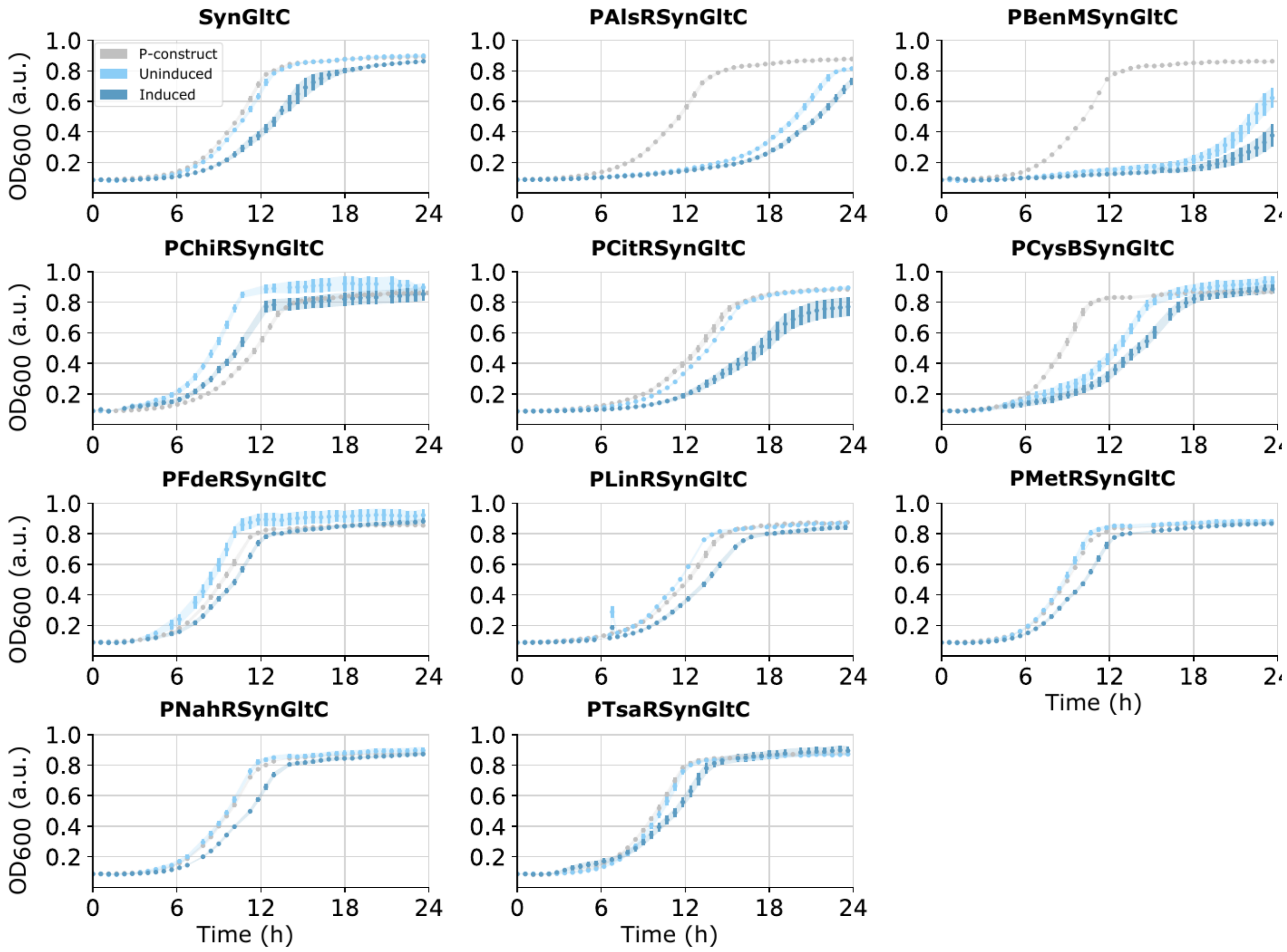


**Supplementary Figure S4 (continued).** GltC.

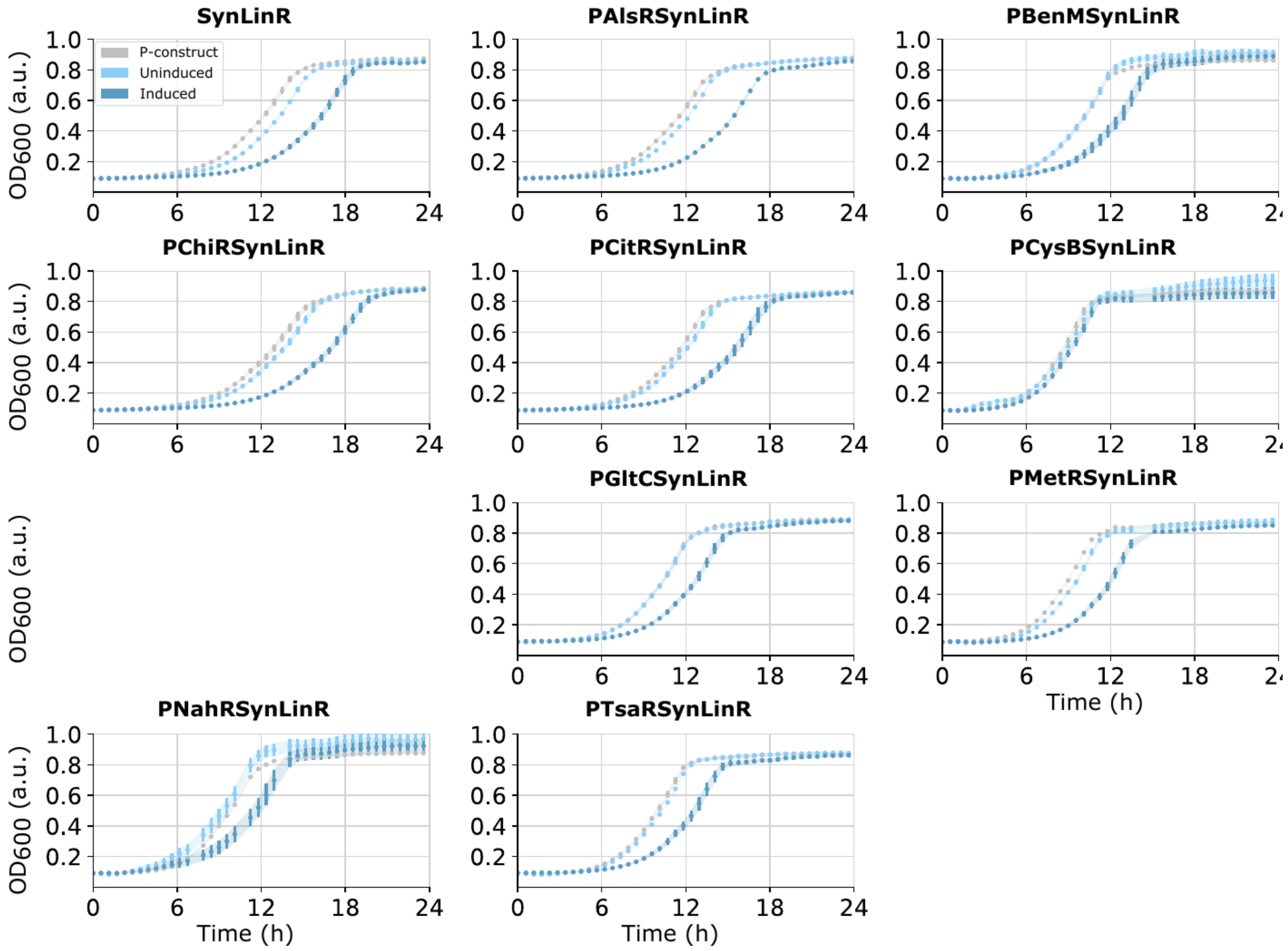


**Supplementary Figure S4 (continued).** LinR.

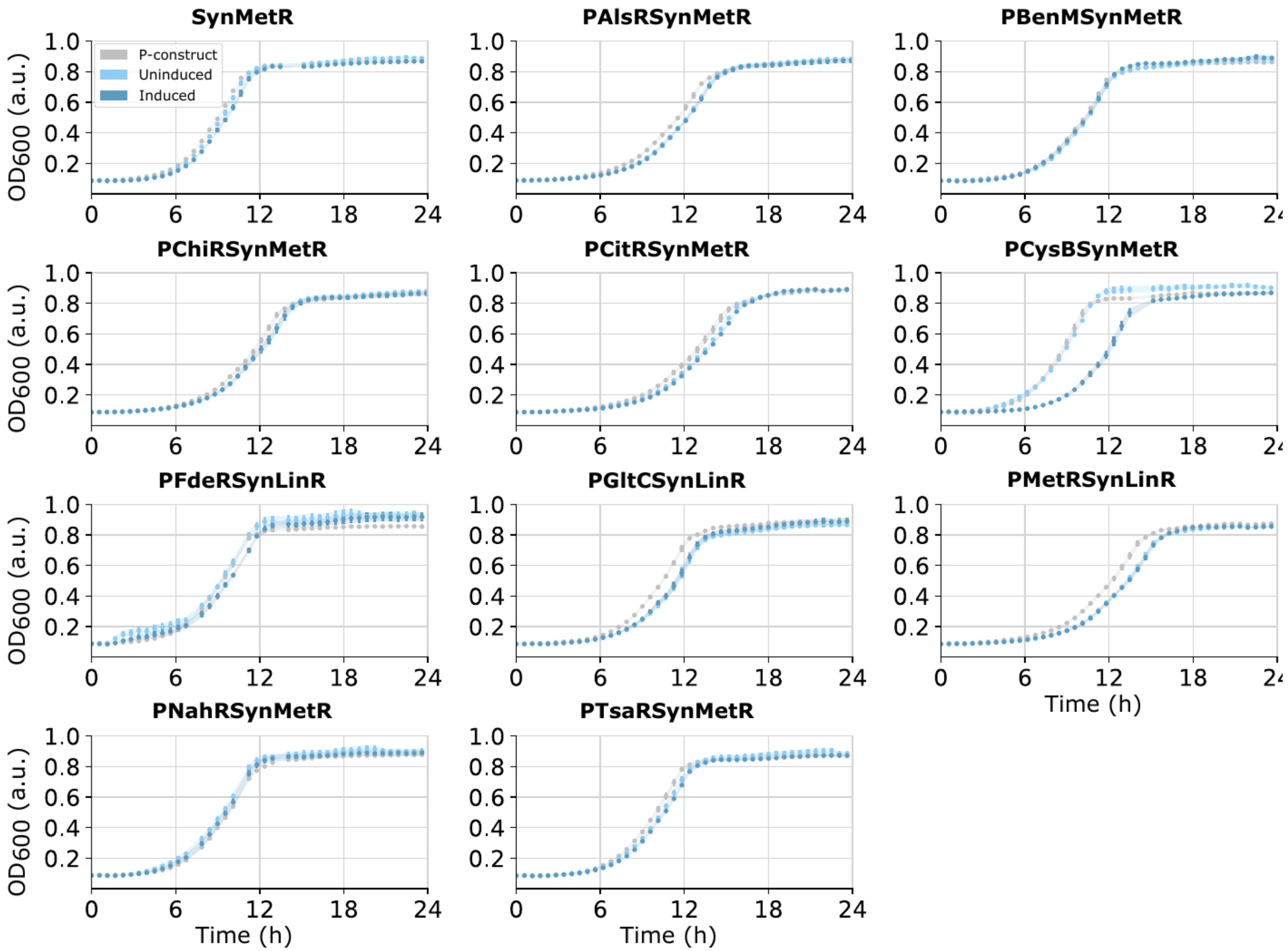


**Supplementary Figure S4 (continued).** MetR.

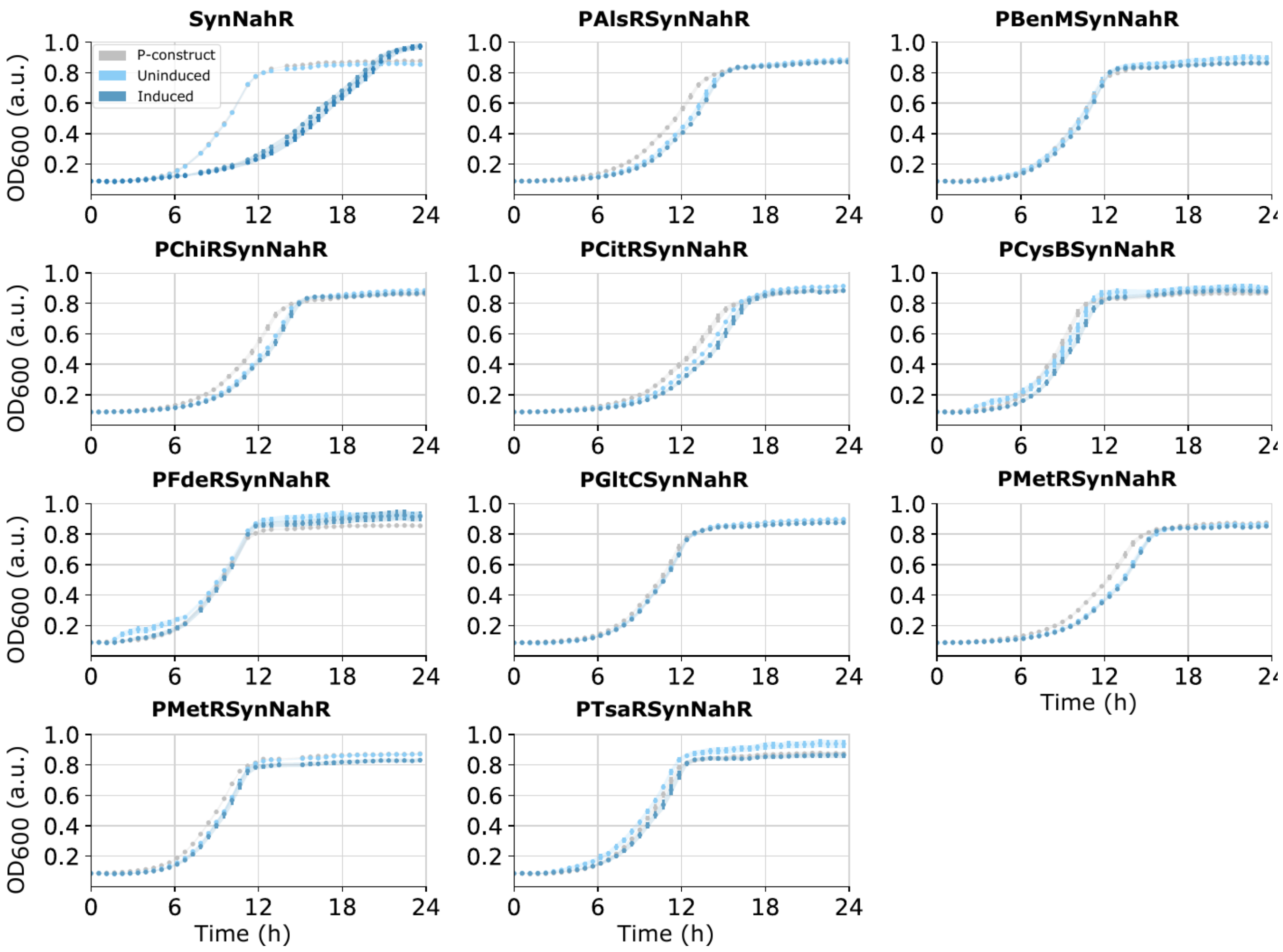


**Supplementary Figure S4 (continued).** NahR.

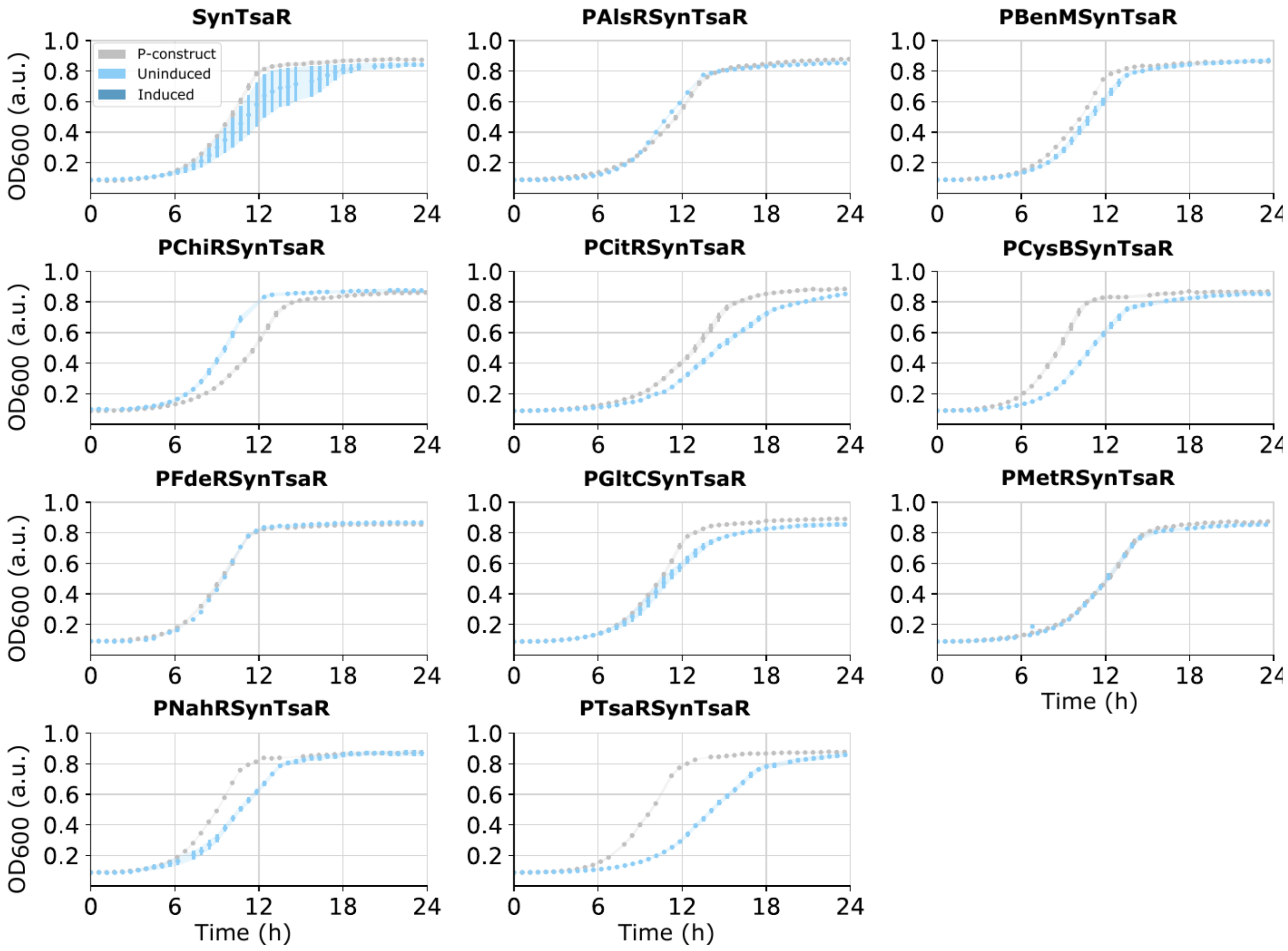


**Supplementary Figure S4 (continued).** TsaR.

### A) DBD design

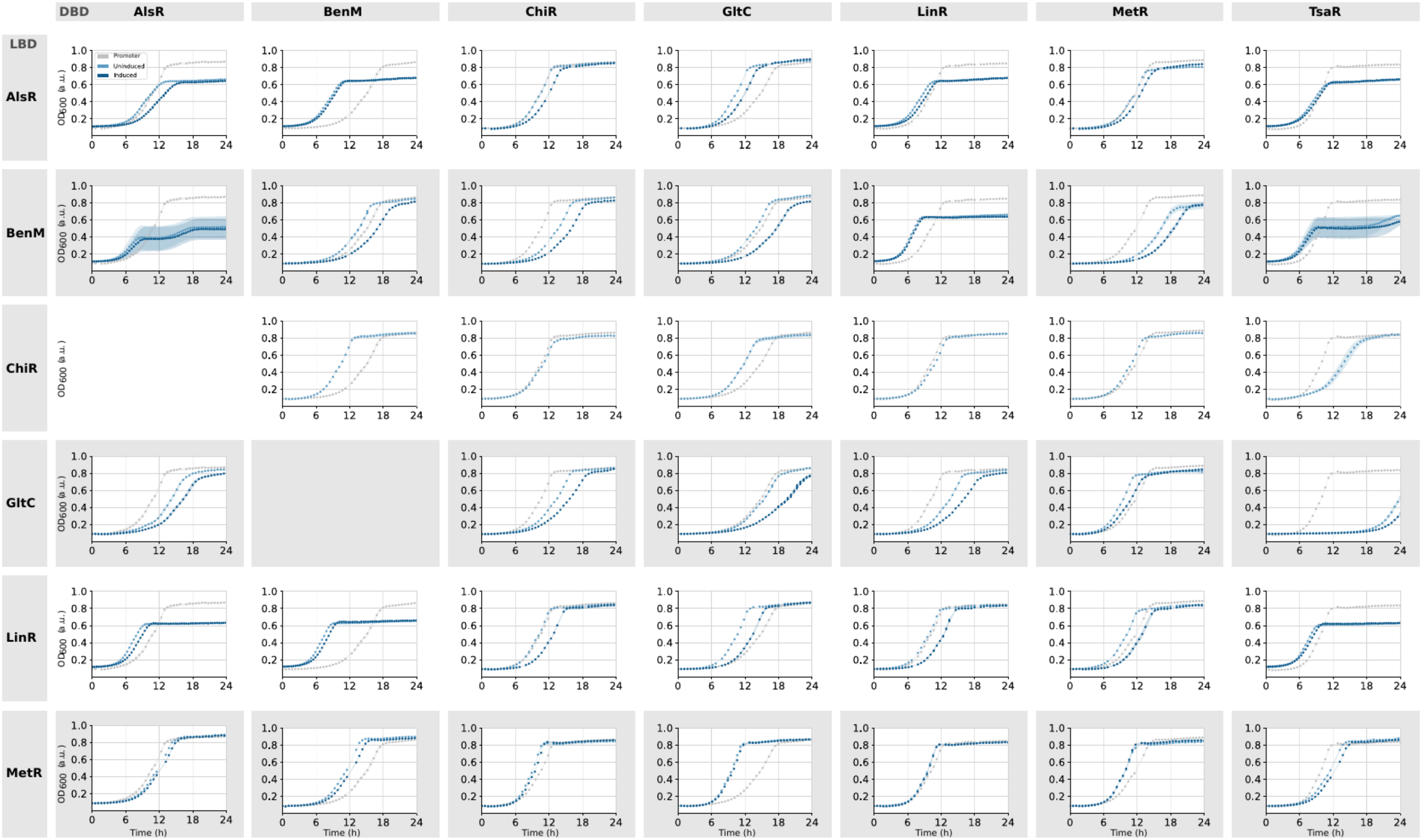


**Supplementary Figure S5 | Per-design domain-swap growth curves.** OD600 growth curves over 24 h for the domain-swapped chimera strains, shown for the three domain-swapping designs — **(a)** DBD, **(b)** DBD+LH and **(c)** DBD+LH+H — with panels arranged by DNA-binding-domain donor (columns) and ligand-binding-domain donor (rows). Each panel shows the target promoter control (grey) and the chimera in the absence (light blue, uninduced) and presence (dark blue, induced) of externally added ligand; ligand concentrations are given in Supplementary Table S1. Domain-swap circuits that gave non-viable strains are left blank, and the parental 'Syn' biosensors are shown on the diagonal as references. Mean of four biological replicates (n = 4); error bars, standard error. OD600, optical density at 600 nm; a.u., arbitrary units; DBD, DNA-binding domain; LBD, ligand-binding domain; LH, linker helix; H, hinge. ((a) DBD design).

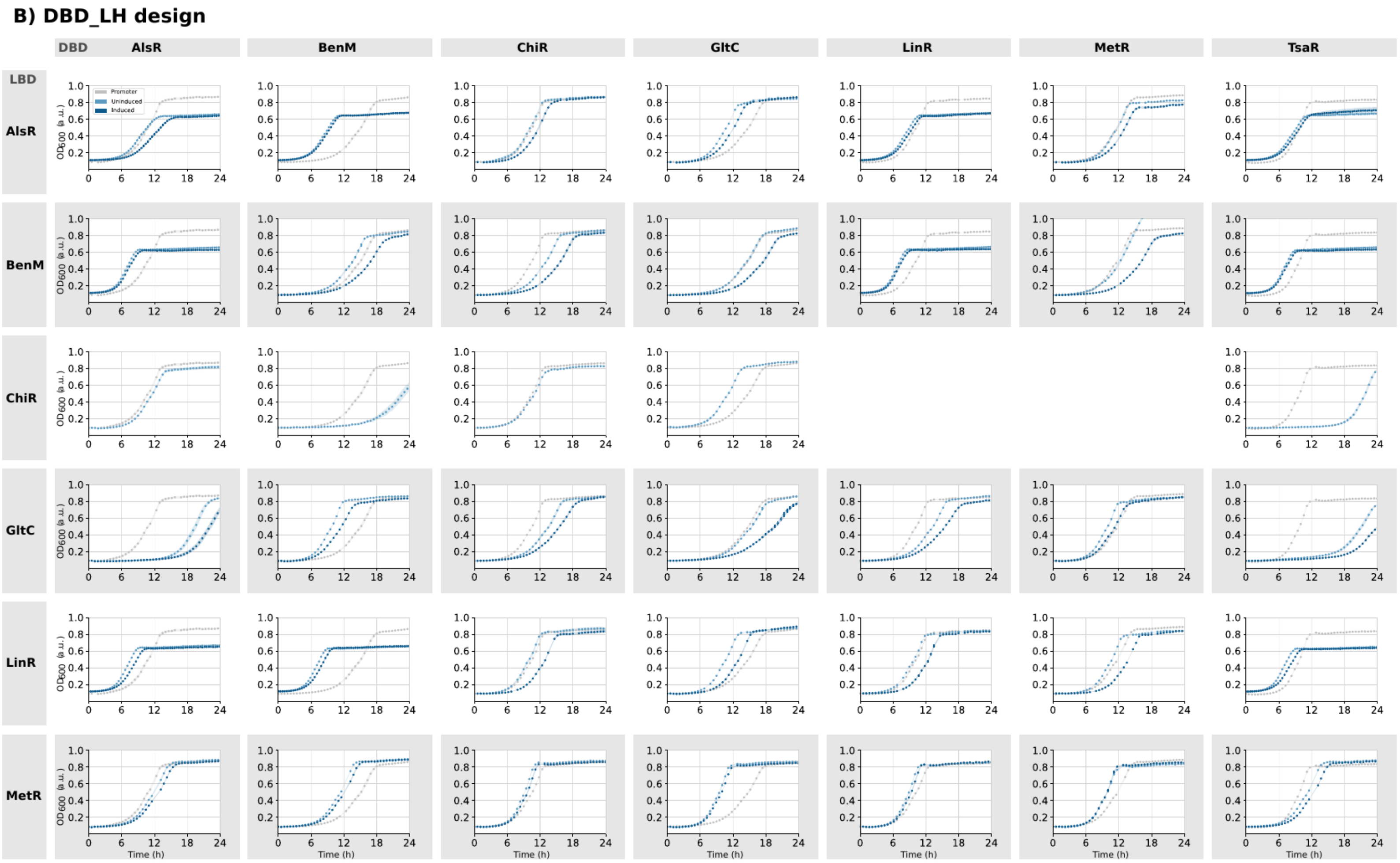


**Supplementary Figure S5 (continued).** (b) DBD+LH design.

### C) DBD_LH_H design

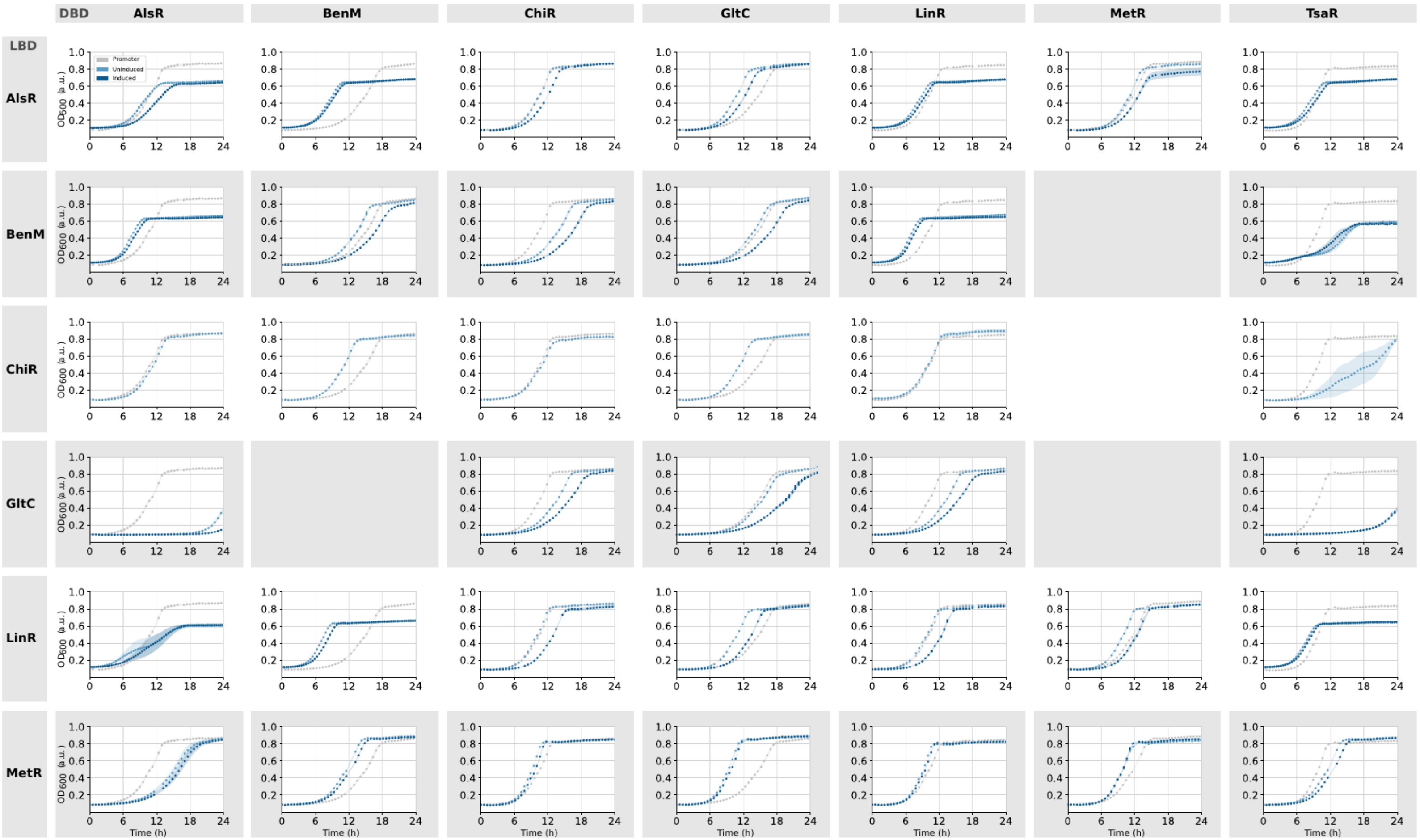


**Supplementary Figure S5 (continued).** (c) DBD+LH+H design.

**Supplementary Table S1 | Ligand concentrations (orthogonality and domain-swapping experiments).** Inducer concentrations used in the orthogonality and domain-swapping induction experiments; functionality-assay ligand preparations are given per regulator in Supplementary File 1. N.a. = not applicable.

| **Transcription factor** | **Ligand** | **Concentration orthogonality experiment** | **Concentration domain swapping experiment** |
|---|---|---|---|
| AlsR | K-Acetate | 50 mM | 10 mM |
| BenM | Na-Benzoate | 6 mM | 4 mM |
| FdeR | Naringenin | 0.2 mM | n.a. |
| GltC | α-Ketoglutarate | 6 mM | 4.5 mM |
| LinR | Chlorohydroquinone | 0.75 mM | 0.5 mM |
| MetR | L-Homocysteine | 0.6 mM | 0.1 mM |
| NahR | Salicylate | 0.2 mM | n.a. |

**Supplementary Table S2 | Primer list.** Oligonucleotide primers used to construct the domain-swap plasmids (S2a) and the orthogonality case-study plasmids (S2b). All primers given 5'-3'.

#### S2a — Domain-swapping primers

| DBD-part | Primer | LBD-part | Primer |
|---|---|---|---|
| AlsR$_{\text{DBD(B)}}$ | TAAAAAAAGTGACCCTCAGGGGTCAGTTCAACAAAACGTTTGGTACGTTTCAGC | BenM$_{\text{LBD_H_LH(A)}}$ | AACGTTTTGTTGAACTGACCCCTGAGGGTCACTTTTTTTATCAGTACGCCATC |
| AlsR$_{\text{DBD_LH(B)}}$ | ATGGTTTTTTCAACCGAGGCGGTACGCTGTGCCAGTTCAATACC | BenM$_{\text{LBD_H(A)}}$ | TTGAACTGGCACAGCGTACCGCCTCGGTTGAAAAAACCATTCG |
| AlsR$_{\text{DBD_LH_H(B)}}$ | CCCACAAATCCGATACGAATACCCTGTTCACCACGTGCGGTAC | BenM$_{\text{LBD(A)}}$ | CCGCACGTGGTGAACAGGGTATTCGTATCGGATTTGTGGG |
| AlsR$_{\text{DBD(C)}}$ | AGAAATTCCCCTTCATCCGTGGTCAGTTCAACAAAACGTTTGGTACGTTTCAGC | ChiR$_{\text{LBD_H_LH(A)}}$ | AACGTTTTGTTGAACTGACCACGGATGAAGGGGAATTTCTC |
| AlsR$_{\text{DBD_LH(C)}}$ | ATATTATTGTCCAGCTTGGGGGTACGCTGTGCCAGTTCAATACC | ChiR$_{\text{LBD_H(A)}}$ | TTGAACTGGCACAGCGTACCCCCAAGCTGGACAATAATATCCGCTTGGCGGTAG |
| AlsR$_{\text{DBD_LH_H(C)}}$ | GTGTCTACCGCCAAGCGGATACCCTGTTCACCACGTGCGGTAC | ChiR$_{\text{LBD(A)}}$ | CCGCACGTGGTGAACAGGGTATCCGCTTGGCGGTAGACAC |
| AlsR$_{\text{DBD(G)}}$ | AGAAATTCTTTACCAATCGGGGTCAGTTCAACAAAACGTTTGGTACGTTTCAGC | GltC$_{\text{LBD_H_LH(A)}}$ | AACGTTTTGTTGAACTGACCCCGATTGGTAAAGAATTTCTGATCC |
| AlsR$_{\text{DBD_LH(G)}}$ | ACGGTGCCACGATGCGGATCGGTACGCTGTGCCAGTTCAATACC | GltC$_{\text{LBD_H(A)}}$ | TTGAACTGGCACAGCGTACCGATCCGCATCGTGGCACCGTTAAAATTG |
| AlsR$_{\text{DBD_LH_H(G)}}$ | GTCGGAAAACCAATTTTAACACCCTGTTCACCACGTGCGGTAC | GltC$_{\text{LBD(A)}}$ | CCGCACGTGGTGAACAGGGTGTTAAAATTGGTTTTCCGACCAGC |
| AlsR$_{\text{DBD(L)}}$ | GCGATCGCCTCGGCACGTGGGGTCAGTTCAACAAAACGTTTGGTACGTTTCAGC | LinR$_{\text{LBD_H_LH(A)}}$ | AACGTTTTGTTGAACTGACCCCACGTGCCGAGGCGATC |
| AlsR$_{\text{DBD_LH(L)}}$ | CGCTTAAGACGGTCAGGTGAGGTACGCTGTGCCAGTTCAATACC | LinR$_{\text{LBD_H(A)}}$ | TTGAACTGGCACAGCGTACCTCACCTGACCGTCTTAAGC |
| AlsR$_{\text{DBD_LH_H(L)}}$ | CCGGCAATGATGAATTCGCGACCCTGTTCACCACGTGCGGTAC | LinR$_{\text{LBD(A)}}$ | CCGCACGTGGTGAACAGGGTCGCGAATTCATCATTGCCGGATCGGAC |
| AlsR$_{\text{DBD(M)}}$ | AGCAGAATTTCACCCTGTGCGGTCAGTTCAACAAAACGTTTGGTACGTTTCAGC | MetR$_{\text{LBD_H_LH(A)}}$ | AACGTTTTGTTGAACTGACCGCACAGGGTGAAATTCTGCTGCAGCTGTC |
| AlsR$_{\text{DBD_LH(M)}}$ | ATCTGGTGGGTCTTTTTGCAGGTACGCTGTGCCAGTTCAATACC | MetR$_{\text{LBD_H(A)}}$ | TTGAACTGGCACAGCGTACCTGCAAAAAGACCCACCAG |
| AlsR$_{\text{DBD_LH_H(M)}}$ | ATGGCCAGTTTGATAATCATACCCTGTTCACCACGTGCGGTAC | MetR$_{\text{LBD(A)}}$ | CCGCACGTGGTGAACAGGGTATGATTATCAAACTGGCCATTG |
| BenM$_{\text{DBD(A)}}$ | AGAAAAAATTTCACCGGCTGCTGTTGTCTTGACCGGTCTG | AlsR$_{\text{LBD_H_LH(B)}}$ | GCAGACCGGTCAAGACAACAGCAGCCGGTGAAATTTTTCTG |
| BenM$_{\text{DBD_LH(A)}}$ | AGACCCTGTTCACCACGTGCAATGCGCTTGGTCATGGAAAC | AlsR$_{\text{LBD_H(B)}}$ | TTTCCATGACCAAGCGCATTGCACGTGGTGAACAGGGTCTGCTGGTTATTG |
| BenM$_{\text{DBD_LH_H(A)}}$ | ACAAAACCAATAACCAGCAGGGTTTTTTCAACCGAGGCAATGCGCTTGGTC | AlsR$_{\text{LBD(B)}}$ | TTGCCTCGGTTGAAAAAACCCTGCTGGTTATTGGTTTTGTTGGTAG |
| BenM$_{\text{DBD(C)}}$ | AGAAATTCCCCTTCATCCGTTGTTGTCTTGACCGGTCTG | ChiR$_{\text{LBD_H_LH(B)}}$ | GCAGACCGGTCAAGACAACAACGGATGAAGGGGAATTTCTC |
| BenM$_{\text{DBD_LH(C)}}$ | ATATTATTGTCCAGCTTGGGAATGCGCTTGGTCATGGAAAC | ChiR$_{\text{LBD_H(B)}}$ | TTTCCATGACCAAGCGCATTCCCAAGCTGGACAATAATATCCGCTTGGCGGTAG |
| BenM$_{\text{DBD_LH_H(C)}}$ | GTGTCTACCGCCAAGCGGATGGTTTTTTCAACCGAGGCAATGCGCTTGGTC | ChiR$_{\text{LBD(B)}}$ | TTGCCTCGGTTGAAAAAACCATCCGCTTGGCGGTAGACAC |
| BenM$_{\text{DBD(G)}}$ | AGAAATTCTTTACCAATCGGTGTTGTCTTGACCGGTCTG | GltC$_{\text{LBD_H_LH(B)}}$ | GCAGACCGGTCAAGACAACACCGATTGGTAAAGAATTTCTGATCC |
| BenM$_{\text{DBD_LH(G)}}$ | ACGGTGCCACGATGCGGATCAATGCGCTTGGTCATGGAAAC | GltC$_{\text{LBD_H(B)}}$ | TTTCCATGACCAAGCGCATTGATCCGCATCGTGGCACCGTTAAAATTG |
| BenM$_{\text{DBD_LH_H(G)}}$ | GTCGGAAAACCAATTTTAACGGTTTTTTCAACCGAGGCAATGCGCTTGGTC | GltC$_{\text{LBD(B)}}$ | TTGCCTCGGTTGAAAAAACCGTTAAAATTGGTTTTCCGACCAGC |
| BenM$_{\text{DBD(L)}}$ | GCGATCGCCTCGGCACGTGGTGTTGTCTTGACCGGTCTG | LinR$_{\text{LBD_H_LH(B)}}$ | GCAGACCGGTCAAGACAACACCACGTGCCGAGGCGATC |
| BenM$_{\text{DBD_LH(L)}}$ | CGCTTAAGACGGTCAGGTGAAATGCGCTTGGTCATGGAAAC | LinR$_{\text{LBD_H(B)}}$ | TTTCCATGACCAAGCGCATTTCACCTGACCGTCTTAAGC |
| BenM$_{\text{DBD_LH_H(L)}}$ | CCGGCAATGATGAATTCGCGGGTTTTTTCAACCGAGGCAATGCGCTTGGTC | LinR$_{\text{LBD(B)}}$ | TTGCCTCGGTTGAAAAAACCCGCGAATTCATCATTGCCGGATCGGAC |
| BenM$_{\text{DBD(M)}}$ | AGCAGAATTTCACCCTGTGCTGTTGTCTTGACCGGTCTG | MetR$_{\text{LBD_H_LH(B)}}$ | GCAGACCGGTCAAGACAACAGCACAGGGTGAAATTCTGCTGCAGCTGTC |
| BenM$_{\text{DBD_LH(M)}}$ | ATCTGGTGGGTCTTTTTGCAAATGCGCTTGGTCATGGAAAC | MetR$_{\text{LBD_H(B)}}$ | TTTCCATGACCAAGCGCATTTGCAAAAAGACCCACCAG |
| BenM$_{\text{DBD_LH_H(M)}}$ | ATGGCCAGTTTGATAATCATGGTTTTTTCAACCGAGGCAATGCGCTTGGTC | MetR$_{\text{LBD(B)}}$ | TTGCCTCGGTTGAAAAAACCATGATTATCAAACTGGCCATTG |
| ChiR$_{\text{DBD(A)}}$ | AGAAAAAATTTCACCGGCTGCCAGGGTAATCATGGGGCCATTG | AlsR$_{\text{LBD_H_LH(C)}}$ | ATGGCCCCATGATTACCCTGGCAGCCGGTGAAATTTTTCTG |

| | | | |
|---|---|---|---|
| ChiR$_{\text{DBD_LH(A)}}$ | AGACCCTGTTCACCACGTGCAATGTTGCGCACCCGGCTTTC | AlsR$_{\text{LBD_H(C)}}$ | AAAGCCGGGTGCGCAACATTGCACGTGGTGAACAGGGTCTGCTGGTTATTG |
| ChiR$_{\text{DBD_LH_H(A)}}$ | ACAAAACCAATAACCAGCAGATTATTGTCCAGCTTGGGAATG | AlsR$_{\text{LBD(C)}}$ | TTCCCAAGCTGGACAATAATCTGCTGGTTATTGGTTTTGTTGGTAG |
| ChiR$_{\text{DBD(B)}}$ | TAAAAAAAGTGACCCTCAGGCAGGGTAATCATGGGGCCATTG | BenM$_{\text{LBD_H_LH(C)}}$ | ATGGCCCCATGATTACCCTGCCTGAGGGTCACTTTTTTTATCAGTACGCCATC |
| ChiR$_{\text{DBD_LH(B)}}$ | ATGGTTTTTTCAACCGAGGCAATGTTGCGCACCCGGCTTTC | BenM$_{\text{LBD_H(C)}}$ | AAAGCCGGGTGCGCAACATTGCCTCGGTTGAAAAAACCATTCG |
| ChiR$_{\text{DBD_LH_H(B)}}$ | CCCACAAATCCGATACGAATATTATTGTCCAGCTTGGGAATG | BenM$_{\text{LBD(C)}}$ | TTCCCAAGCTGGACAATAATATTCGTATCGGATTTGTGGG |
| ChiR$_{\text{DBD(G)}}$ | AGAAATTCTTTACCAATCGGCAGGGTAATCATGGGGCCATTG | GltC$_{\text{LBD_H_LH(C)}}$ | ATGGCCCCATGATTACCCTGCCGATTGGTAAAGAATTTCTGATCC |
| ChiR$_{\text{DBD_LH(G)}}$ | ACGGTGCCACGATGCGGATCAATGTTGCGCACCCGGCTTTC | GltC$_{\text{LBD_H(C)}}$ | AAAGCCGGGTGCGCAACATTGATCCGCATCGTGGCACCGTTAAAATTG |
| ChiR$_{\text{DBD_LH_H(G)}}$ | GTCGGAAAACCAATTTTAACATTATTGTCCAGCTTGGGAATG | GltC$_{\text{LBD(C)}}$ | TTCCCAAGCTGGACAATAATGTTAAAATTGGTTTTCCGACCAGC |
| ChiR$_{\text{DBD(L)}}$ | GCGATCGCCTCGGCACGTGGCAGGGTAATCATGGGGCCATTG | LinR$_{\text{LBD_H_LH(C)}}$ | ATGGCCCCATGATTACCCTGCCACGTGCCGAGGCGATC |
| ChiR$_{\text{DBD_LH(L)}}$ | CGCTTAAGACGGTCAGGTGAAATGTTGCGCACCCGGCTTTC | LinR$_{\text{LBD_H(C)}}$ | AAAGCCGGGTGCGCAACATTTCACCTGACCGTCTTAAGC |
| ChiR$_{\text{DBD_LH_H(L)}}$ | CCGGCAATGATGAATTCGCGATTATTGTCCAGCTTGGGAATG | LinR$_{\text{LBD(C)}}$ | TTCCCAAGCTGGACAATAATCGCGAATTCATCATTGCCGGATCGGAC |
| ChiR$_{\text{DBD(M)}}$ | AGCAGAATTTCACCCTGTGCCGTCAGGGTAATCATGGGGCCATTG | MetR$_{\text{LBD_H_LH(C)}}$ | GCCCCATGATTACCCTGACGGCACAGGGTGAAATTCTGCTGCAGCTGTC |
| ChiR$_{\text{DBD_LH(M)}}$ | ATCTGGTGGGTCTTTTTGCAAATGTTGCGCACCCGGCTTTC | MetR$_{\text{LBD_H(C)}}$ | AAAGCCGGGTGCGCAACATTTGCAAAAAGACCCACCAG |
| ChiR$_{\text{DBD_LH_H(M)}}$ | ATGGCCAGTTTGATAATCATATTATTGTCCAGCTTGGGAATG | MetR$_{\text{LBD(C)}}$ | TTCCCAAGCTGGACAATAATATGATTATCAAACTGGCCATTG |
| GltC$_{\text{DBD(A)}}$ | AGAAAAAATTTCACCGGCTGCGGTCAGTTTAATGTTACGACCTTCAC | AlsR$_{\text{LBD_H_LH(G)}}$ | GTCGTAACATTAAACTGACCGCAGCCGGTGAAATTTTTCTG |
| GltC$_{\text{DBD_LH(A)}}$ | AGACCCTGTTCACCACGTGCCAGATATTCATCGATTTGTTCTTTGGCGTAATCAATGGC | AlsR$_{\text{LBD_H(G)}}$ | AACAAATCGATGAATATCTGGCACGTGGTGAACAGGGTCTGCTGGTTATTG |
| GltC$_{\text{DBD_LH_H(A)}}$ | ACAAAACCAATAACCAGCAGGGTGCCACGATGCGGATCCAGATATTC | AlsR$_{\text{LBD(G)}}$ | TGGATCCGCATCGTGGCACCCTGCTGGTTATTGGTTTTGTTGGTAG |
| GltC$_{\text{DBD(B)}}$ | TAAAAAAAGTGACCCTCAGGGGTCAGTTTAATGTTACGACCTTCAC | BenM$_{\text{LBD_H_LH(G)}}$ | GTCGTAACATTAAACTGACCCCTGAGGGTCACTTTTTTTATCAGTACGCCATC |
| GltC$_{\text{DBD_LH(B)}}$ | ATGGTTTTTTCAACCGAGGCCAGATATTCATCGATTTGTTCTTTGGCGTAATCAATGGC | BenM$_{\text{LBD_H(G)}}$ | AACAAATCGATGAATATCTGGCCTCGGTTGAAAAAACCATTCG |
| GltC$_{\text{DBD_LH_H(B)}}$ | CCCACAAATCCGATACGAATGGTGCCACGATGCGGATCCAGATATTC | BenM$_{\text{LBD(G)}}$ | TGGATCCGCATCGTGGCACCATTCGTATCGGATTTGTGGG |
| GltC$_{\text{DBD(C)}}$ | AGAAATTCCCCTTCATCCGTGGTCAGTTTAATGTTACGACCTTCAC | ChiR$_{\text{LBD_H_LH(G)}}$ | GTCGTAACATTAAACTGACCACGGATGAAGGGGAATTTCTC |
| GltC$_{\text{DBD_LH(C)}}$ | ATATTATTGTCCAGCTTGGGCAGATATTCATCGATTTGTTCTTTGGCGTAATCAATGGC | ChiR$_{\text{LBD_H(G)}}$ | AACAAATCGATGAATATCTGCCCAAGCTGGACAATAATATCCGCTTGGCGGTAG |
| GltC$_{\text{DBD_LH_H(C)}}$ | GTGTCTACCGCCAAGCGGATGGTGCCACGATGCGGATCCAGATATTC | ChiR$_{\text{LBD(G)}}$ | TGGATCCGCATCGTGGCACCATCCGCTTGGCGGTAGACAC |
| GltC$_{\text{DBD(L)}}$ | GCGATCGCCTCGGCACGTGGGGTCAGTTTAATGTTACGACCTTCAC | LinR$_{\text{LBD_H_LH(G)}}$ | GTCGTAACATTAAACTGACCCCACGTGCCGAGGCGATC |
| GltC$_{\text{DBD_LH(L)}}$ | CGCTTAAGACGGTCAGGTGACAGATATTCATCGATTTGTTCTTTGGCGTAATCAATGGC | LinR$_{\text{LBD_H(G)}}$ | AACAAATCGATGAATATCTGTCACCTGACCGTCTTAAGC |
| GltC$_{\text{DBD_LH_H(L)}}$ | CCGGCAATGATGAATTCGCGGGTGCCACGATGCGGATCCAGATATTC | LinR$_{\text{LBD(G)}}$ | TGGATCCGCATCGTGGCACCCGCGAATTCATCATTGCCGGATCGGAC |
| GltC$_{\text{DBD(M)}}$ | AGCAGAATTTCACCCTGTGCGGTCAGTTTAATGTTACGACCTTCAC | MetR$_{\text{LBD_H_LH(G)}}$ | GTCGTAACATTAAACTGACCGCACAGGGTGAAATTCTGCTGCAGCTGTC |
| GltC$_{\text{DBD_LH(M)}}$ | ATCTGGTGGGTCTTTTTGCAATATTCATCGATTTGTTCTTTGGC | MetR$_{\text{LBD_H(G)}}$ | AAGAACAAATCGATGAATATTGCAAAAAGACCCACCAG |
| GltC$_{\text{DBD_LH_H(M)}}$ | ATGGCCAGTTTGATAATCATGCCACGATGCGGATCCAGATATTC | MetR$_{\text{LBD(G)}}$ | ATCTGGATCCGCATCGTGGCATGATTATCAAACTGGCCATTG |
| LinR$_{\text{DBD(A)}}$ | AGAAAAAATTTCACCGGCTGCTGTCGGCTCCATGCCGTCGC | AlsR$_{\text{LBD_H_LH(L)}}$ | GCGACGGCATGGAGCCGACAGCAGCCGGTGAAATTTTTCTG |
| LinR$_{\text{DBD_LH(A)}}$ | AGACCCTGTTCACCACGTGCGAATGTCTGCCCGCCTTCAG | AlsR$_{\text{LBD_H(L)}}$ | CTGAAGGCGGGCAGACATTCGCACGTGGTGAACAGGGTCTGCTGGTTATTG |
| LinR$_{\text{DBD_LH_H(A)}}$ | ACAAAACCAATAACCAGCAGCTTAAGACGGTCAGGTGAGAATGTCTGC | AlsR$_{\text{LBD(L)}}$ | TCTCACCTGACCGTCTTAAGCTGCTGGTTATTGGTTTTGTTGGTAG |
| LinR$_{\text{DBD(B)}}$ | TAAAAAAAGTGACCCTCAGGTGTCGGCTCCATGCCGTCGC | BenM$_{\text{LBD_H_LH(L)}}$ | GCGACGGCATGGAGCCGACACCTGAGGGTCACTTTTTTTATCAGTACGCCATC |
| LinR$_{\text{DBD_LH(B)}}$ | ATGGTTTTTTCAACCGAGGCGAATGTCTGCCCGCCTTCAG | BenM$_{\text{LBD_H(L)}}$ | CTGAAGGCGGGCAGACATTCGCCTCGGTTGAAAAAACCATTCG |
| LinR$_{\text{DBD_LH_H(B)}}$ | CCCACAAATCCGATACGAATCTTAAGACGGTCAGGTGAGAATGTCTGC | BenM$_{\text{LBD(L)}}$ | TCTCACCTGACCGTCTTAAGATTCGTATCGGATTTGTGGG |
| LinR$_{\text{DBD(C)}}$ | AGAAATTCCCCTTCATCCGTTGTCGGCTCCATGCCGTCGC | ChiR$_{\text{LBD_H_LH(L)}}$ | GCGACGGCATGGAGCCGACAACGGATGAAGGGGAATTTCTC |
| LinR$_{\text{DBD_LH(C)}}$ | ATATTATTGTCCAGCTTGGGGAATGTCTGCCCGCCTTCAG | ChiR$_{\text{LBD_H(L)}}$ | CTGAAGGCGGGCAGACATTCCCCAAGCTGGACAATAATATCCGCTTGGCGGTAG |
| LinR$_{\text{DBD_LH_H(C)}}$ | GTGTCTACCGCCAAGCGGATCTTAAGACGGTCAGGTGAGAATGTCTGC | ChiR$_{\text{LBD(L)}}$ | TCTCACCTGACCGTCTTAAGATCCGCTTGGCGGTAGACAC |
| LinR$_{\text{DBD(G)}}$ | AGAAATTCTTTACCAATCGGTGTCGGCTCCATGCCGTCGC | GltC$_{\text{LBD_H_LH(L)}}$ | GCGACGGCATGGAGCCGACACCGATTGGTAAAGAATTTCTGATCC |

| | | | |
|---|---|---|---|
| LinR$_{DBD_LH(G)}$ | ACGGTGCCACGATGCGGATCGAATGTCTGCCCGCCTTCAG | GltC$_{LBD_H(L)}$ | CTGAAGGCGGGCAGACATTCGATCCGCATCGTGGCACCGTTAAAATTG |
| LinR$_{DBD_LH_H(G)}$ | GTCGGAAAACCAATTTTAACCTTAAGACGGTCAGGTGAGAATGTCTGC | GltC$_{LBD(L)}$ | TCTCACCTGACCGTCTTAAGGTTAAAATTGGTTTTCCGACCAGC |
| LinR$_{DBD(M)}$ | AGCAGAATTTCACCCTGTGCTGTCGGCTCCATGCCGTCGC | MetR$_{LBD_H_LH(L)}$ | GCGACGGCATGGAGCCGACAGCACAGGGTGAAATTCTGCTGCAGCTGTC |
| LinR$_{DBD_LH(M)}$ | ATCTGGTGGGTCTTTTTGCAGAATGTCTGCCCGCCTTCAG | MetR$_{LBD_H(L)}$ | CTGAAGGCGGGCAGACATTCTGCAAAAAGACCCACCAG |
| LinR$_{DBD_LH_H(M)}$ | ATGGCCAGTTTGATAATCATCTTAAGACGGTCAGGTGAGAATGTCTGC | MetR$_{LBD(L)}$ | TCTCACCTGACCGTCTTAAGATGATTATCAAACTGGCCATTG |
| MetR$_{DBD(A)}$ | AGAAAAAATTTCACCGGCTGCGGTAAATTTGATCGGGTTGC | AlsR$_{LBD_H_LH(M)}$ | GCAACCCGATCAAATTTACCGCAGCCGGTGAAATTTTTCTG |
| MetR$_{DBD_LH(A)}$ | AGACCCTGTTCACCACGTGCATTCTGGATGGCTTTATGAATTTTCGG | AlsR$_{LBD_H(M)}$ | TTCATAAAGCCATCCAGAATGCACGTGGTGAACAGGGTCTGCTGGTTATTG |
| MetR$_{DBD_LH_H(A)}$ | ACAAAACCAATAACCAGCAGCTGGTGGGTCTTTTTGCAATTC | AlsR$_{LBD(M)}$ | ATTGCAAAAAGACCCACCAGCTGCTGGTTATTGGTTTTGTTGGTAG |
| MetR$_{DBD(B)}$ | TAAAAAAAGTGACCCTCAGGGGTAAATTTGATCGGGTTGC | BenM$_{LBD_H_LH(M)}$ | GCAACCCGATCAAATTTACCCCTGAGGGTCACTTTTTTTATCAGTACGCCATC |
| MetR$_{DBD_LH(B)}$ | ATGGTTTTTTCAACCGAGGCATTCTGGATGGCTTTATGAATTTTCGG | BenM$_{LBD_H(M)}$ | TTCATAAAGCCATCCAGAATGCCTCGGTTGAAAAAACCATTCG |
| MetR$_{DBD_LH_H(B)}$ | CCCACAAATCCGATACGAATCTGGTGGGTCTTTTTGCAATTC | BenM$_{LBD(M)}$ | ATTGCAAAAAGACCCACCAGATTCGTATCGGATTTGTGGG |
| MetR$_{DBD(C)}$ | AGGAGAAATTCCCCTTCATCGGTAAATTTGATCGGGTTGC | ChiR$_{LBD_H_LH(M)}$ | GCAACCCGATCAAATTTACCGATGAAGGGGAATTTCTCCTGC |
| MetR$_{DBD_LH(C)}$ | ATATTATTGTCCAGCTTGGGATTCTGGATGGCTTTATGAATTTTCGG | ChiR$_{LBD_H(M)}$ | TTCATAAAGCCATCCAGAATCCCAAGCTGGACAATAATATCCGCTTGGCGGTAG |
| MetR$_{DBD_LH_H(C)}$ | GTGTCTACCGCCAAGCGGATCTGGTGGGTCTTTTTGCAATTC | ChiR$_{LBD(M)}$ | ATTGCAAAAAGACCCACCAGATCCGCTTGGCGGTAGACAC |
| MetR$_{DBD(G)}$ | AGAAATTCTTTACCAATCGGGGTAAATTTGATCGGGTTGC | GltC$_{LBD_H_LH(M)}$ | GCAACCCGATCAAATTTACCCCGATTGGTAAAGAATTTCTGATCC |
| MetR$_{DBD_LH(G)}$ | GTGCCACGATGCGGATCCAGATTCTGGATGGCTTTATGAATTTTCGG | GltC$_{LBD_H(M)}$ | TTCATAAAGCCATCCAGAATCTGGATCCGCATCGTGGCACCGTTAAAATTG |
| MetR$_{DBD_LH_H(G)}$ | GGAAAACCAATTTTAACGGTCTGGTGGGTCTTTTTGCAATTC | GltC$_{LBD(M)}$ | ATTGCAAAAAGACCCACCAGACCGTTAAAATTGGTTTTCCGACCAGC |
| MetR$_{DBD(L)}$ | GCGATCGCCTCGGCACGTGGGGTAAATTTGATCGGGTTGC | LinR$_{LBD_H_LH(M)}$ | GCAACCCGATCAAATTTACCCCACGTGCCGAGGCGATC |
| MetR$_{DBD_LH(L)}$ | CGCTTAAGACGGTCAGGTGAATTCTGGATGGCTTTATGAATTTTCGG | LinR$_{LBD_H(M)}$ | TTCATAAAGCCATCCAGAATTCACCTGACCGTCTTAAGC |
| MetR$_{DBD_LH_H(L)}$ | CCGGCAATGATGAATTCGCGCTGGTGGGTCTTTTTGCAATTC | LinR$_{LBD(M)}$ | ATTGCAAAAAGACCCACCAGCGCGAATTCATCATTGCCGGATCGGAC |
| TsaR$_{DBD(A)}$ | AGAAAAAATTTCACCGGCTGCGGTCAGGCTAACACCACGTTTG | AlsR$_{LBD_H_LH(T)}$ | AACGTGGTGTTAGCCTGACCGCAGCCGGTGAAATTTTTCTG |
| TsaR$_{DBD_LH(A)}$ | AGACCCTGTTCACCACGTGCCAGCTGACCAATTTCTTCTTGTGCACGACG | AlsR$_{LBD_H(T)}$ | AAGAAGAAATTGGTCAGCTGGCACGTGGTGAACAGGGTCTGCTGGTTATTG |
| TsaR$_{DBD_LH_H(A)}$ | ACAAAACCAATAACCAGCAGACCTTCCCAACGACCACGCAG | AlsR$_{LBD(T)}$ | TGCGTGGTCGTTGGGAAGGTCTGCTGGTTATTGGTTTTGTTGGTAG |
| TsaR$_{DBD(B)}$ | TAAAAAAAGTGACCCTCAGGGGTCAGGCTAACACCACGTTTG | BenM$_{LBD_H_LH(T)}$ | AACGTGGTGTTAGCCTGACCCCTGAGGGTCACTTTTTTTATCAGTACGCCATC |
| TsaR$_{DBD_LH(B)}$ | ATGGTTTTTTCAACCGAGGCCAGCTGACCAATTTCTTCTTGTGCACGACG | BenM$_{LBD_H(T)}$ | AAGAAGAAATTGGTCAGCTGGCCTCGGTTGAAAAAACCATTCG |
| TsaR$_{DBD_LH_H(B)}$ | CCCACAAATCCGATACGAATACCTTCCCAACGACCACGCAG | BenM$_{LBD(T)}$ | TGCGTGGTCGTTGGGAAGGTATTCGTATCGGATTTGTGGG |
| TsaR$_{DBD(C)}$ | AGAAATTCCCCTTCATCCGTGGTCAGGCTAACACCACGTTTG | ChiR$_{LBD_H_LH(T)}$ | AACGTGGTGTTAGCCTGACCACGGATGAAGGGGAATTTCTC |
| TsaR$_{DBD_LH(C)}$ | ATATTATTGTCCAGCTTGGGCAGCTGACCAATTTCTTCTTGTGCACGACG | ChiR$_{LBD_H(T)}$ | AAGAAGAAATTGGTCAGCTGCCCAAGCTGGACAATAATATCCGCTTGGCGGTAG |
| TsaR$_{DBD_LH_H(C)}$ | GTGTCTACCGCCAAGCGGATACCTTCCCAACGACCACGCAG | ChiR$_{LBD(T)}$ | TGCGTGGTCGTTGGGAAGGTATCCGCTTGGCGGTAGACAC |
| TsaR$_{DBD(G)}$ | AGAAATTCTTTACCAATCGGGGTCAGGCTAACACCACGTTTG | GltC$_{LBD_H_LH(T)}$ | AACGTGGTGTTAGCCTGACCCCGATTGGTAAAGAATTTCTGATCC |
| TsaR$_{DBD_LH(G)}$ | ACGGTGCCACGATGCGGATCCAGCTGACCAATTTCTTCTTGTGCACGACG | GltC$_{LBD_H(T)}$ | AAGAAGAAATTGGTCAGCTGGATCCGCATCGTGGCACCGTTAAAATTG |
| TsaR$_{DBD_LH_H(G)}$ | GTCGGAAAACCAATTTTAACACCTTCCCAACGACCACGCAG | GltC$_{LBD(T)}$ | TGCGTGGTCGTTGGGAAGGTGTTAAAATTGGTTTTCCGACCAGC |
| TsaR$_{DBD(L)}$ | GCGATCGCCTCGGCACGTGGGGTCAGGCTAACACCACGTTTG | LinR$_{LBD_H_LH(T)}$ | AACGTGGTGTTAGCCTGACCCCACGTGCCGAGGCGATC |
| TsaR$_{DBD_LH(L)}$ | CGCTTAAGACGGTCAGGTGACAGCTGACCAATTTCTTCTTGTGCACGACG | LinR$_{LBD_H(T)}$ | AAGAAGAAATTGGTCAGCTGTCACCTGACCGTCTTAAGC |
| TsaR$_{DBD_LH_H(L)}$ | CCGGCAATGATGAATTCGCGACCTTCCCAACGACCACGCAG | LinR$_{LBD(T)}$ | TGCGTGGTCGTTGGGAAGGTCGCGAATTCATCATTGCCGGATCGGAC |
| TsaR$_{DBD(M)}$ | AGCAGAATTTCACCCTGTGCGGTCAGGCTAACACCACGTTTG | MetR$_{LBD_H_LH(T)}$ | AACGTGGTGTTAGCCTGACCGCACAGGGTGAAATTCTGCTGCAGCTGTC |
| TsaR$_{DBD_LH(M)}$ | ATCTGGTGGGTCTTTTTGCACAGCTGACCAATTTCTTCTTGTGCACGACG | MetR$_{LBD_H(T)}$ | AAGAAGAAATTGGTCAGCTGTGCAAAAAGACCCACCAG |
| TsaR$_{DBD_LH_H(M)}$ | ATGGCCAGTTTGATAATCATACCTTCCCAACGACCACGCAG | MetR$_{LBD(T)}$ | TGCGTGGTCGTTGGGAAGGTATGATTATCAAACTGGCCATTG |

| | | | |
|---|---|---|---|
| **Backbone RV** | AACAGCAGGCGGCGTTCTTC | **Backbone FW** | GTGCGCCTTCTCGAAGAAC |

**S2b — Orthogonality case-study primers**

| **PCR part** | **FW_primer** | **RV_primer** |
|---|---|---|
| Chimera_part | CAGAGCAGCCGATTGTCTGTTGTGCCCAGTCATAG | GAGGAGCGCCTTCAGTTGAC |
| P_part | GTCAACTGAAGGCGCTCCTC | GCTATTCGGCTATGACTGGGCACAACAGACAATCG |

**Supplementary Table S3 | Plasmids and strains used in this work.** Every plasmid/strain (each E. coli TOP10 strain carries one plasmid), grouped as control, biosensor, orthogonality-test and domain-swapped-chimera constructs.

| Strain | Transcription factor/chimera | Promoter region | Source |
|---|---|---|---|
| **Control plasmids** | | | |
| SynJunk | TF_junk | PTFBS_junk | Demeester 2023 |
| ProB_mKate2 | TF_junk | ProB_mKate2 | Demeester 2023 |
| PAllS | TF_junk | $P_{allD}$ | This work |
| PAlsR | TF_junk | $P_{alsS/alsR}$ | This work |
| PAmpR | TF_junk | $P_{ampC/ampR}$ | This work |
| PBenM | TF_junk | $P_{benA/benM}$ | This work |
| PChiR | TF_junk | $P_{CBP21/chiR}$ | This work |
| PCitR | TF_junk | $P_{citA/citR}$ | This work |
| PCysB | TF_junk | $P_{cysI/cysB}$ | This work |
| PFdeR | TF_junk | $P_{fdeA/fdeR}$ | This work |
| PGltC | TF_junk | $P_{gltA/gltC}$ | This work |
| PLinR | TF_junk | $P_{linE/linR}$ | Demeester 2023 |
| PLysG | TF_junk | $P_{lysE/lysG}$ | This work |
| PMetR | TF_junk | $P_{metE/metR}$ | Demeester 2023 |
| PNagR | TF_junk | $P_{nagA/nagR}$ | This work |
| PNahR | TF_junk | $P_{nahRmar}$ | This work |
| POccR | TF_junk | $P_{occQ/occR}$ | This work |
| PPqsR | TF_junk | $P_{pqsA\_trunc}$ | This work |
| PTsaR | TF_junk | $P_{tsaM/tsaR}$ | This work |
| PTtuA | TF_junk | $P_{ttuB/ttuA}$ | This work |
| **Biosensor plasmids** | | | |
| SynAllS | AllS | $P_{allD}$ | This work |
| SynAlsR | AlsR | $P_{alsS/alsR}$ | This work |
| NatAlsR | AlsR | $P_{alsS/alsR}$ | This work |
| SynAmpR | AmpR | $P_{ampC/ampR}$ | This work |
| NatAmpR | AmpR | $P_{ampC/ampR}$ | This work |
| SynBenM | BenM | $P_{benA/benM}$ | This work |
| NatBenM | BenM | $P_{benA/benM}$ | This work |
| SynChiR | ChiR | $P_{CBP21/chiR}$ | This work |
| NatChiR | ChiR | $P_{CBP21/chiR}$ | This work |
| SynCitR | CitR | $P_{citA/citR}$ | This work |
| NatCitR | CitR | $P_{citA/citR}$ | This work |
| SynCysB | CysB | $P_{cysI/cysB}$ | This work |
| NatCysB | CysB | $P_{cysI/cysB}$ | This work |
| SynFdeR | FdeR | $P_{fdeA/fdeR}$ | This work |
| NatFdeR | FdeR | $P_{fdeA/fdeR}$ | This work |
| SynGltC | GltC | $P_{gltA/gltC}$ | This work |
| NatGltC | GltC | $P_{gltA/gltC}$ | This work |
| SynLinR | LinR | $P_{linE/linR}$ | Demeester 2023 |
| NatLinR | LinR | $P_{linE/linR}$ | Demeester 2023 |
| SynLysG | LysG | $P_{lysE/lysG}$ | This work |
| NatLysG | LysG | $P_{lysE/lysG}$ | This work |
| SynMetR | MetR | $P_{metE/metR}$ | Demeester 2023 |
| NatMetR | MetR | $P_{metE/metR}$ | Demeester 2023 |

| | | | |
|---|---|---|---|
| SynNagR | NagR | $P_{nagA/nagR}$ | This work |
| NatNagR | NagR | $P_{nagA/nagR}$ | This work |
| SynNahR | NahR | $P_{nahRmar}$ | This work |
| NatNahR | NahR | $P_{nahRmar}$ | This work |
| SynOccR | OccR | $P_{occQ/occR}$ | This work |
| NatOccR | OccR | $P_{occQ/occR}$ | This work |
| SynPqsR | PqsR | $P_{pqsA\_trunc}$ | This work |
| SynTsaR | TsaR | $P_{tsaM/tsaR}$ | This work |
| NatTsaR | TsaR | $P_{tsaM/tsaR}$ | This work |
| SynTtuA | TtuA | $P_{ttuB/ttuA}$ | This work |
| NatTtuA | TtuA | $P_{ttuB/ttuA}$ | This work |
| **Orthogonality test plasmids** | | | |
| PAlsRSynBenM | BenM | $P_{alsS/alsR}$ | This work |
| PAlsRSynChiR | ChiR | $P_{alsS/alsR}$ | This work |
| PAlsRSynCitR | CitR | $P_{alsS/alsR}$ | This work |
| PAlsRSynCysB | CysB | $P_{alsS/alsR}$ | This work |
| PAlsRSynFdeR | FdeR | $P_{alsS/alsR}$ | This work |
| PAlsRSynGltC | GltC | $P_{alsS/alsR}$ | This work |
| PAlsRSynLinR | LinR | $P_{alsS/alsR}$ | This work |
| PAlsRSynMetR | MetR | $P_{alsS/alsR}$ | This work |
| PAlsRSynNahR | NahR | $P_{alsS/alsR}$ | This work |
| PAlsRSynTsaR | TsaR | $P_{alsS/alsR}$ | This work |
| PBenMSynAlsR | AlsR | $P_{benA/benM}$ | This work |
| PBenMSynChiR | ChiR | $P_{benA/benM}$ | This work |
| PBenMSynCitR | CitR | $P_{benA/benM}$ | This work |
| PBenMSynCysB | CysB | $P_{benA/benM}$ | This work |
| PBenMSynFdeR | FdeR | $P_{benA/benM}$ | This work |
| PBenMSynGltC | GltC | $P_{benA/benM}$ | This work |
| PBenMSynLinR | LinR | $P_{benA/benM}$ | This work |
| PBenMSynMetR | MetR | $P_{benA/benM}$ | This work |
| PBenMSynNahR | NahR | $P_{benA/benM}$ | This work |
| PBenMSynTsaR | TsaR | $P_{benA/benM}$ | This work |
| PChiRSynAlsR | AlsR | $P_{CBP21/chiR}$ | This work |
| PChiRSynBenM | BenM | $P_{CBP21/chiR}$ | This work |
| PChiRSynCitR | CitR | $P_{CBP21/chiR}$ | This work |
| PChiRSynCysB | CysB | $P_{CBP21/chiR}$ | This work |
| PChiRSynFdeR | FdeR | $P_{CBP21/chiR}$ | This work |
| PChiRSynGltC | GltC | $P_{CBP21/chiR}$ | This work |
| PChiRSynLinR | LinR | $P_{CBP21/chiR}$ | This work |
| PChiRSynMetR | MetR | $P_{CBP21/chiR}$ | This work |
| PChiRSynNahR | NahR | $P_{CBP21/chiR}$ | This work |
| PChiRSynTsaR | TsaR | $P_{CBP21/chiR}$ | This work |
| PCitRSynAlsR | AlsR | $P_{citA/citR}$ | This work |
| PCitRSynBenM | BenM | $P_{citA/citR}$ | This work |
| PCitRSynChiR | ChiR | $P_{citA/citR}$ | This work |
| PCitRSynCysB | CysB | $P_{citA/citR}$ | This work |
| PCitRSynFdeR | FdeR | $P_{citA/citR}$ | This work |
| PCitRSynGltC | GltC | $P_{citA/citR}$ | This work |
| PCitRSynLinR | LinR | $P_{citA/citR}$ | This work |

| | | | |
|---|---|---|---|
| PCitRSynMetR | MetR | $P_{citA/citR}$ | This work |
| PCitRSynNahR | NahR | $P_{citA/citR}$ | This work |
| PCitRSynTsaR | TsaR | $P_{citA/citR}$ | This work |
| PCysBSynAlsR | AlsR | $P_{cysI/cysB}$ | This work |
| PCysBSynBenM | BenM | $P_{cysI/cysB}$ | This work |
| PCysBSynChiR | ChiR | $P_{cysI/cysB}$ | This work |
| PCysBSynCitR | CitR | $P_{cysI/cysB}$ | This work |
| PCysBSynFdeR | FdeR | $P_{cysI/cysB}$ | This work |
| PCysBSynGltC | GltC | $P_{cysI/cysB}$ | This work |
| PCysBSynLinR | LinR | $P_{cysI/cysB}$ | This work |
| PCysBSynMetR | MetR | $P_{cysI/cysB}$ | This work |
| PCysBSynNahR | NahR | $P_{cysI/cysB}$ | This work |
| PCysBSynTsaR | TsaR | $P_{cysI/cysB}$ | This work |
| PFdeRSynAlsR | AlsR | $P_{fdeA/fdeR}$ | This work |
| PFdeRSynBenM | BenM | $P_{fdeA/fdeR}$ | This work |
| PFdeRSynChiR | ChiR | $P_{fdeA/fdeR}$ | This work |
| PFdeRSynCitR | CitR | $P_{fdeA/fdeR}$ | This work |
| PFdeRSynCysB | CysB | $P_{fdeA/fdeR}$ | This work |
| PFdeRSynGltC | GltC | $P_{fdeA/fdeR}$ | This work |
| PFdeRSynMetR | MetR | $P_{fdeA/fdeR}$ | This work |
| PFdeRSynNahR | NahR | $P_{fdeA/fdeR}$ | This work |
| PFdeRSynTsaR | TsaR | $P_{fdeA/fdeR}$ | This work |
| PGltCSynAlsR | AlsR | $P_{gltA/gltC}$ | This work |
| PGltCSynBenM | BenM | $P_{gltA/gltC}$ | This work |
| PGltCSynChiR | ChiR | $P_{gltA/gltC}$ | This work |
| PGltCSynCitR | CitR | $P_{gltA/gltC}$ | This work |
| PGltCSynCysB | CysB | $P_{gltA/gltC}$ | This work |
| PGltCSynFdeR | FdeR | $P_{gltA/gltC}$ | This work |
| PGltCSynLinR | LinR | $P_{gltA/gltC}$ | This work |
| PGltCSynMetR | MetR | $P_{gltA/gltC}$ | This work |
| PGltCSynNahR | NahR | $P_{gltA/gltC}$ | This work |
| PGltCSynTsaR | TsaR | $P_{gltA/gltC}$ | This work |
| PLinRSynAlsR | AlsR | $P_{linE/linR}$ | This work |
| PLinRSynBenM | BenM | $P_{linE/linR}$ | This work |
| PLinRSynChiR | ChiR | $P_{linE/linR}$ | This work |
| PLinRSynCitR | CitR | $P_{linE/linR}$ | This work |
| PLinRSynCysB | CysB | $P_{linE/linR}$ | This work |
| PLinRSynFdeR | FdeR | $P_{linE/linR}$ | This work |
| PLinRSynGltC | GltC | $P_{linE/linR}$ | This work |
| PLinRSynMetR | MetR | $P_{linE/linR}$ | This work |
| PLinRSynNahR | NahR | $P_{linE/linR}$ | This work |
| PLinRSynTsaR | TsaR | $P_{linE/linR}$ | This work |
| PMetRSynAlsR | AlsR | $P_{metE/metR}$ | This work |
| PMetRSynBenM | BenM | $P_{metE/metR}$ | This work |
| PMetRSynChiR | ChiR | $P_{metE/metR}$ | This work |
| PMetRSynCysB | CysB | $P_{metE/metR}$ | This work |
| PMetRSynFdeR | FdeR | $P_{metE/metR}$ | This work |
| PMetRSynGltC | GltC | $P_{metE/metR}$ | This work |
| PMetRSynLinR | LinR | $P_{metE/metR}$ | This work |

| | | | |
|---|---|---|---|
| PMetRSynNahR | NahR | $P_{metE/metR}$ | This work |
| PMetRSynTsaR | TsaR | $P_{metE/metR}$ | This work |
| PNahRSynAlsR | AlsR | $P_{nahRmar}$ | This work |
| PNahRSynBenM | BenM | $P_{nahRmar}$ | This work |
| PNahRSynChiR | ChiR | $P_{nahRmar}$ | This work |
| PNahRSynCitR | CitR | $P_{nahRmar}$ | This work |
| PNahRSynCysB | CysB | $P_{nahRmar}$ | This work |
| PNahRSynFdeR | FdeR | $P_{nahRmar}$ | This work |
| PNahRSynGltC | GltC | $P_{nahRmar}$ | This work |
| PNahRSynLinR | LinR | $P_{nahRmar}$ | This work |
| PNahRSynMetR | MetR | $P_{nahRmar}$ | This work |
| PNahRSynTsaR | TsaR | $P_{nahRmar}$ | This work |
| PTsaRSynAlsR | AlsR | $P_{tsaM/tsaR}$ | This work |
| PTsaRSynBenM | BenM | $P_{tsaM/tsaR}$ | This work |
| PTsaRSynChiR | ChiR | $P_{tsaM/tsaR}$ | This work |
| PTsaRSynCysB | CysB | $P_{tsaM/tsaR}$ | This work |
| PTsaRSynFdeR | FdeR | $P_{tsaM/tsaR}$ | This work |
| PTsaRSynGltC | GltC | $P_{tsaM/tsaR}$ | This work |
| PTsaRSynLinR | LinR | $P_{tsaM/tsaR}$ | This work |
| PTsaRSynMetR | MetR | $P_{tsaM/tsaR}$ | This work |
| PTsaRSynNahR | NahR | $P_{tsaM/tsaR}$ | This work |
| **Domain swapped chimera** | | | |
| SynAlsR$_{DBD}$BenM$_{LBD_H_LH}$ | AlsR$_{DBD}$BenM$_{LBD_H_LH}$ | $P_{alsS/alsR}$ | This work |
| SynAlsR$_{DBD_LH}$BenM$_{LBD_H}$ | AlsR$_{DBD_LH}$BenM$_{LBD_H}$ | $P_{alsS/alsR}$ | This work |
| SynAlsR$_{DBD_LH_H}$BenM$_{LBD}$ | AlsR$_{DBD_LH_H}$BenM$_{LBD}$ | $P_{alsS/alsR}$ | This work |
| SynAlsR$_{DBD}$ChiR$_{LBD_H_LH}$ | AlsR$_{DBD}$ChiR$_{LBD_H_LH}$ | $P_{alsS/alsR}$ | This work |
| SynAlsR$_{DBD_LH}$ChiR$_{LBD_H}$ | AlsR$_{DBD_LH}$ChiR$_{LBD_H}$ | $P_{alsS/alsR}$ | This work |
| SynAlsR$_{DBD_LH_H}$ChiR$_{LBD}$ | AlsR$_{DBD_LH_H}$ChiR$_{LBD}$ | $P_{alsS/alsR}$ | This work |
| SynAlsR$_{DBD}$GltC$_{LBD_H_LH}$ | AlsR$_{DBD}$GltC$_{LBD_H_LH}$ | $P_{alsS/alsR}$ | This work |
| SynAlsR$_{DBD_LH}$GltC$_{LBD_H}$ | AlsR$_{DBD_LH}$GltC$_{LBD_H}$ | $P_{alsS/alsR}$ | This work |
| SynAlsR$_{DBD_LH_H}$GltC$_{LBD}$ | AlsR$_{DBD_LH_H}$GltC$_{LBD}$ | $P_{alsS/alsR}$ | This work |
| SynAlsR$_{DBD}$LinR$_{LBD_H_LH}$ | AlsR$_{DBD}$LinR$_{LBD_H_LH}$ | $P_{alsS/alsR}$ | This work |
| SynAlsR$_{DBD_LH}$LinR$_{LBD_H}$ | AlsR$_{DBD_LH}$LinR$_{LBD_H}$ | $P_{alsS/alsR}$ | This work |
| SynAlsR$_{DBD_LH_H}$LinR$_{LBD}$ | AlsR$_{DBD_LH_H}$LinR$_{LBD}$ | $P_{alsS/alsR}$ | This work |
| SynAlsR$_{DBD}$MetR$_{LBD_H_LH}$ | AlsR$_{DBD}$MetR$_{LBD_H_LH}$ | $P_{alsS/alsR}$ | This work |
| SynAlsR$_{DBD_LH}$MetR$_{LBD_H}$ | AlsR$_{DBD_LH}$MetR$_{LBD_H}$ | $P_{alsS/alsR}$ | This work |
| SynAlsR$_{DBD_LH_H}$MetR$_{LBD}$ | AlsR$_{DBD_LH_H}$MetR$_{LBD}$ | $P_{alsS/alsR}$ | This work |
| SynBenM$_{DBD}$AlsR$_{LBD_H_LH}$ | BenM$_{DBD}$AlsR$_{LBD_H_LH}$ | $P_{benA/benM}$ | This work |
| SynBenM$_{DBD_LH}$AlsR$_{LBD_H}$ | BenM$_{DBD_LH}$AlsR$_{LBD_H}$ | $P_{benA/benM}$ | This work |
| SynBenM$_{DBD_LH_H}$AlsR$_{LBD}$ | BenM$_{DBD_LH_H}$AlsR$_{LBD}$ | $P_{benA/benM}$ | This work |
| SynBenM$_{DBD}$ChiR$_{LBD_H_LH}$ | BenM$_{DBD}$ChiR$_{LBD_H_LH}$ | $P_{benA/benM}$ | This work |
| SynBenM$_{DBD_LH}$ChiR$_{LBD_H}$ | BenM$_{DBD_LH}$ChiR$_{LBD_H}$ | $P_{benA/benM}$ | This work |
| SynBenM$_{DBD_LH_H}$ChiR$_{LBD}$ | BenM$_{DBD_LH_H}$ChiR$_{LBD}$ | $P_{benA/benM}$ | This work |
| SynBenM$_{DBD}$GltC$_{LBD_H_LH}$ | BenM$_{DBD}$GltC$_{LBD_H_LH}$ | $P_{benA/benM}$ | This work |
| SynBenM$_{DBD_LH}$GltC$_{LBD_H}$ | BenM$_{DBD_LH}$GltC$_{LBD_H}$ | $P_{benA/benM}$ | This work |
| SynBenM$_{DBD_LH_H}$GltC$_{LBD}$ | BenM$_{DBD_LH_H}$GltC$_{LBD}$ | $P_{benA/benM}$ | This work |
| SynBenM$_{DBD}$LinR$_{LBD_H_LH}$ | BenM$_{DBD}$LinR$_{LBD_H_LH}$ | $P_{benA/benM}$ | This work |
| SynBenM$_{DBD_LH}$LinR$_{LBD_H}$ | BenM$_{DBD_LH}$LinR$_{LBD_H}$ | $P_{benA/benM}$ | This work |
| SynBenM$_{DBD_LH_H}$LinR$_{LBD}$ | BenM$_{DBD_LH_H}$LinR$_{LBD}$ | $P_{benA/benM}$ | This work |

| | | | |
|---|---|---|---|
| SynBenM$_{DBD}$MetR$_{LBD_H_LH}$ | BenM$_{DBD}$MetR$_{LBD_H_LH}$ | P$_{benA/benM}$ | This work |
| SynBenM$_{DBD_LH}$MetR$_{LBD_H}$ | BenM$_{DBD_LH}$MetR$_{LBD_H}$ | P$_{benA/benM}$ | This work |
| SynBenM$_{DBD_LH_H}$MetR$_{LBD}$ | BenM$_{DBD_LH_H}$MetR$_{LBD}$ | P$_{benA/benM}$ | This work |
| SynChiR$_{DBD}$AlsR$_{LBD_H_LH}$ | ChiR$_{DBD}$AlsR$_{LBD_H_LH}$ | P$_{CBP21/chiR}$ | This work |
| SynChiR$_{DBD_LH}$AlsR$_{LBD_H}$ | ChiR$_{DBD_LH}$AlsR$_{LBD_H}$ | P$_{CBP21/chiR}$ | This work |
| SynChiR$_{DBD_LH_H}$AlsR$_{LBD}$ | ChiR$_{DBD_LH_H}$AlsR$_{LBD}$ | P$_{CBP21/chiR}$ | This work |
| SynChiR$_{DBD}$BenM$_{LBD_H_LH}$ | ChiR$_{DBD}$BenM$_{LBD_H_LH}$ | P$_{CBP21/chiR}$ | This work |
| SynChiR$_{DBD_LH}$BenM$_{LBD_H}$ | ChiR$_{DBD_LH}$BenM$_{LBD_H}$ | P$_{CBP21/chiR}$ | This work |
| SynChiR$_{DBD_LH_H}$BenM$_{LBD}$ | ChiR$_{DBD_LH_H}$BenM$_{LBD}$ | P$_{CBP21/chiR}$ | This work |
| SynChiR$_{DBD}$GltC$_{LBD_H_LH}$ | ChiR$_{DBD}$GltC$_{LBD_H_LH}$ | P$_{CBP21/chiR}$ | This work |
| SynChiR$_{DBD_LH}$GltC$_{LBD_H}$ | ChiR$_{DBD_LH}$GltC$_{LBD_H}$ | P$_{CBP21/chiR}$ | This work |
| SynChiR$_{DBD_LH_H}$GltC$_{LBD}$ | ChiR$_{DBD_LH_H}$GltC$_{LBD}$ | P$_{CBP21/chiR}$ | This work |
| SynChiR$_{DBD}$LinR$_{LBD_H_LH}$ | ChiR$_{DBD}$LinR$_{LBD_H_LH}$ | P$_{CBP21/chiR}$ | This work |
| SynChiR$_{DBD_LH}$LinR$_{LBD_H}$ | ChiR$_{DBD_LH}$LinR$_{LBD_H}$ | P$_{CBP21/chiR}$ | This work |
| SynChiR$_{DBD_LH_H}$LinR$_{LBD}$ | ChiR$_{DBD_LH_H}$LinR$_{LBD}$ | P$_{CBP21/chiR}$ | This work |
| SynChiR$_{DBD}$MetR$_{LBD_H_LH}$ | ChiR$_{DBD}$MetR$_{LBD_H_LH}$ | P$_{CBP21/chiR}$ | This work |
| SynChiR$_{DBD_LH}$MetR$_{LBD_H}$ | ChiR$_{DBD_LH}$MetR$_{LBD_H}$ | P$_{CBP21/chiR}$ | This work |
| SynChiR$_{DBD_LH_H}$MetR$_{LBD}$ | ChiR$_{DBD_LH_H}$MetR$_{LBD}$ | P$_{CBP21/chiR}$ | This work |
| SynGltC$_{DBD}$AlsR$_{LBD_H_LH}$ | GltC$_{DBD}$AlsR$_{LBD_H_LH}$ | P$_{gltA/gltC}$ | This work |
| SynGltC$_{DBD_LH}$AlsR$_{LBD_H}$ | GltC$_{DBD_LH}$AlsR$_{LBD_H}$ | P$_{gltA/gltC}$ | This work |
| SynGltC$_{DBD_LH_H}$AlsR$_{LBD}$ | GltC$_{DBD_LH_H}$AlsR$_{LBD}$ | P$_{gltA/gltC}$ | This work |
| SynGltC$_{DBD}$BenM$_{LBD_H_LH}$ | GltC$_{DBD}$BenM$_{LBD_H_LH}$ | P$_{gltA/gltC}$ | This work |
| SynGltC$_{DBD_LH}$BenM$_{LBD_H}$ | GltC$_{DBD_LH}$BenM$_{LBD_H}$ | P$_{gltA/gltC}$ | This work |
| SynGltC$_{DBD_LH_H}$BenM$_{LBD}$ | GltC$_{DBD_LH_H}$BenM$_{LBD}$ | P$_{gltA/gltC}$ | This work |
| SynGltC$_{DBD}$ChiR$_{LBD_H_LH}$ | GltC$_{DBD}$ChiR$_{LBD_H_LH}$ | P$_{gltA/gltC}$ | This work |
| SynGltC$_{DBD_LH}$ChiR$_{LBD_H}$ | GltC$_{DBD_LH}$ChiR$_{LBD_H}$ | P$_{gltA/gltC}$ | This work |
| SynGltC$_{DBD_LH_H}$ChiR$_{LBD}$ | GltC$_{DBD_LH_H}$ChiR$_{LBD}$ | P$_{gltA/gltC}$ | This work |
| SynGltC$_{DBD}$LinR$_{LBD_H_LH}$ | GltC$_{DBD}$LinR$_{LBD_H_LH}$ | P$_{gltA/gltC}$ | This work |
| SynGltC$_{DBD_LH}$LinR$_{LBD_H}$ | GltC$_{DBD_LH}$LinR$_{LBD_H}$ | P$_{gltA/gltC}$ | This work |
| SynGltC$_{DBD_LH_H}$LinR$_{LBD}$ | GltC$_{DBD_LH_H}$LinR$_{LBD}$ | P$_{gltA/gltC}$ | This work |
| SynGltC$_{DBD}$MetR$_{LBD_H_LH}$ | GltC$_{DBD}$MetR$_{LBD_H_LH}$ | P$_{gltA/gltC}$ | This work |
| SynGltC$_{DBD_LH}$MetR$_{LBD_H}$ | GltC$_{DBD_LH}$MetR$_{LBD_H}$ | P$_{gltA/gltC}$ | This work |
| SynGltC$_{DBD_LH_H}$MetR$_{LBD}$ | GltC$_{DBD_LH_H}$MetR$_{LBD}$ | P$_{gltA/gltC}$ | This work |
| SynLinR$_{DBD}$GltC$_{LBD_H_LH}$ | LinR$_{DBD}$GltC$_{LBD_H_LH}$ | P$_{linE/linR}$ | This work |
| SynLinR$_{DBD_LH}$GltC$_{LBD_H}$ | LinR$_{DBD_LH}$GltC$_{LBD_H}$ | P$_{linE/linR}$ | This work |
| SynLinR$_{DBD_LH_H}$GltC$_{LBD}$ | LinR$_{DBD_LH_H}$GltC$_{LBD}$ | P$_{linE/linR}$ | This work |
| SynLinR$_{DBD}$AlsR$_{LBD_H_LH}$ | LinR$_{DBD}$AlsR$_{LBD_H_LH}$ | P$_{linE/linR}$ | This work |
| SynLinR$_{DBD_LH}$AlsR$_{LBD_H}$ | LinR$_{DBD_LH}$AlsR$_{LBD_H}$ | P$_{linE/linR}$ | This work |
| SynLinR$_{DBD_LH_H}$AlsR$_{LBD}$ | LinR$_{DBD_LH_H}$AlsR$_{LBD}$ | P$_{linE/linR}$ | This work |
| SynLinR$_{DBD}$BenM$_{LBD_H_LH}$ | LinR$_{DBD}$BenM$_{LBD_H_LH}$ | P$_{linE/linR}$ | This work |
| SynLinR$_{DBD_LH}$BenM$_{LBD_H}$ | LinR$_{DBD_LH}$BenM$_{LBD_H}$ | P$_{linE/linR}$ | This work |
| SynLinR$_{DBD_LH_H}$BenM$_{LBD}$ | LinR$_{DBD_LH_H}$BenM$_{LBD}$ | P$_{linE/linR}$ | This work |
| SynLinR$_{DBD}$ChiR$_{LBD_H_LH}$ | LinR$_{DBD}$ChiR$_{LBD_H_LH}$ | P$_{linE/linR}$ | This work |
| SynLinR$_{DBD_LH}$ChiR$_{LBD_H}$ | LinR$_{DBD_LH}$ChiR$_{LBD_H}$ | P$_{linE/linR}$ | This work |
| SynLinR$_{DBD_LH_H}$ChiR$_{LBD}$ | LinR$_{DBD_LH_H}$ChiR$_{LBD}$ | P$_{linE/linR}$ | This work |
| SynLinR$_{DBD}$MetR$_{LBD_H_LH}$ | LinR$_{DBD}$MetR$_{LBD_H_LH}$ | P$_{linE/linR}$ | This work |
| SynLinR$_{DBD_LH}$MetR$_{LBD_H}$ | LinR$_{DBD_LH}$MetR$_{LBD_H}$ | P$_{linE/linR}$ | This work |
| SynLinR$_{DBD_LH_H}$MetR$_{LBD}$ | LinR$_{DBD_LH_H}$MetR$_{LBD}$ | P$_{linE/linR}$ | This work |
| SynMetR$_{DBD}$AlsR$_{LBD_H_LH}$ | MetR$_{DBD}$AlsR$_{LBD_H_LH}$ | P$_{metE/metR}$ | This work |

| | | | |
|---|---|---|---|
| SynMetR$_{DBD_LH}$AlsR$_{LBD_H}$ | MetR$_{DBD_LH}$AlsR$_{LBD_H}$ | P$_{metE/metR}$ | This work |
| SynMetR$_{DBD_LH_H}$AlsR$_{LBD}$ | MetR$_{DBD_LH_H}$AlsR$_{LBD}$ | P$_{metE/metR}$ | This work |
| SynMetR$_{DBD}$BenM$_{LBD_H_LH}$ | MetR$_{DBD}$BenM$_{LBD_H_LH}$ | P$_{metE/metR}$ | This work |
| SynMetR$_{DBD_LH}$BenM$_{LBD_H}$ | MetR$_{DBD_LH}$BenM$_{LBD_H}$ | P$_{metE/metR}$ | This work |
| SynMetR$_{DBD_LH_H}$BenM$_{LBD}$ | MetR$_{DBD_LH_H}$BenM$_{LBD}$ | P$_{metE/metR}$ | This work |
| SynMetR$_{DBD}$ChiR$_{LBD_H_LH}$ | MetR$_{DBD}$ChiR$_{LBD_H_LH}$ | P$_{metE/metR}$ | This work |
| SynMetR$_{DBD_LH}$ChiR$_{LBD_H}$ | MetR$_{DBD_LH}$ChiR$_{LBD_H}$ | P$_{metE/metR}$ | This work |
| SynMetR$_{DBD_LH_H}$ChiR$_{LBD}$ | MetR$_{DBD_LH_H}$ChiR$_{LBD}$ | P$_{metE/metR}$ | This work |
| SynMetR$_{DBD}$GltC$_{LBD_H_LH}$ | MetR$_{DBD}$GltC$_{LBD_H_LH}$ | P$_{metE/metR}$ | This work |
| SynMetR$_{DBD_LH}$GltC$_{LBD_H}$ | MetR$_{DBD_LH}$GltC$_{LBD_H}$ | P$_{metE/metR}$ | This work |
| SynMetR$_{DBD_LH_H}$GltC$_{LBD}$ | MetR$_{DBD_LH_H}$GltC$_{LBD}$ | P$_{metE/metR}$ | This work |
| SynMetR$_{DBD}$LinR$_{LBD_H_LH}$ | MetR$_{DBD}$LinR$_{LBD_H_LH}$ | P$_{metE/metR}$ | This work |
| SynMetR$_{DBD_LH}$LinR$_{LBD_H}$ | MetR$_{DBD_LH}$LinR$_{LBD_H}$ | P$_{metE/metR}$ | This work |
| SynMetR$_{DBD_LH_H}$LinR$_{LBD}$ | MetR$_{DBD_LH_H}$LinR$_{LBD}$ | P$_{metE/metR}$ | This work |
| SynTsaR$_{DBD}$AlsR$_{LBD_H_LH}$ | TsaR$_{DBD}$AlsR$_{LBD_H_LH}$ | P$_{tsaM/tsaR}$ | This work |
| SynTsaR$_{DBD_LH}$AlsR$_{LBD_H}$ | TsaR$_{DBD_LH}$AlsR$_{LBD_H}$ | P$_{tsaM/tsaR}$ | This work |
| SynTsaR$_{DBD_LH_H}$AlsR$_{LBD}$ | TsaR$_{DBD_LH_H}$AlsR$_{LBD}$ | P$_{tsaM/tsaR}$ | This work |
| SynTsaR$_{DBD}$BenM$_{LBD_H_LH}$ | TsaR$_{DBD}$BenM$_{LBD_H_LH}$ | P$_{tsaM/tsaR}$ | This work |
| SynTsaR$_{DBD_LH}$BenM$_{LBD_H}$ | TsaR$_{DBD_LH}$BenM$_{LBD_H}$ | P$_{tsaM/tsaR}$ | This work |
| SynTsaR$_{DBD_LH_H}$BenM$_{LBD}$ | TsaR$_{DBD_LH_H}$BenM$_{LBD}$ | P$_{tsaM/tsaR}$ | This work |
| SynTsaR$_{DBD}$ChiR$_{LBD_H_LH}$ | TsaR$_{DBD}$ChiR$_{LBD_H_LH}$ | P$_{tsaM/tsaR}$ | This work |
| SynTsaR$_{DBD_LH}$ChiR$_{LBD_H}$ | TsaR$_{DBD_LH}$ChiR$_{LBD_H}$ | P$_{tsaM/tsaR}$ | This work |
| SynTsaR$_{DBD_LH_H}$ChiR$_{LBD}$ | TsaR$_{DBD_LH_H}$ChiR$_{LBD}$ | P$_{tsaM/tsaR}$ | This work |
| SynTsaR$_{DBD}$GltC$_{LBD_H_LH}$ | TsaR$_{DBD}$GltC$_{LBD_H_LH}$ | P$_{tsaM/tsaR}$ | This work |
| SynTsaR$_{DBD_LH}$GltC$_{LBD_H}$ | TsaR$_{DBD_LH}$GltC$_{LBD_H}$ | P$_{tsaM/tsaR}$ | This work |
| SynTsaR$_{DBD_LH_H}$GltC$_{LBD}$ | TsaR$_{DBD_LH_H}$GltC$_{LBD}$ | P$_{tsaM/tsaR}$ | This work |
| SynTsaR$_{DBD}$LinR$_{LBD_H_LH}$ | TsaR$_{DBD}$LinR$_{LBD_H_LH}$ | P$_{tsaM/tsaR}$ | This work |
| SynTsaR$_{DBD_LH}$LinR$_{LBD_H}$ | TsaR$_{DBD_LH}$LinR$_{LBD_H}$ | P$_{tsaM/tsaR}$ | This work |
| SynTsaR$_{DBD_LH_H}$LinR$_{LBD}$ | TsaR$_{DBD_LH_H}$LinR$_{LBD}$ | P$_{tsaM/tsaR}$ | This work |
| SynTsaR$_{DBD}$MetR$_{LBD_H_LH}$ | TsaR$_{DBD}$MetR$_{LBD_H_LH}$ | P$_{tsaM/tsaR}$ | This work |
| SynTsaR$_{DBD_LH}$MetR$_{LBD_H}$ | TsaR$_{DBD_LH}$MetR$_{LBD_H}$ | P$_{tsaM/tsaR}$ | This work |
| SynTsaR$_{DBD_LH_H}$MetR$_{LBD}$ | TsaR$_{DBD_LH_H}$MetR$_{LBD}$ | P$_{tsaM/tsaR}$ | This work |
| **Orthogonality analysis chimera** | | | |
| GltC$_{DBD_LH}$BenM$_{LBD_H}$ | GltC$_{DBD_LH}$BenM$_{LBD_H}$ | P$_{linE/linR}$ | This work |

**Supplementary Table S4 | Statistical analysis — functionality assay.** T/F values and p-values (Welch's t-tests for TF-expression; one-way ANOVA for ligand induction) for every strain in the functional screen; significant values in bold.

| | TF-promoter test | | Ligand induction test | |
|---|---|---|---|---|
| **Responsive biosensors** | | | | |
| **Plasmid** | ***T*** | ***p*-value** | ***F*** | ***p*-value** |
| SynAlsR | -89.56 | **$2.89*10^{-6}$** | 1473.92 | **$4.74*10^{-12}$** |
| NatAlsR | -39.30 | **$3.41*10^{-5}$** | 1444.87 | **$5.18*10^{-12}$** |
| SynBenM | 23.94 | **$4.55*10^{-4}$** | 34.00 | **$6.40*10^{-5}$** |
| NatBenM | 23.76 | **$7.01*10^{-5}$** | 8.13 | **$9.63*10^{-3}$** |
| SynGltC | -2.32 | 0.10 | 4.52 | **0.049** |
| NatGltC | -0.82 | 0.45 | 2.91 | 0.11 |
| SynLinR | -0.46 | 0.67 | 36.67 | **$4.70*10^{-5}$** |
| NatLinR | 0.94 | 0.41 | 10.81 | **$4.04*10^{-3}$** |
| SynMetR | 2.00 | 0.092 | 34.84 | **$5.80*10^{-5}$** |
| NatMetR | 4.88 | **$3.00*10^{-3}$** | 246.61 | **$1.38*10^{-8}$** |
| SynNahR | 22.83 | **$3.98*10^{-5}$** | 33.26 | **$6.96*10^{-5}$** |
| NatNahR | 23.90 | **$9.74*10^{-5}$** | 330.05 | **$3.80*10^{-9}$** |
| **Non-responsive biosensors** | | | | |
| **Plasmid** | ***T*** | ***p*-value** | ***T*** | ***p*-value** |
| SynCitR (Na-citrate) | 3.97 | **0.028** | -0.19 | 0.86 |
| NatCitR (Na-citrate) | 15.08 | **$2.14*10^{-5}$** | 1.87 | 0.11 |
| SynCitR (oxaloac) | Same values as for Na-citrate | | -0.85 | 0.43 |
| NatCitR (oxaloac) | Same values as for Na-citrate | | 1.35 | 0.23 |
| **Plasmid** | ***T*** | ***p*-value** | ***F*** | ***p*-value** |
| SynTsaR | 1.17 | **$7.66*10^{-3}$** | 2.90 | 0.11 |
| NatTsaR | 4.85 | 0.30 | 0.32 | 0.73 |
| SynCysB | -42.56 | **$5.41*10^{-4}$** | / | |
| NatCysB | -5.44 | **$3.98*10^{-3}$** | / | |
| **Non-functional biosensors** | | | | |
| SynAllS | 1.31 | 0.25 | Interference of ligand | |
| SynAmpR | 0.65 | 0.54 | 2.88 | 0.11 |
| NatAmpR | -0.91 | 0.42 | 0.78 | 0.49 |
| SynLysG | 2.16 | 0.11 | 0.094 | 0.91 |
| NatLysG | 2.53 | 0.085 | 0.62 | 0.56 |
| SynNagR | 1.21 | 0.30 | 0.86 | 0.45 |
| NatNagR | 1.74 | 0.31 | 0.98 | 0.40 |
| SynOccR | 1.64 | 0.20 | 0.90 | 0.44 |
| NatOccR | -0.15 | 0.88 | 0.18 | 0.84 |
| SynPqsR | -1.33 | 0.25 | 0.50 | 0.62 |
| SynTtuA | 0.21 | 0.84 | 2.69 | 0.12 |
| NatTtuA | 0.60 | 0.59 | 1.32 | 0.31 |

**Supplementary Table S5 | Statistical analysis — orthogonality assay.** Welch's t-test statistics (TF-expression and ligand-addition tests) for all orthogonality combinations; significant values in bold.

| | TF-promoter test | | Induced-uninduced test | |
|---|---|---|---|---|
| **Responsive regulators** | | | | |
| **Plasmid** | ***T*** | ***p*-value** | ***T*** | ***p*-value** |
| PAlsRSynBenM | 5.52 | **$4.58*10^{-3}$** | -4.598 | **$3.75*10^{-3}$** |
| PAlsRSynChiR | 1.23 | 0.27 | / | |
| PAlsRSynCitR | 2.47 | **0.048** | / | |
| PAlsRSynCysB | -8.57 | **$2.22*10^{-3}$** | / | |
| PAlsRSynFdeR | 1.92 | 0.11622 | -0.51 | 0.6314 |
| PAlsRSynGltC | -14.71 | **$4.59*10^{-4}$** | 3.16 | **0.022** |
| PAlsRSynLinR | -0.87 | 0.41574 | -4.87 | **$5.45*10^{-3}$** |
| PAlsRSynMetR | 3.61 | **0.011** | 0.94 | 0.38 |
| PAlsRSynNahR | 0.15 | 0.89 | -2.54 | **0.044** |
| PAlsRSynTsaR | 46.04 | **$2.24*10^{-5}$** | / | |
| PBenMSynAlsR | -0.45 | 0.68 | -1.46 | 0.20 |
| PBenMSynChiR | 0.19 | 0.86 | / | |
| PBenMSynCitR | -1.23 | 0.31 | / | |
| PBenMSynCysB | -1.74 | 0.18 | / | |
| PBenMSynFdeR | -3.37 | **0.041** | -1.44 | 0.22 |
| PBenMSynGltC | 5.11 | **0.013** | -1.20 | 0.28 |
| PBenMSynLinR | -3.33 | **0.031** | -0.68 | 0.52 |
| PBenMSynMetR | 2.63 | 0.059 | -0.077 | 0.94 |
| PBenMSynNahR | -2.82 | **0.036** | -2.18 | 0.079 |
| PBenMSynTsaR | -9.44 | **$1.79*10^{-3}$** | / | |
| PChiRSynAlsR | 3.36 | **0.018** | -11.10 | **$1.56*10^{-3}$** |
| PChiRSynBenM | 9.73 | **$7.93*10^{-4}$** | -1.12 | 0.31 |
| PChiRSynCitR | -1.13 | 0.31 | / | |
| PChiRSynCysB | 1.59 | 0.18 | / | |
| PChiRSynFdeR | 14.59 | **$1.07*10^{-4}$** | -0.59 | 0.58 |
| PChiRSynGltC | 4.18 | **0.015** | 3.93 | **$7.74*10^{-3}$** |
| PChiRSynLinR | -0.35 | 0.74 | -17.61 | **$3.76*10^{-6}$** |
| PChiRSynMetR | 9.45 | **$1.69*10^{-4}$** | -0.99 | 0.36 |
| PChiRSynNahR | -0.50 | 0.65 | -2.03 | 0.12 |
| PChiRSynTsaR | 0.94 | 0.41 | / | |
| PCitRSynAlsR | 2.47 | 0.064 | 3.62 | **0.011** |
| PCitRSynBenM | 30.27 | **$1.44*10^{-7}$** | 5.20 | **$3.62*10^{-3}$** |
| PCitRSynChiR | 4.08 | **0.026** | / | |
| PCitRSynCysB | 31.37 | **$9.21*10^{-7}$** | / | |
| PCitRSynFdeR | -1.75 | 0.14 | -0.63 | 0.59 |
| PCitRSynGltC | 6.31 | **$1.30*10^{-3}$** | 10.58 | **$5.49*10^{-5}$** |
| PCitRSynLinR | -1.22 | 0.27 | 13.74 | **$1.84*10^{-4}$** |
| PCitRSynMetR | 7.55 | **$2.83*10^{-4}$** | 1.05 | 0.34 |
| PCitRSynNahR | 1.63 | 1.54 | -0.24 | 0.82 |
| PCitRSynTsaR | -5.41 | **$8.25*10^{-3}$** | / | |
| PCysBSynAlsR | -3.67 | **0.029** | 2.24 | 0.069 |
| PCysBSynBenM | -3.78 | **0.024** | 1.48 | 0.20 |

| | | | | |
|---|---|---|---|---|
| PCysBSynChiR | -2.24 | 0.10 | / | |
| PCysBSynCitR | -2.64 | 0.077 | / | |
| PCysBSynFdeR | -124.18 | **5.02*10$^{-9}$** | 0.097 | 0.93 |
| PCysBSynGltC | -2.09 | 0.11 | 0.69 | 0.51 |
| PCysBSynLinR | -0.43 | 0.69 | -0.96 | 0.39 |
| PCysBSynMetR | -0.11 | 0.92 | -24.91 | **5.22*10$^{-5}$** |
| PCysBSynNahR | 0.091 | 0.93 | -1.04 | 0.34 |
| PCysBSynTsaR | -2.20 | 0.076 | / | |
| PFdeRSynAlsR | 0.42 | 0.69 | 0.42 | 0.69 |
| PFdeRSynBenM | 1.72 | 0.14 | -3.30 | **0.037** |
| PFdeRSynChiR | -0.89 | 0.42 | / | |
| PFdeRSynCitR | 1.06 | 0.34 | / | |
| PFdeRSynCysB | -0.88 | 0.41 | / | |
| PFdeRSynGltC | 0.50 | 0.64 | 3.83 | **8.78*10$^{-3}$** |
| PFdeRSynMetR | 1.03 | 0.36 | -4.28*10$^{-3}$ | 0.99 |
| PFdeRSynNahR | -1.16 | 0.30 | -2.33 | 0.060 |
| PFdeRSynTsaR | -0.44 | 0.68 | / | |
| PGltCSynAlsR | 0.15 | 0.88 | 2.78 | **0.033** |
| PGltCSynBenM | 6.30 | **8.30*10$^{-4}$** | -4.62 | **4.88*10$^{-3}$** |
| PGltCSynChiR | -0.017 | 0.99 | / | |
| PGltCSynCitR | 9.56 | **1.65*10$^{-3}$** | / | |
| PGltCSynCysB | -2.67 | 0.061 | / | |
| PGltCSynFdeR | -5.36 | **7.18*10$^{-3}$** | -1.02 | 0.36 |
| PGltCSynLinR | -0.83 | 0.45 | 0.19 | 0.86 |
| PGltCSynMetR | 5.14 | **2.53*10$^{-3}$** | 0.74 | 0.49 |
| PGltCSynNahR | -0.90 | 0.40 | -3.25 | **0.023** |
| PGltCSynTsaR | -6.37 | **4.00*10$^{-3}$** | / | |
| PLinRSynAlsR | 0.043 | 0.97 | 0.57 | 0.60 |
| PLinRSynBenM | 1.99 | 0.12 | -1.13 | 0.33 |
| PLinRSynChiR | 0.84 | 0.44 | / | |
| PLinRSynCitR | -1.17 | 0.33 | / | |
| PLinRSynCysB | -3.95 | **0.013** | / | |
| PLinRSynFdeR | -0.82 | 0.45 | -2.23 | 0.10 |
| PLinRSynGltC | -1.80 | 0.13 | 1.20 | 0.29 |
| PLinRSynMetR | 0.10 | 0.92 | 1.06 | 0.33 |
| PLinRSynNahR | -0.039 | 0.97 | -4.28 | **7.83*10$^{-3}$** |
| PLinRSynTsaR | 1.59 | 0.17 | / | |
| PMetRSynAlsR | 0.51 | 0.65 | 0.33 | 0.75 |
| PMetRSynBenM | 4.92 | **0.011** | -3.76 | **0.012** |
| PMetRSynChiR | 2.11 | 0.097 | / | |
| PMetRSynCysB | 0.043 | 0.97 | / | |
| PMetRSynFdeR | 0.76 | 0.49 | -1.65 | 0.15 |
| PMetRSynGltC | 0.91 | 0.41 | -7.28 | **1.01*10$^{-3}$** |
| PMetRSynLinR | -1.17 | 0.32 | -2.24 | 0.072 |
| PMetRSynNahR | 0.39 | 0.71 | -3.34 | **0.023** |
| PMetRSynTsaR | -8.06 | **2.22*10$^{-3}$** | / | |
| PNahRSynAlsR | 4.72 | **7.36*10$^{-3}$** | 2.31 | 0.065 |
| PNahRSynBenM | 12.72 | **1.45*10$^{-5}$** | 6.69 | **1.55*10$^{-3}$** |
| PNahRSynChiR | -2.84 | **0.030** | / | |

| | | | | |
|---|---|---|---|---|
| PNahRSynCitR | 1.62 | 0.20 | / | |
| PNahRSynCysB | 8.00 | **$3.44*10^{-3}$** | / | |
| PNahRSynFdeR | -0.36 | 0.73 | 1.57 | 0.19 |
| PNahRSynGltC | 2.08 | 0.084 | 5.93 | **$1.05*10^{-3}$** |
| PNahRSynLinR | -0.86 | 0.45 | -0.20 | 0.85 |
| PNahRSynMetR | -1.71 | 0.15 | -0.87 | 0.42 |
| PNahRSynTsaR | -10.19 | **$5.58*10^{-4}$** | / | |
| PTsaRSynAlsR | 10.82 | **$4.09*10^{-5}$** | 3.04 | **0.040** |
| PTsaRSynBenM | 7.69 | **$4.24*10^{-4}$** | -0.34 | 0.75 |
| PTsaRSynChiR | 12.67 | **$1.63*10^{-5}$** | / | |
| PTsaRSynCysB | -10.38 | **$1.51*10^{-3}$** | / | |
| PTsaRSynFdeR | 8.32 | **$2.16*10^{-4}$** | -0.24 | 0.82 |
| PTsaRSynGltC | 2.89 | **0.036** | 7.40 | **$3.96*10^{-4}$** |
| PTsaRSynLinR | 4.24 | **$6.21*10^{-3}$** | -13.72 | **$1.27*10^{-5}$** |
| PTsaRSynMetR | 4.91 | **$3.46*10^{-3}$** | 0.010 | 0.99 |
| PTsaRSynNahR | 2.37 | 0.056 | -1.24 | 0.26 |

**Supplementary Table S6 | Statistical analysis — domain-swapping analysis.** Welch's t-test statistics for chimera-expression and ligand-induction effects across the three domain-swap designs; significant values in bold.

| | Chimer-promoter test | | Induced-uninduced test | |
|---|---|---|---|---|
| **Plasmid** | ***T*** | ***p*-value** | ***T*** | ***p*-value** |
| **DBD design** | | | | |
| SynAlsR$_{DBD}$BenM$_{LBD_H_LH}$ | -0.043 | 0.97 | -0.41 | 0.70 |
| SynAlsR$_{DBD}$ChiR$_{LBD_H_LH}$ | Circuit non-viable | | | |
| SynAlsR$_{DBD}$GltC$_{LBD_H_LH}$ | -40.29 | **1.52*10$^{-14}$** | -10.14 | **1.33*10$^{-4}$** |
| SynAlsR$_{DBD}$LinR$_{LBD_H_LH}$ | 5.60 | **7.29*10$^{-4}$** | -5.550 | **1.45*10$^{-3}$** |
| SynAlsR$_{DBD}$MetR$_{LBD_H_LH}$ | 0.28 | 0.79 | 0.14 | 0.89 |
| SynBenM$_{DBD}$AlsR$_{LBD_H_LH}$ | 20.64 | **4.80*10$^{-7}$** | 0.61 | 0.55 |
| SynBenM$_{DBD}$ChiR$_{LBD_H_LH}$ | -20.90 | **3.66*10$^{-6}$** | No induction tested | |
| SynBenM$_{DBD}$GltC$_{LBD_H_LH}$ | Circuit non-viable | | | |
| SynBenM$_{DBD}$LinR$_{LBD_H_LH}$ | 21.82 | **6.41*10$^{-6}$** | -5.50 | **0.023** |
| SynBenM$_{DBD}$MetR$_{LBD_H_LH}$ | 4.54 | **4.93*10$^{-3}$** | -0.64 | 0.55 |
| SynChiR$_{DBD}$AlsR$_{LBD_H_LH}$ | -0.015 | 0.99 | 2.20 | 0.70 |
| SynChiR$_{DBD}$BenM$_{LBD_H_LH}$ | 6.67 | **1.41*10$^{-3}$** | 1.41 | 0.21 |
| SynChiR$_{DBD}$GltC$_{LBD_H_LH}$ | 3.97 | **0.012** | 0.55 | 0.61 |
| SynChiR$_{DBD}$LinR$_{LBD_H_LH}$ | -1.38 | 0.22 | -13.00 | **1.29*10$^{-5}$** |
| SynChiR$_{DBD}$MetR$_{LBD_H_LH}$ | 4.58 | **0.015** | 0.59 | 0.59 |
| SynGltC$_{DBD}$AlsR$_{LBD_H_LH}$ | 1.85 | 0.094 | -14.78 | **6.08*10$^{-6}$** |
| SynGltC$_{DBD}$BenM$_{LBD_H_LH}$ | -1.62 | 0.14 | 0.78 | 0.47 |
| SynGltC$_{DBD}$ChiR$_{LBD_H_LH}$ | -10.45 | **2.11*10$^{-6}$** | No induction tested | |
| SynGltC$_{DBD}$LinR$_{LBD_H_LH}$ | -0.70 | 0.50 | 1.18 | 0.29 |
| SynGltC$_{DBD}$MetR$_{LBD_H_LH}$ | 7.25 | **8.37*10$^{-5}$** | 3.54 | **0.025** |
| SynLinR$_{DBD}$AlsR$_{LBD_H_LH}$ | -1.72 | 0.13 | 1.42 | 0.19 |
| SynLinR$_{DBD}$BenM$_{LBD_H_LH}$ | 1.60 | 0.15 | -0.90 | 0.43 |
| SynLinR$_{DBD}$ChiR$_{LBD_H_LH}$ | -0.15 | 0.89 | No induction tested | |
| SynLinR$_{DBD}$GltC$_{LBD_H_LH}$ | 0.048 | 0.96 | -0.017 | 0.99 |
| SynLinR$_{DBD}$MetR$_{LBD_H_LH}$ | -2.87 | **0.30** | -1.71 | 0.14 |
| SynMetR$_{DBD}$AlsR$_{LBD_H_LH}$ | 1.50 | 0.22 | -0.40 | 0.70 |
| SynMetR$_{DBD}$BenM$_{LBD_H_LH}$ | 6.04 | **2.50*10$^{-3}$** | 0.18 | 0.86 |
| SynMetR$_{DBD}$ChiR$_{LBD_H_LH}$ | -57.78 | **3.07*10$^{-9}$** | No induction tested | |
| SynMetR$_{DBD}$GltC$_{LBD_H_LH}$ | 1.24 | 0.27 | -3.97 | **7.94*10$^{-3}$** |
| SynMetR$_{DBD}$LinR$_{LBD_H_LH}$ | 8.31 | **2.58*10$^{-4}$** | 0.26 | 0.81 |
| SynTsaR$_{DBD}$AlsR$_{LBD_H_LH}$ | 0.40 | 0.70 | 1.13 | 0.28 |
| SynTsaR$_{DBD}$BenM$_{LBD_H_LH}$ | 11.47 | **2.05*10$^{-4}$** | -1.30 | 0.28 |
| SynTsaR$_{DBD}$ChiR$_{LBD_H_LH}$ | 3.08 | **0.029** | No induction tested | |
| SynTsaR$_{DBD}$GltC$_{LBD_H_LH}$ | 11.90 | **6.08*10$^{-4}$** | 0.12 | 0.91 |
| SynTsaR$_{DBD}$LinR$_{LBD_H_LH}$ | 6.08 | **1.62*10$^{-3}$** | -11.35 | **1.01*10$^{-3}$** |
| SynTsaR$_{DBD}$MetR$_{LBD_H_LH}$ | 0.94 | 0.39 | 0.19 | 0.86 |
| **DBD_LH design** | | | | |
| SynAlsR$_{DBD_LH}$BenM$_{LBD_H}$ | 10.84 | **1.37*10$^{-8}$** | -3.21 | **0.024** |
| SynAlsR$_{DBD_LH}$ChiR$_{LBD_H}$ | -15.32 | **4.21*10$^{-10}$** | No induction tested | |
| SynAlsR$_{DBD_LH}$GltC$_{LBD_H}$ | -56.57 | **6.13*10$^{-9}$** | 2.54 | 0.075 |
| SynAlsR$_{DBD_LH}$LinR$_{LBD_H}$ | 3.79 | **6.44*10$^{-3}$** | -5.29 | **2.74*10$^{-3}$** |
| SynAlsR$_{DBD_LH}$MetR$_{LBD_H}$ | 0.10 | 0.92 | 0.23 | 0.83 |

| | | | | |
|---|---|---|---|---|
| SynBenM$_{DBD_LH}$AlsR$_{LBD_H}$ | 22.67 | **3.16*10$^{-8}$** | 2.21 | **0.046** |
| SynBenM$_{DBD_LH}$ChiR$_{LBD_H}$ | -4.03 | **0.028** | No induction tested | |
| SynBenM$_{DBD_LH}$GltC$_{LBD_H}$ | -9.37 | **0.00014** | -8.07 | **0.00020** |
| SynBenM$_{DBD_LH}$LinR$_{LBD_H}$ | 17.06 | **1.67*10$^{-5}$** | -5.47 | **1.66*10$^{-3}$** |
| SynBenM$_{DBD_LH}$MetR$_{LBD_H}$ | 5.45 | **2.47*10$^{-3}$** | -2.52 | **0.049** |
| SynChiR$_{DBD_LH}$AlsR$_{LBD_H}$ | 2.82 | **0.035** | -0.54 | 0.61 |
| SynChiR$_{DBD_LH}$BenM$_{LBD_H}$ | 7.56 | **1.77*10$^{-3}$** | -2.16 | 0.087 |
| SynChiR$_{DBD_LH}$GltC$_{LBD_H}$ | 2.00 | 0.094 | 2.41 | 0.059 |
| SynChiR$_{DBD_LH}$LinR$_{LBD_H}$ | -0.015 | 0.99 | -13.48 | **1.04*10$^{-5}$** |
| SynChiR$_{DBD_LH}$MetR$_{LBD_H}$ | 2.04 | 0.088 | 0.50 | 0.63 |
| SynGltC$_{DBD_LH}$AlsR$_{LBD_H}$ | -3.39 | **6.94*10$^{-3}$** | -1.56 | **0.021** |
| SynGltC$_{DBD_LH}$BenM$_{LBD_H}$ | -8.08 | **4.33*10$^{-5}$** | -4.15 | **0.013** |
| SynGltC$_{DBD_LH}$ChiR$_{LBD_H}$ | -9.02 | **7.51*10$^{-6}$** | No induction tested | |
| SynGltC$_{DBD_LH}$LinR$_{LBD_H}$ | -6.46 | **1.28*10$^{-4}$** | -0.89 | 0.41 |
| SynGltC$_{DBD_LH}$MetR$_{LBD_H}$ | 7.31 | **5.83*10$^{-4}$** | 1.09 | 0.32 |
| SynLinR$_{DBD_LH}$AlsR$_{LBD_H}$ | -1.26 | 0.25 | 0.61 | 0.56 |
| SynLinR$_{DBD_LH}$BenM$_{LBD_H}$ | -0.79 | 0.47 | -0.60 | 0.57 |
| SynLinR$_{DBD_LH}$ChiR$_{LBD_H}$ | Circuit non-viable | | | |
| SynLinR$_{DBD_LH}$GltC$_{LBD_H}$ | 1.02 | 0.33 | -0.22 | 0.84 |
| SynLinR$_{DBD_LH}$MetR$_{LBD_H}$ | -2.94 | **0.033** | 0.12 | 0.91 |
| SynMetR$_{DBD_LH}$AlsR$_{LBD_H}$ | 13.08 | **5.69*10$^{-5}$** | 1.01 | 0.36 |
| SynMetR$_{DBD_LH}$BenM$_{LBD_H}$ | 37.37 | **4.34*10$^{-8}$** | -11.59 | **7.79*10$^{-4}$** |
| SynMetR$_{DBD_LH}$ChiR$_{LBD_H}$ | Circuit non-viable | | | |
| SynMetR$_{DBD_LH}$GltC$_{LBD_H}$ | 4.14 | 0.023 | -0.47 | 0.66 |
| SynMetR$_{DBD_LH}$LinR$_{LBD_H}$ | 13.67 | **2.29*10$^{-5}$** | -12.56 | **1.59*10$^{-5}$** |
| SynTsaR$_{DBD_LH}$AlsR$_{LBD_H}$ | 1.06 | 0.32 | -1.39 | 0.21 |
| SynTsaR$_{DBD_LH}$BenM$_{LBD_H}$ | 15.32 | **2.90*10$^{-5}$** | 0.62 | 0.57 |
| SynTsaR$_{DBD_LH}$ChiR$_{LBD_H}$ | 6.69 | **6.09*10$^{-4}$** | No induction tested | |
| SynTsaR$_{DBD_LH}$GltC$_{LBD_H}$ | 3.55 | **0.012** | 7.73 | **3.24*10$^{-3}$** |
| SynTsaR$_{DBD_LH}$LinR$_{LBD_H}$ | 6.03 | **3.78*10$^{-3}$** | -23.28 | **1.05*10$^{-6}$** |
| SynTsaR$_{DBD_LH}$MetR$_{LBD_H}$ | -0.029 | 0.98 | 0.54 | 0.61 |
| **DBD_LH_H design** | | | | |
| SynAlsR$_{DBD_LH_H}$BenM$_{LBD}$ | -2.11 | 0.096 | -0.42 | 0.69 |
| SynAlsR$_{DBD_LH_H}$ChiR$_{LBD}$ | -0.067 | 0.95 | No induction tested | |
| SynAlsR$_{DBD_LH_H}$GltC$_{LBD}$ | -18.23 | **1.67*10$^{-4}$** | -1.10 | 0.31 |
| SynAlsR$_{DBD_LH_H}$LinR$_{LBD}$ | -4.05 | **0.027** | 0.087 | 0.93 |
| SynAlsR$_{DBD_LH_H}$MetR$_{LBD}$ | 0.45 | 0.68 | 0.047 | 0.96 |
| SynBenM$_{DBD_LH_H}$AlsR$_{LBD}$ | -0.95 | 0.36 | 2.90 | **0.012** |
| SynBenM$_{DBD_LH_H}$ChiR$_{LBD}$ | -0.76 | 0.50 | No induction tested | |
| SynBenM$_{DBD_LH_H}$GltC$_{LBD}$ | Circuit non-viable | | | |
| SynBenM$_{DBD_LH_H}$LinR$_{LBD}$ | 42.87 | **8.51*10$^{-7}$** | -17.67 | **6.73*10$^{-6}$** |
| SynBenM$_{DBD_LH_H}$MetR$_{LBD}$ | 2.09 | 0.086 | -0.54 | 0.62 |
| SynChiR$_{DBD_LH_H}$AlsR$_{LBD}$ | 2.69 | **0.036** | 0.85 | 0.43 |
| SynChiR$_{DBD_LH_H}$BenM$_{LBD}$ | 7.74 | **3.36*10$^{-3}$** | 2.89 | 0.49 |
| SynChiR$_{DBD_LH_H}$GltC$_{LBD}$ | 3.64 | **0.012** | 2.80 | **0.031** |
| SynChiR$_{DBD_LH_H}$LinR$_{LBD}$ | -0.65 | 0.54 | -10.23 | **4.94*10$^{-4}$** |
| SynChiR$_{DBD_LH_H}$MetR$_{LBD}$ | 3.80 | **0.013** | -0.71 | 0.51 |
| SynGltC$_{DBD_LH_H}$AlsR$_{LBD}$ | -0.070 | 0.95 | 2.34 | 0.075 |
| SynGltC$_{DBD_LH_H}$BenM$_{LBD}$ | -4.41 | **0.013** | 2.86 | **0.030** |

| | | | | |
|---|---|---|---|---|
| SynGltC$_{DBD_LH_H}$ChiR$_{LBD}$ | 0.49 | 0.64 | No induction tested | |
| SynGltC$_{DBD_LH_H}$LinR$_{LBD}$ | -2.00 | 0.074 | 0.61 | 0.57 |
| SynGltC$_{DBD_LH_H}$MetR$_{LBD}$ | 7.93 | **$2.26*10^{-5}$** | 0.95 | 0.38 |
| SynLinR$_{DBD_LH_H}$AlsR$_{LBD}$ | -0.88 | 0.40 | -0.53 | 0.61 |
| SynLinR$_{DBD_LH_H}$BenM$_{LBD}$ | -1.45 | 0.24 | 1.50 | 0.22 |
| SynLinR$_{DBD_LH_H}$ChiR$_{LBD}$ | 1.77 | 0.12 | No induction tested | |
| SynLinR$_{DBD_LH_H}$GltC$_{LBD}$ | 0.97 | 0.35 | 1.00 | 0.39 |
| SynLinR$_{DBD_LH_H}$MetR$_{LBD}$ | -4.77 | **$8.48*10^{-3}$** | 3.09 | **0.031** |
| SynMetR$_{DBD_LH_H}$AlsR$_{LBD}$ | 6.75 | **$2.87*10^{-3}$** | -2.78 | **0.049** |
| SynMetR$_{DBD_LH_H}$BenM$_{LBD}$ | Circuit non-viable | | | |
| SynMetR$_{DBD_LH_H}$ChiR$_{LBD}$ | Circuit non-viable | | | |
| SynMetR$_{DBD_LH_H}$GltC$_{LBD}$ | Circuit non-viable | | | |
| SynMetR$_{DBD_LH_H}$LinR$_{LBD}$ | 6.45 | **$7.35*10^{-4}$** | -4.24 | **$5.79*10^{-3}$** |
| SynTsaR$_{DBD_LH_H}$AlsR$_{LBD}$ | 3.20 | **0.011** | -1.02 | 0.33 |
| SynTsaR$_{DBD_LH_H}$BenM$_{LBD}$ | 3.25 | **0.020** | 0.72 | 0.50 |
| SynTsaR$_{DBD_LH_H}$ChiR$_{LBD}$ | 5.96 | **$1.65*10^{-3}$** | No induction tested | |
| SynTsaR$_{DBD_LH_H}$GltC$_{LBD}$ | 16.71 | **$3.29*10^{-5}$** | 1.96 | 0.14 |
| SynTsaR$_{DBD_LH_H}$LinR$_{LBD}$ | 2.47 | **0.076** | -7.37 | **$2.63*10^{-3}$** |
| SynTsaR$_{DBD_LH_H}$MetR$_{LBD}$ | 1.95 | 0.10 | 0.80 | 0.45 |
